# Assembly of Macromolecular Complexes in the Whole-Cell Model of a Minimal Cell

**DOI:** 10.1101/2025.07.02.662439

**Authors:** Enguang Fu, Zane R. Thornburg, Troy A. Brier, Rong Wei, Bo Yuan, Benjamin R. Gilbert, Shulei Wang, Zaida Luthey-Schulten

## Abstract

Macromolecular complexes in the genetically minimized bacterium, JCVI-syn3A, support gene expression (RNA polymerase, ribosome, degradosome), metabolism (ABC transporters, ATP synthase) and chromosome dynamics. In this work, we further incorporate the assembly of 21 unique macromolecular complexes into the existing whole-cell kinetic model of Syn3A. The synthesis and translocation of protein subunits in membrane complexes occur through distinct pathways. A range of 2D association rates for membrane complexes were considered to guarantee a high yield of assembly given the existing time scales of gene expression. By alleviating the undesired kinetically trapped intermediates in ATP synthase assembly, the efficiency was improved. The assembly of RNA polymerase, ribosome, and degradosome influence the speed and efficiency of protein synthesis. Collectively, this model predicted time-dependent cellular behaviors consistent with experiments. A machine learning analysis of the time-dependent metabolomics and metabolic fluxes highlighted the effect of introducing complex assembly into our whole-cell model.

## Introduction

The minimal cell, JCVI-syn3A, is a bacterium with a chemically synthesized genome derived from the parental organism *Mycoplasma mycoides subsp. capri* strain GM12.^1,2^ Syn3A exhibits a doubling time of less than two hours^2–4^ and possesses the smallest genome known among autonomously replicating organisms. This minimal genome comprises 543 kbp, encoding 452 protein-coding genes, 6 rRNA-coding (two sets of rRNA operons), and 29 tRNA-coding genes. Of the protein-coding genes, roughly 90 genes have unknown functions, the lowest proportion among model organisms such as *Mycoplasma pneumoniae* and *Escherichia coli*.^2^ During the reduction over half of the *M. mycoides* genome was removed including the majority of regulation factors. ^1^

In 2022,^5^ we integrated kinetic parameters, omics data, ^2^ and spatial data (cell size and ribosome distribution) from cryo-electron tomography (cryo-ET)^6^ to construct a comprehensive whole-cell model (WCM) of Syn3A. The genetic information processes of DNA replication, transcription, translation, and mRNA degradation ^7^ were treated stochastically using chemical master equations (CMEs), while the essential metabolism^2^ with thermodynamically consistent kinetic parameters^5^ was deterministically simulated using ordinary differential equations (ODEs). To simulate cooperative dynamics between two subsystems, a hybrid CME-ODE algorithm was used in which the simulations were discretized in time, and information between them occurred at a prescribed communication time.^5,8^ This approach enabled the calculation of time-dependent abundances of all genes, RNAs, and proteins, and metabolites. We simulated trajectories throughout the cell cycle and investigated the balance of energy cost and metabolic resources.^5^ This early model used probabilistic factors obtained from short 3D simulations using the static morphology and ribosome distributions obtained from the cyro-electron tomograms. It also used abundances of specified subunits from the proteomics data to represent the complexes and did not consider formation of intermediates which we now consider.

In all organisms including Syn3A, key macromolecular complexes mediate essential cellular processes. In genetic information processing, RNA polymerase (RNAP) and ribosomes decode genetic information into mRNAs and functional proteins via transcription and translation, respectively. Degradosomes, a loose complex of endoribonuclease, exoribonuclease, and a few glycolytic enzymes compete with ribosomes for the binding and processing of mRNA. The balance of this competition serves as the primary strategy controlling protein generation within the minimal cell. Several essential complexes interact with chromosomes and facilitate chromosome partitioning during cell division. ^9^ The essential metabolism of Syn3A depends on the activated transport of (deoxy)nucleosides, amino acids, vitamins and fatty acids into the cell by membrane complexes with multiple subunits. The Sec translocon is a vital membrane-embedded machinery for translocation of transmembrane proteins and secretory proteins.^10^ ATP synthase has to assemble eight kinds of subunits in varying stoichiometry in order to function properly. In nucleotide metabolism, ribonucleosidediphosphate reducatese (RNDR) converts ribonucleoside diphosphates to deoxyribonucleoside diphosphates to help balance the conversion from NTP to dNTP when uptake of (deoxy)nucleosides is insufficient.

The complexity of *in vivo* macromolecular complex assembly arises from the following aspects. First, the complicated assembly landscape consists of the composition of the macromolecular complex, the pathways that depend on the variation in affinity between the protein subunits, and rates for each bimolecular binding step. Second, the different dimensionalities including 3D cytoplasm, 2D cell membrane, and the 1D chromosome further complicate the assembly. The *in vivo* translocation of membrane-related proteins, that is, the placement of proteins in the membrane, occurs through an orchestrated network that includes actively translating ribosomes, targeting factors, and membrane-embedded insertases and translocons. ^11,12^ Lastly, macromolecular assembly must be kinetically consistent with the continuing subunit synthesis from gene expression along the cell cycle. Slow assembly leads to the accumulation of unassembled subunit precursors that possibly form aggregates and hinder cellular activities.^13^

Experimental investigation of assembly pathways typically begins with the identification of assembly intermediates using imaging, ^14,15^ chromatography,^16^ and mass spectrometry,^17^ followed by the elucidation of the preferred pathways through *in vitro* reconstruction^18,19^ and time-delayed assembly experiments.^20^ Quantification of the bimolecular binding rates along the assembly pathways is difficult because of the diverse protein-protein interactions and numerous assembly intermediates. One example is the classic investigation on ribosome biogenesis. The assembly of bacterial ribosomes (ribosome biogenesis) is crucial for protein synthesis and is highly relevant to cell growth. The bacterial ribosome comprises a small subunit (30S SSU) and a large subunit (50S LSU), each of which is a ribonucleoprotein complex that assembles independently. The assembly pathways of both subunits are hierarchical proceeding from the 5’ to the 3’ as indicated by the pioneering exploration of the SSU by Nomura *et al*.^21^ and in the LSU by Nierhaus *et al*.^22^ and Williamson *et al*.^15^ For SSU, 16S rRNA is first bound by primary binding proteins, and then secondary and tertiary ribosomal proteins bind by interacting with rRNA and bound proteins, called thermodynamic dependencies. Similar hierarchy was also found for LSU but with less clear separation in the order in more recent studies.^19,23^ In 2015, we simulated the SSU assembly in *E. coli* with coupled transcription, translation, and mRNA degradation using stochastic reaction-diffusion master equations with rigorous parameterization of ribosome subunits binding rates by fit to the pulse-chase data of *in vitro* SSU reconstruction from the Williamson laboratory.^24^ The SSU assembly pathways comprising 145 intermediates and multiple parallel pathways are shown in Figure 1(a). Later, the model was extended to include a simplified treatment of cell growth, DNA replication, and cell division.^25^ Here, in the context of a WCM, we investigate the effects on ribosome biogenesis upon reducing the parallel assembly pathway of SSU to a linear sequential one with only 19 intermediates that marked by green arrows.

**Figure 1.**
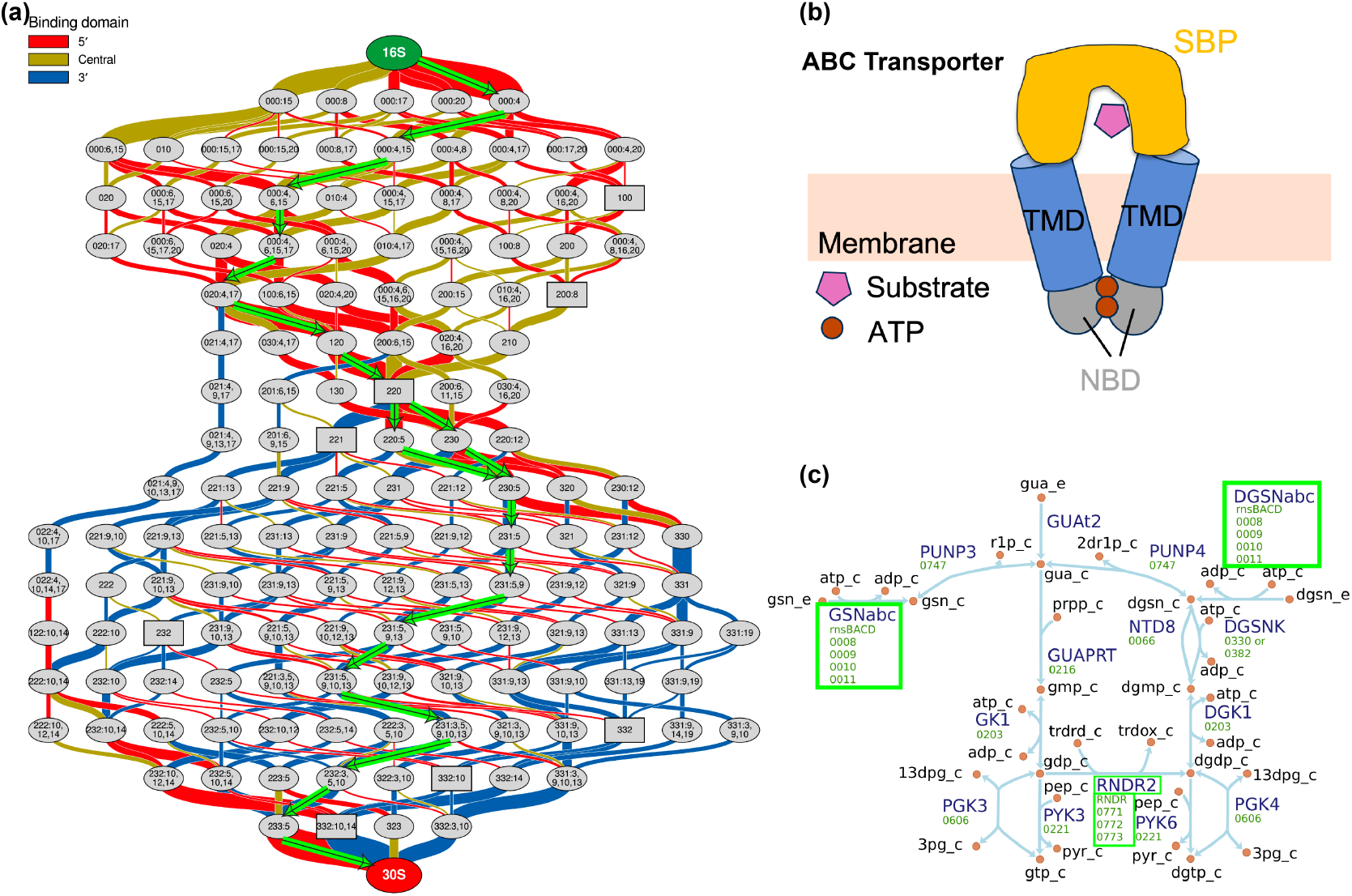
Ribosome 30S small subunit (SSU) assembly pathway, domain composition of ABC transporters, and nucleotide metabolism in Syn3A. (a) Hierarchical and parallel assembly pathway of SSU with 145 assembly intermediates,^26^ where the thickness of edges represents the fluxes. The marked green pathway was the reduced assembly pathway with the largest flux and 19 intermediates. Adapted with permission from Earnest, T. M.; Lai, J.; Chen, K.; Hallock, M. J.; Williamson, J. R.; Luthey-Schulten, Z. Toward a Whole-Cell Model of Ribosome Biogenesis: Kinetic Modeling of SSU Assembly. *Biophysical Journal* **2015**, *109*, 1117–1135. Copyright 2015/ELSEVIER. (b) Typical domain composition of ABC transporter in Gram-positive organisms. SBP: Substrate Binding Protein, TMD: Transmembrane Domain, NBD: Nucleotide Binding Domain. (c) Nucleotide metabolism of GTP and dGTP with reactions catalyzed by protein complexes circled. ABC transporter rnsBACD import guanosine and deoxyguanosine in GSNabc and DGSNabc reactions.

In the previous Syn3A WCMs, we used representative subunits to specify the abundance of complete macromolecular complexes. For example, RNAP subunit *α* encoded by gene *rpoA*/0645 was picked for RNAP, and degradosome subunit RNase Y encoded by *rny* /0359 degradosome. For enzymatic protein complexes in metabolism, the Gene-Protein-Reaction (GPR) rules^2^ were applied to the catalyzed reactions. For example, ABC transporters in Gram-positive organisms have a domain composition of two peripheral nucleotide binding domains (NBD) that provide ATP hydrolysis, two transmembrane domains (TMD) that form the permease channel, and one substrate binding protein (SBP) that delivers substrates to TMDs^27^ as shown in Figure 1(b). The nucleoside ABC transporter rnsBACD comprises 4 distinct subunits of proteins RnsD/0008 (TMD), RnsC/0009 (TMD), RnsA/0010 (2NBDs), and RnsB/0011 (SBP) with 1:1:1:1 stoichiometries, where protein subunit RnsA/0010 corresponds to two NBDs.^28^ Previously, we used the AND GPR rule on two subunits RnsC/0009 and RnsA/0010 to take their minimum to represent the abundance of the ABC transporter rnsBACD in the membrane when simulating the import of nucleosides in the nucleotide metabolism marked in Figure 1(c). For completeness, the functional roles of the complexes in this study are listed in Table 1, and visualized in metabolic maps as Figure S1. In the pioneering WCMs of *Mycoplasma genitalium* ^29^ and *E. coli*,^30^ the formation of protein complexes was simplified as an one-step reaction from multiple subunits to the complete complex without incorporation of the assembly pathways, assembly intermediates, and spatial heterogeneity of the subunits.

**Table 1:** Macromolecular Complexes and their Functions in Syn3A.

| Location | Name | Function | Init. Cplx.<br>Cnt |
| --- | --- | --- | --- |
| Cytoplasm | RNAP | Gene Transcription | 93 |
|  | Ribosome | Translation/Translocation | 500 |
|  | SMC-ScpAB | Structural Maintenance of<br>Chromosomes | 101 |
|  | Gyrase | DNA Gyrase | 122 |
|  | Topo-IV | DNA Topoisomerase | 78 |
|  | RNDR | Ribonucleoside-diphosphate<br>reductase | 50 |
| Membrane | Degradosome | mRNA Degradation | 120 |
|  | Sec Translocon | Translocation | 66 |
|  | ATP Synthase | Reversible ATP Synthesis | 108 |
|  | rnsBACD | Import nucleoside | 145 |
|  | Opp | Import amino acids | 241 |
|  | PotABC | Import spermine | 104 |
|  | ThiBPQ | Import thiamine diphosphate | 20 |
|  | Pst system | Import phosphate ions | 123 |
|  | ribflECF | Import riboflavin | 10 |
|  | p5pECF | Import pyridoxal phosphate | 10 |
|  | 5fthfECF | Import 5-Formyltetrahydrofolate | 10 |
|  | nacECF | Import Nicotinate | 150 |
|  | coaECF | Import Coenzyme A | 10 |
|  | KtrCD | Import K<sup>+</sup> | 38 |
|  | Fak | Phosphorylation of Fatty Acid | 206 |

*In vivo* macromolecular complex assembly is believed to be evolutionarily optimized for efficiency in both time and resource usage. A key requirement is to the avoidance of kinetic traps—intermediates that are not fully assembled but cannot proceed productively.^14^ Recent in vitro theoretical studies have shown that kinetic hierarchies—arising from intrinsic differences in subunit affinities and association rates, as well as the timing of subunit supply—can significantly reduce kinetic trapping and improve yield. ^31–33^

Taking into account the different dimensionalities, the 3D cytoplasm, the 2D membrane, and the 1D DNA, and their coupling within a single cell,^34^ the assembly landscape is even more complicated. The association kinetics of 3D diffusive proteins have been extensively studied through a series of theoretical works.^35,36^ The impact of dimensionality reduction—from three-dimensional (3D) diffusion in solution to two-dimensional (2D) or even one-dimensional (1D) constrained motion—was first proposed by Adam and Delbrück in 1968^37^ and has since been quantified by both theoretical and experimental studies. Analytical models have shown that 2D confinement can accelerate dimerization by increasing the effective concentration and the probability of productive encounters.^38,39^ This prediction is supported by experimental comparisons of bimolecular association in solution versus on membranes.^40^ Nevertheless, only a few studies have reported absolute association rates for membrane-anchored proteins or DNA, primarily in the context of signaling pathways, ^40,41^ and comparable data for the assembly of multi-subunit membrane-embedded protein complexes remains scarce.

Here, we investigate the assembly kinetics of 20 complexes and the reduced model of ribosome biogenesis (See Table 1 for the total 21 complexes and their functions) in the context of our whole-cell model of Syn3A. Complexes’ compositions were determined by cross-checking the existing genome/proteome annotation^5^ and the homology-based function annotation. The assembly pathways, as series of bimolecular association reactions, of the ribosome,^24^ RNAP,^14^ and ATP synthase^42^ were taken as reported. The others’ pathways were inferred from interactions between subunits with enforced association order to avoid the kinetically trapped states. Specifically, only for ATP synthase, multiple strategies were used to discriminate the formation of one pair of kinetically trapped intermediates to improve the assembly efficiency. The heterogeneity of association rates was introduced for ribosome^15,24^and ATP synthase^42^ with experimental support, while not considered as approximations for other complexes. The whole-cell network for the translocation of membrane proteins in Syn3A were curated as shown in Figure 2. For membrane complexes, we varied the 2D association rates to investigate the effect of association rates in the context of coupled gene expression that further quantified by an analytical formula of the unassembled fraction for a simple dimerization model that correlated with transcription, translation, and mRNA degradation.

**Figure 2.**
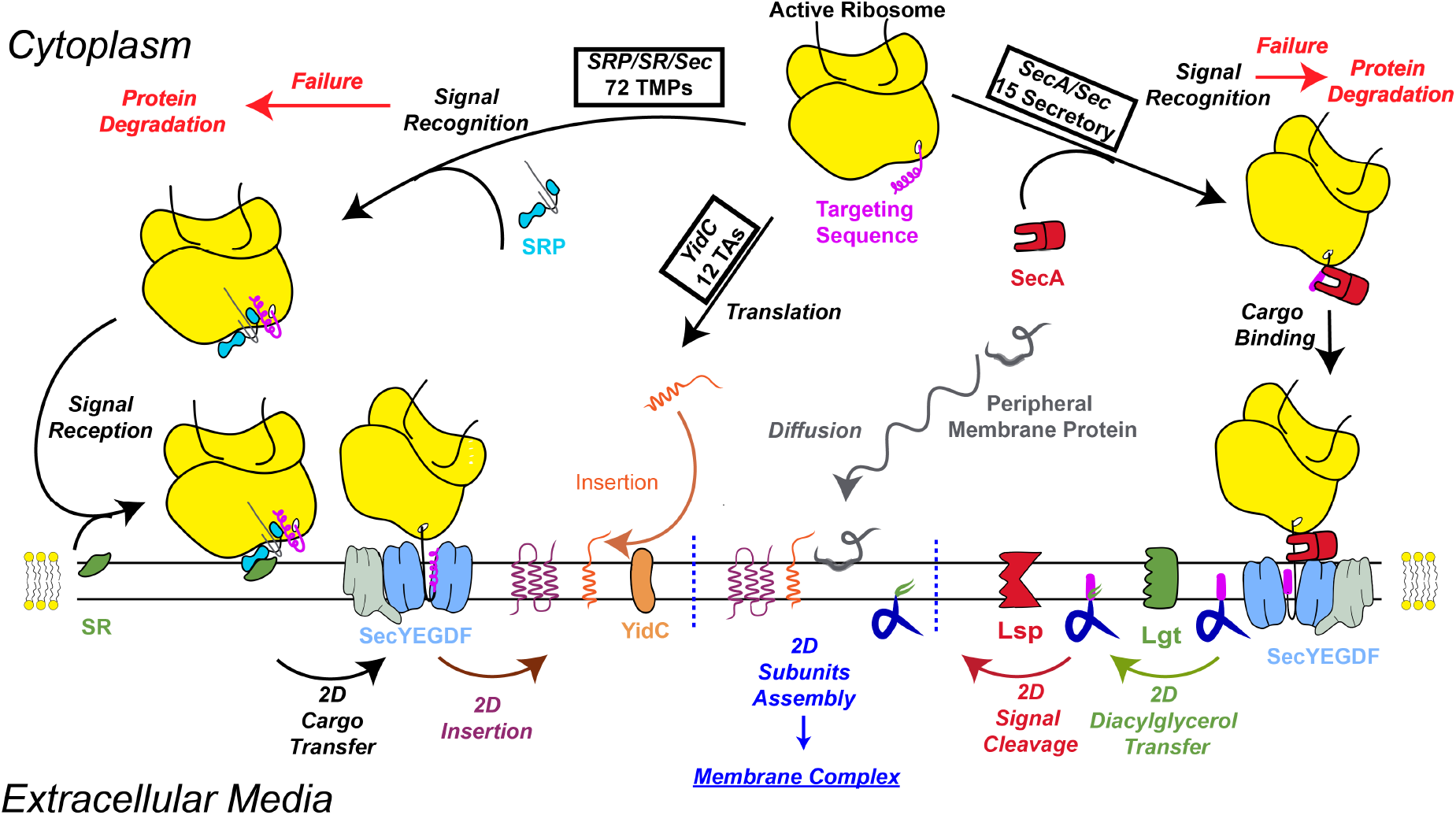
Translocation pathways for transmembrane proteins, and lipoproteins, and peripheral membrane proteins in Syn3A. Protein subunits’ localizations are classified as the cytoplasm, peripheral membrane, transmembrane, and secretory in Syn3A. Most kinds of transmembrane proteins (72/84) are co-translationally inserted via the SRP/SR/Sec pathway that recognized by SRP during translation. 12 unique kinds of tail-anchored (TA) membrane proteins can be post-translationally inserted via YidC.^12^ Secretory proteins are translocated via the SecA/Sec translocon that recognized by SecA; 13 kinds of lipoproteins, after modification anchor to and diffuse within the outer membrane leaflet, while 2 others are secreted to extracellular space. The failure of targeting in cotranslational translocation is considered and will lead to protein degradation by transmembrane protease, FtsH. Peripheral membrane proteins reversibly associate with the membrane by binding to transmembrane proteins and/or membrane lipids, thereby carrying out their functions at the membrane interface. Proteins in cytoplasm (omitted) are translated and then function in the cytoplasm. Reactions constrained on membrane are labeled as 2D. Signal reception in SRP/SR/Sec and cargo binding in SecA/Sec are coupled reactions between 3D cytoplasm and 2D membrane. Adapted with permission from Du Plessis, D. J.; Nouwen, N.; Driessen, A. J. The Sec translocase. *Biochimica et Biophysica Acta (BBA) - Biomembranes* **2011**, *1808*, 851-865. Copyright 2011/ELSEVIER

We evaluate the results of the augmented WCM of Syn3A by the assembly kinetics of all 21 complexes with the full translocation network of membrane proteins by simulating 100 replicates over the cell cycle. The used spatially homogeneous hybrid stochastic-deterministic algorithms are available in the Lattice Microbes software.^8,43^ In doing so we can separate out the timescales for gene expression, membrane insertion, complex assembly, and metabolism. By omitting the explicit evaluation of diffusion and incorporating its average effect into the binding rates, the simulations require about 6 hours to cover the complete cell cycle that can easily run in parallel for tens of cell replicates. The code of augmented Syn3A’s WCM is available on Github.

Finally, given the recent success of deep learning algorithms to analyze various biological data sets,^44^ and specifically their application to learn low-dimensional representations of the data,^45^ we examine the time-dependent variation of the data over the cell cycle from the simulation. The simulated time-dependent quantities contain information about the divergence of cell phenotypes. To investigate how metabolic profiles diverge, we analyzed the metabolomics of simulated cells using a machine learning analysis of the trajectory inference pipeline. Specifically, an autoencoder architecture was used to learn a low-dimensional representation of all 148 time-dependent intracellular metabolite concentrations. ^45^ Then, these learned embeddings were used to construct a trajectory tree,^46^ and each lineage reflects a specific cell population characterized with a distinct pattern of metabolomics. To identify potential drivers of cell divergence, we performed differential analysis of metabolite species and reactions at lineage branching points. We also conducted comparative analysis of metabolomics with and without additional complex assembly to show that incorporation of complex assembly prevents the overestimation of transporter, leading to elevated import of nutrients and more generation of energetic metabolites.

## Methods

### Hybrid Stochastic/Deterministic Algorithm for WCM of Syn3A

Lattice microbe (LM) was developed to simulate stochastic processes in biological cells using the Chemical Master Equation (CME) and the Reaction Diffusion Master Equation (RDME).^43,47,48^ Through a python hook, other algorithms can also be incorporated, including the ordinary differential equation (ODE, for metabolic kinetics)^5,8^ and Brownian dynamics (BD, for chromosome dynamics).

In this study, we simulate the kinetics at the whole-cell scale in Syn3A using a spatially homogeneous hybrid stochastic/deterministic algorithm CME-ODE, first published on the yeast’s galactose switch model and then applied to Syn3A.^5,8^ The genetic information process and the macromolecular complex assembly were simulated in CME, and the metabolism simulated in ODE. See File S1 Section Communication between Genetic Information Process and Metabolism for detailed explanations. The diffusion of particles are not explicitly simulated in the spatially homogeneous CME. Organization of the chromosome through Brownian dynamics and its interaction with nucleoid associated proteins are not included, but the assembly kinetics of these proteins and DNA replication are simulated.

Total 100 unique and independent cell replicates were sampled to capture the stochasticity of gene expression. As shown by the convergence of produced protein FtsY/0429 in Figure S7(a), both mean and variance stabilize to a plateau with cell replicates larger than 50, confirming that the adequacy of 100 cells for estimating expected behavior.

### Genetic Information Processes and Metabolism

As the majority of kinetic parameters remain the same as those used in our previous spatially homogeneous WCM of Syn3A, ^5^ we discuss only the few relevant modifications made and how the kinetics of complex assembly formation was incorporated into the genetic information and metabolic processes.

In Syn3A, DNA replication initiation occurs by sequential binding of the multi-domain DnaAs to the DNA signature sequences near *Ori* region to start bidirectional replication of the circular genome. Only one round of replication initiation and replication is now allowed as a low *Ori:Ter* ratio of 1.21 was calculated from DNA sequencing experiments (available at NCBI SRA PRJNA1257452). Initiation of DNA replication can begin after 20 or more DnaA proteins (Domain III) are bound to the ssDNA near the origin and following 3 DnaA (Domain IV) binding to the 3 DnaA boxes (each 9 NT) on the dsDNA. The previous constraint of 30 DnaA filament along the ssDNA was reduced after calculating the minimal size of the DNA bubble required to allow the binding of helicase in the replisome complex. Furthermore, the on and off rates of DnaA binding to ssDNA during filament formation obtained from smFRET experiments^49^ were modified from the average quantities (1 × 10^5^ to 1.4 × 10^5^ M^−1^s^−1^) and from (0.55 to 0.42 s^−1^), respectively, that were measured for genetic constructs with DnaA boxes similar to Syn3A’ *Ori* preceding the ssDNA signatures. We approximated the loading of two replisomes in the ssDNA pocket by binding the representative DNA polymerase III subunit *δ* HolA/0044 twice at a fast rate of 10^6^ M^−1^s^−1^ as long as the filament is longer than 20. Then, the replication of the circular genome happened bidirectionally from *Ori* to *Ter*, gene by gene.

As in the previous model, each gene is independently expressed in transcription, and its associated mRNA translated and degraded. What is different in this model, these processes begin with an irreversible binding step (RNAP to DNA gene, mRNA to ribosome or degradasome, respectively) followed by elongation/(de)polymerization reactions. By explicitly simulating the binding, we can identify the occupancy states of the RNAP, ribosome and degradosome and determine if they are actively processing the mRNA.

The Hofmeyr rate form^50^ is applied to catalyzed template-directed polymerization of genes in DNA replication, RNAs in transcription, and proteins in translation as described in our previous work.^5^ For RNA synthesis (i.e. transcription elongation), the enzyme and template pair is RNAP and DNA, and for protein synthesis (i.e. translation elongation), the pair is ribosome and mRNA. The formula for the Hofmeyr rate—excluding the step where the enzyme binds to the template polymer—takes the form:

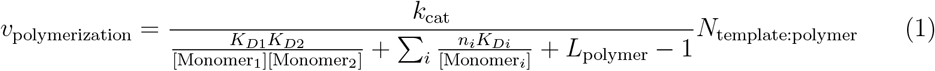

Here, *k*_cat_ is the intrinsic elongation speed of enzyme on the template in units of length (bp/nt/aa) per second. *K*_*D*,1_ and *K*_*D*,2_ are the dissociation constants for the first two monomers (Monomer_1_ and Monomer_2_) in the polymer sequence. *K*_*D,i*_s are the dissociation constants of monomers of type *i* in the polymer (e.g., ATP, CTP, UTP and GTP for RNAs, amino acid charged tRNAs for proteins), and *n*_*i*_s are the numbers of type *i* monomer in the polymer. Monomer concentrations [Monomer] are dynamic during the cell cycle, and shortages, which occur if consumption exceeds synthesis, can slow or in some cases even halt chain elongation. For transcription, the monomers are NTPs with *K*_*D*,NTP_ of 0.1 mM.^5^

Gromadski and Rodnina reported the dissociation rate of 2 *µ*M for Phe-tRNA^Phe^ with mRNA-programmed ribosome with cognate UUU codon. ^51^ Later in 2008, Ledoux and Uhlenbeck quantified the binding of ten different aa:tRNAs with their respective fully complementary cognate codon in *E. coli* to show the dissociation constants were uniform around 1.5 nM.^52^ Here, we assume a uniform dissociation rate for all aa:tRNAs *K*_*D*,aa:tRNA_ of 1 *µ*M, one order of magnitude smaller than the typical concentration of aa:tRNAs of 10 *µ*M.^5^ 1 *µ*M will not hinder the translation with the rich monomer pool, and also reflect the halt of translation upon the transient shortage of certain aa:tRNAs as shown in Figure S7(d). Minor variation of protein synthesis speed of up or down by 10% was quantified with *K*_*D*,aa:tRNA_ of 1.5 nM or 2 *µ*M. Using a uniform dissociation rate for all aa:tRNAs, we are not differentiating the binding affinities of different aa:tRNAs to their codons in active ribosomes. Thus, the overall translation rates in Hofmeyr rate law would be dependent on the length of the protein sequence, the numbers of each type of amino acids in this protein and the concentrations of corresponding aa:tRNAs. Therefore, the translation kinetics still vary from protein to protein based on their length and composition of sequences.

The expression levels of proteins were differentiated by introducing proxy promoter strength that scaled with proteomics abundances at the transcription process. The detailed reaction schemes and parameters of the gene expression are provided in File S1 Section Genetic Information Process. In addition, the parameters were summarized in File S3.

The kinetics of cotranslational translocation of membrane proteins involving SRP/SR, SecA, YidC and Sec translocon upon production of the target sequence are now explicitly included. The detailed reaction schemes and parameters of protein translocation are provided in File S1 Section Translocation Networks in Syn3A.

The 175 metabolic reactions that connect 148 intracellular metabolites and 49 extracellular metabolites in the growth media are the same as previously published,^5^ except the concentrations of any holocomplexes now depend on the assembly kinetics instead of just the representative subunit (Exact reactions in Figure S1, and the list of related complexes in Table 1). In the previous WCM of Syn3A,^5^ 16% of the 207 simulated cells died due to depletion of phosphoenolpyruvic acid (PEP) and then ATP in the main glycolysis pathway. This motivated a search for other parameters published in the literature for the phosphorelay system and some of the downstream glycolysis pathways to avoid losing cell replicates. (See File S1 Section 1.1 Metabolism for detailed explanation; See File S3 for the current set of metabolic parameters). As done previously,^5^ the thermodynamic analysis of the kinetic parameters was repeated to assure the correct free energy drop along the glycolysis pathway as shown in Figure S2. We also adjusted the import rates of (deoxy)nucleosides based on the requirements from the cellular activities to maintain the sufficient and balanced pool of NTP and dNTPs.

As previously done,^5^ the time-dependent surface area was calculated as the sum of the area of transmembrane proteins and membrane lipids assuming symmetric cell division starting with a spherical cell of radius 200 nm obtained from cryo-electron tomograms.^6^

### Abundances of Protein Complexes and Involved Protein Subunits

We initialized the macromolecular complexes and the involved protein subunits based on the existing proteomics with the following strategy. A holocomplex refers to a complete, functional macromolecular complex that includes all of its essential subunits or cofactors required for full biological activity. Initial counts of protein holocomplexes (annotated as *Init. Cplx. Cnt*) were estimated by their composition and proteomics counts of the subunits, except the ribosome, whose initial count of 500 was derived from cryo-ET tomograms.^5,6^ Proteomics coverage for short transmembrane subunits is often not accurately reported, so the counts of the holocomplex were based only peripheral protein subunits for ABC transporters and ATP synthase. Exceptions also included using the SecY subunit count to represent the Sec translocon, the SMC subunit to SMC-ScpAB, and the EcfT subunit to the ECF module, under the argument that these subunits are the major contributor in both structure and function of the holocomplexes. Complex assembly intermediates were initially assumed to not exist.

Another subsequent question was how to partition the abundances of protein subunits measured in the proteomics study (annotated as *Exp. Ptn Cnt*) into free proteins (annotated as *Init. Free Ptn Cnt*) and proteins already incorporated into complexes (annotated as *Init. Assembled Ptn Cnt*). The number of proteins incorporated into complexes was calculated by multiplying the initial counts of complexes by their stoichiometries, summing across complexes if a protein appeared in multiple complexes. After subtracting the number of proteins incorporated into complexes from the measured proteomic abundances, the remaining counts represented the initial free proteins. If this calculation resulted in negative numbers, these were reset to zero. Thus, the total initial protein count (annotated as *Init. Total Ptn Cnt*) was defined as the sum of the initial free count and the count incorporated into the complexes. See File S2 for detailed partitioning of the initial counts of 455 proteins. Ribosomal proteins exist both as free precursors and in assembly intermediates, typically less than approximately 5% of the total intact ribosomes.^17^ Therefore, we assumed that at least 5% of ribosomal proteins are in a free state. Specifically, for ribosomal proteins with measured proteomics counts lower than 525 (representing 500 proteins incorporated into ribosomes and 25 free), the initial free protein count was set to 25. For proteins with counts greater than 525, the initial free count was calculated as the proteomics count minus 500.

### Initial Conditions

In the spatially homogeneous CME-ODE kinetic simulation, which does not account for spatial heterogeneity, the state of the system is described by the time-dependent abundances/quantities of each species.

The computational growth medium and initial concentrations of intracellular metabolites remain the same as in our previous model (see File S2 for exact conditions). We assume that the medium concentrations remain constant throughout the 100-minute cell cycle.

The simulation begins with a single copy of the complete circular chromosome, with no DnaA bound to the *Ori*. Most rRNAs are bound to ribosomal proteins and incorporated into ribosomes; therefore, only two free copies each of 5S, 16S, and 23S rRNAs, from each of the two rRNA-coding genes, were initialized. A Poisson distribution was used to sample the initial count of each unique mRNA species. The mean abundance of each mRNA that used as the sole parameter of the distribution was calculated as the steady-state abundance under fixed cell growth. As a result, the total initial number of mRNA molecules was approximately 400, which doubled during the cell growth. The initial abundances of proteins and macromolecular complexes were assigned as described in the preceding sections.

### Compositions and Assembly Pathways of Protein Complexes

The protein complexes existing in Syn3A were determined by the annotations of the functional protein complexes discussed in previous research on Syn3A. ^2,5,7,9,53^ The compositions were determined through the comparative studies of these complexes in other model organisms, and also by the prediction on the structure organization of the protein subunits. The assembly pathways of the holocomplexes, if not reported, were inferred from the deposited subcomplex/intermediate structure and the interactions of the subunits. See File S1 Section Macromolecular Complexes in Syn3A and Assembly Pathways for detailed discussions. The list of macromolecular complexes, their functions and initial counts are listed in Table 1. The detailed compositions and assembly pathways of each complexes are in Table S2 and Table S3. Because Syn3A lacks direct measurements, we adopted assembly pathways and rates from model organisms as proxies, given the conserved structures and roles of these complexes. While this provides a mechanistic basis, organism-specific variations in protein subunits and cellular conditions factors in Syn3A may alter the assembly landscapes, and future measurements will refine these assumptions.

Complex assembly is constrained to proceed through a predefined hierarchy of bimolecular association reactions. A key approximation in our model is the exclusion of kinetic traps for most complexes, with the exception of ATP synthase, where kinetic traps were explicitly incorporated to evaluate their impact on assembly efficiency.

The ribosomal L7/L12 stalk is an exception to the otherwise 1:1 stoichiometry of ribosomal proteins. Structural studies in *E. coli* have shown that the stalk contains two L7/L12 dimers (four monomers) in LSU, which are essential for recruiting and activating translation factors through interactions mediated by the C-terminal globular domain. ^54^ The proteomics count of L7/L12 protein is 765. In simulations that included four copies, we initialized 2000 L7/L12 proteins that incorporated into the ribosome and 25 remained free. Based on solution structures of the L7/L12 dimer,^55^ we assumed rapid dimerization (rate constant of 10^6^ M^−1^ s^−1^) independent of LSU incorporation. L7/L12 dimmer associates into LSU as the last step. The assembly pathway of LSU with four copies L7/L12 proteins are in File S3.

When four copies were modeled, 23S rRNA synthesis remained the dominant bottleneck, but the increased transcriptional demand of the L7/L12 gene slightly reduced 23S rRNA production, decreasing the median number of fully assembled ribosomes at the end of the cell cycle from 400 to 375. Because this adjustment had no significant impact on ribosome assembly intermediates, cell growth, proteome doubling, or metabolomic profiles, the main text presents results from the model with one copy of L7/L12 protein while noting this refinement for completeness.

### Bimolecular Association Rate of Complex Assembly

Under the spatially homogeneous assumption in CME kinetics, a lumped association reaction of two particles to form a complex (A+B to C) contains first the relative diffusion between two particles and then the intermolecular interactions to overcome the barrier to form a stable complex.^36^

The heterogeneity and kinetic hierarchy of the assembly rates were incorporated to varying extends across different macromolecular complexes. For the reduced linear assembly pathway of ribosome SSU, association rates vary from 2×10^4^ M^−1^s^−1^ (S12 as the last binder) to 3.1×10^7^ M^−1^s^−1^ (S6 as the third binder). For LSU, guided by strong or weak protein binding cooperativities measurements by Nierhaus, ^15^ we assigned slow and fast association of 10^4^ M^−1^s^−1^ and 10^6^ M^−1^s^−1^. In the case of ATP synthase, we differentiated the binding rates among *ab*_2_, *α*_3_*β*_3_*γϵc*_10_, and *δ*, three modular intermediates at the last stage to demonstrate that kinetic hierarchy alleviated the kinetic trapped intermediates *α*_3_*β*_3_*γϵc*_10_*δ* and *ab*_2_*δ*, improving the yield. Meanwhile, reversible dissociation of trapped intermediates similarly achieved the improvements by escaping from the trapped states. However, due to the lack of quantitative data on intermediate populations, particularly for membrane complexes, and the limited tools to estimate the interaction strengths among subunits in high throughput, we adopted a simplified linear pathway with uniform bimolecular association rates for other complexes. This assumption promotes efficient assembly by eliminating the potential kinetic traps.

One association is diffusion-controlled when the rate of binding is limited by the rate at which the proteins diffuse through the medium to encounter each other, rather than by the intrinsic reaction kinetics of binding. Without considering steric specificity, the Smoluchowski equation (*k*_*D*_ = 4*πDR*, where *D* is the relative diffusion coefficient and equals *D*_1_ + *D*_2_, *R* is the contact distance between the centers of the two spheres) gave a binding rate on the order of 10^7^ M^−1^s^−1^ between 3D diffusive cytoplasmic proteins. Considering the steric specificity of the proteins, the diffusion-controlled association rates were estimated between 10^4^ and 10^6^ M^−1^s^−1^ with no inclusion of long-range attractions like electrostatics.^35^ The electrostatic attraction could enhance the binding kinetics much faster folds via stabilizing the transient association intermediates for certain complexes.^36,56^ The faster rates have been observed in the assembly of ribosome SSU. We chose to ignore the contributions of long-range attractions for cytoplasmic protein complexes (except for ribosome biogenesis) and adopted the association rate for cytoplasmic complexes, 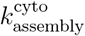 of 10^5^ M^−1^s^−1^. 10^5^ M^−1^s^−1^ is a relatively fast rate compared to gene expression, thus not hindering the assembly of RNAP, RNDR, and other cytosolic complexes by long association waiting times among subunits.

The association of protein subunits can be significantly altered by their localization. Transmembrane proteins diffuse one to two magnitudes slower in confined 2D membrane than cytoplasmic proteins. For example, diffusion coefficient of 0.02 *µ*m^2^s^−1^ was observed for the transmembrane protein LacSΔIIA-GFP in Gram-positive *Lactococcus lactis*, and a much faster coefficient of 7 *µ*m^2^s^−1^ for its counterpart cytoplasmic protein GFP^57^). However, the pre-orientation of reactive regions and the longer lifetime of the transient complex largely compensate for the reduced diffusion coefficient.^34,58^ The same logic applies to the peripheral membrane proteins that slide on the membrane.

Quantifying absolute association rates in two dimensions (2D) remains rare. An early study by Gavutis *et al*. in 2006^41^ reported 2D association rates between interferon receptor domains on solid-supported lipid bilayers ranging from 6.6 ×10^−3^ to 4.6 ×10^−2^ *µ*m^2^ s^−1^. More recently, Huang *et al*. measured a 2D association rate of 2.0 ×10^−3^ *µ*m^2^ s^−1^ for complementary DNA strands on supported membranes.^40^ However, these values pertain to membrane-anchored particles, not fully embedded transmembrane proteins. Due to the limited understanding of intrinsic 2D association kinetics, we explored how varying this parameter affects complex formation. Specifically, three values for the membrane association rate 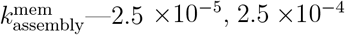, and 2.5 ×10^−3^ *µ*m^2^ s^−1^—were applied to reactions between peripheral and transmembrane subunits, as well as between two transmembrane subunits. These rates correspond to effective 3D association rates of 10^3^, 10^4^, and 10^5^ M^−1^ s^−1^ for a 200-nm-radius spherical volume. In the chemical master equation framework, the reaction propensity *a*_*r*_ (the probability per unit time) is computed as *a*_*r*_ = *k*_stochastic_*n*_*a*_ *n*_*b*_, where 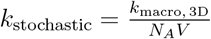 in solution and 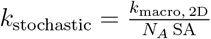 on membranes, with *V* as the cell volume and SA as the surface area.

### Construction of Protein Translocation Network

To clarify the three translocation pathways in Syn3A shown in Figure 2, the related entities were summarized in Table 2. The total number of ribosomes was estimated from cryo-electron tomograms of a cell with a radius of 200 nm; _6_ the abundances of proteins (subunits) came from our proteomics study using quantitative mass spectrometry.^2^

**Table 2:** Abundances of Macromolecules in Syn3A’s Translation/Translocation Network ^*a*^.

| Category | Translocation Entities | Absolute Count | Localization<sup>b</sup> | Ess.<sup>c</sup> |
| --- | --- | --- | --- | --- |
| Ribosome | Total Ribosome | 500 | CP | E |
| SRP | Ffh/0360 | 365 | CP | E |
|  | 4.5S RNA/0049 | - | CP | E |
| SR | FtsY/0429 | 88 | PMP | E |
| SecA | SecA/0095 | 573 | CP | E |
| Sec Translocon | SecY/0652 | 66 | TMP | E |
|  | SecE/0839 | 8 | TMP | E |
|  | SecG/0774 | 4 | TMP | Q |
|  | SecDF/0412 | 215 | TMP | E |
| YidC | YidC/0908 | 120 | TMP | E |
| Lipoprotein Modifier | Lgt/0820, 0818 | 13 | TMP | E |
|  | LspA/0518 | 3 | TMP | E |
| Proteases | Lon/0394 | 757 | CP | Q |
|  | FtsH/0039 | 148 | TMP | E |
|  | YqgP-GplG/0516<sup>53</sup> | 63 | TMP | Q |

<sup>a</sup> Absolute Counts in the initial of the simulation
<sup>b</sup> CP: Cytoplasmic protein, PMP: Peripheral membrane protein, TMP: Transmembrane protein
<sup>c</sup> Ess.: Essentiality, E: Essential, Q: Quasiessential, N: Nonessential

The SecYEGDF translocation machinery (bacterial SecY family member ^12^ with auxiliary SecDF subunit,^59^ hereafter Sec translocon) and YidC insertase (bacterial Oxa1 family member^12^) are responsible for the insertion of transmembrane proteins and secretory proteins. YidC insertase, independent of the Sec translocon, could promote the insertion of a subset of TMPs with short translocated regions, ^12^ as well as functioning in conjunction with the core Sec translocase to assist the insertion of long translocated regions into the membrane.

The signal recognition particle (SRP, consisting of protein Ffh/0360 and 4.5S non-coding RNA from gene 0049) and the signal receptor (SR, FtsY/0429) mediate the cotranslational translocation of transmembrane proteins. SecA/0095 was previously known to insert secretory protein across the membrane in concert with the SecB chaperon and the Sec translocon,^60^ yet recent selective ribosome profiling proved its role in cotranslational recognition and insertion without chaperons.^61^ It should be noted that the count of SRP is almost the same as ribosomes, while in *E. coli*, the number of ribosomes largely exceeds SRP and SecA by up to two orders of magnitude. This could lead to the loss of SRP targeting specificity, as verified by selective ribosome profiling experiments in *E. coli* when overexpressed SRP bound to cytoplasmic, periplasmic, and outer membrane proteins.^62^ In Syn3A, this reduced substrate specificity could presumably decrease the efficiency of translocation of transmembrane proteins with unnecessary Sec translocon occupation. Without measurement of ribosome/SRP interactome from selective ribosome profiling, here we assume that SRP can only target transmembrane proteins.

Two lipoprotein modifiers, lipoprotein diacylglyceryl transferase (Lgt/0818 and Lgt/0820, EC 2.5.1.145) and lipoprotein signal peptidase (SPaseII, LspA/0518, EC 3.4.23.36, PDB entry 5DIR) were found in Syn3A. Due to the minimized genome, Syn3A only possesses DnaK/DnaJ/GrpE and ClpB chaperon systems for protein folding and disaggregation, respectively. Two transmembrane proteases, FtsH/0039 and YqgP-GplG/0516, and one cytoplasmic protease Lon/0394 degrade proteins to peptides, and five peptidases further decompose them into amino acids to prevent protein aggregates and to recycle amino acids.

Three curated translocation pathways summarized in Figure 2 are, the cotranslational SRP/SR/Sec and SecA/Sec pathways and the post-translational sole YidC pathway. Substrate proteins of different translocation pathways were determined with bioinformatic predictions, together with gene functional annotations.

In the SRP/SR/Sec pathway, translation starts from mRNA diffusing and binding to a ribosome, forming the ribosome nascent chain complex, and the elongation of the polypeptide chain will expose the targeting sequences. For the SRP/SR/Sec pathway, SRP can diffuse and recognize the ribosome to form a SRP_Ribosome complex. Since bacterial SRPs do not have an additional Alu domain to arrest translation elongation, the polypeptide chain is allowed to grow with normal speed.^62^ The binding of SRP to SR occurs through the interactions of each GTPase domain near the peripheral membrane and is fast enough with *k*_*on*_=10^7^ M^−1^ s^−1^ with SRP when loaded with ribosome.^63^ The cytosolic loops of the Sec translocon could interact the A domain of FtsY and the ribosomal protein L23 at the nascent polypeptide exit site, thus facilitating the effective formation of the quaternary complex with *K*_*d*_ =2 *µ*M.^64^ The Sec translocon activates the GTP hydrolysis of SRP_SR targeting complex and the handover of ribosome to the Sec translocon channel at a rate of 0.95 s^−1^.^64^ The last step is the elongation and translocation, and the translation elongation serves as the driving force in inserting the polypeptide chain into the membrane. The rate of insertion and folding of transmembrane proteins was estimated to be above 20 aa/s based on one *in vitro* study on protein lactose permease inserted by SecYEG and YidC,^65^ while the chain elongation speed was 12 aa/s, thus the translation elongation is the rate-determining step. The two-step kinetics of simultaneous translation and translocation was combined to one reaction, where the product was the synthesized and translocated membrane proteins. Correspondingly, Hofmeyr rate law was used to depict the rate of the lumped reaction. ^50^

YidC and Sec translocon collaborate to insert different parts of membrane proteins, where YidC is responsible for short hydrophilic segments and Sec translocon for long hydrophilic segments.^12^ The copurification experiment indicated that YidC binds to the Sec translocon by interacting with the SecDF subunit.^66^ Here, it was assumed that YidC will only associate with the Ribosome_Sec translocon complex transiently via its interaction with SecDF to finish the insertion, after which it will disassociate from the Sec translocon.

The failure of cotranslational targeting occurs when the emerging polypeptide chain grows too long, weakening the interaction between the ribosome and SRP. According to selective ribosome profiling, the length of the nascent polypeptide chain lies between 50 and 100 residues when SRP binds to ribosome. ^62,67^ Here we use the metric of the length of polypeptide chain shorter than 100 AAs before the SRP recognition. In this case, the degradation of the protein presumably occurs to prevent the protein aggregates and to recycle amino acids. From the mutant experiments of two proteases, Lon and FtsH in *Mycoplasma pneumoniae*,^68^ membrane proteins are the main targets for the transmembrane protease FtsH. Considering the transmembrane region in the nascent polypeptide chain as the hydrophobic signal target, here it was assumed that the degradation of the unsuccessfully translocated transmembrane protein occurs through its binding to FtsH, supported by the *in vitro* degradation of SecY by FtsH.^69^ The degradation kinetics and the cost of ATP depend on the stability of the protein substrate and the hydrolysis rates of ATP.^70^ Here we take the maximum degradation rate 0.22 min^−1^. The further decomposition from the peptide to amino acids by peptidase is approximated to be combined within this reaction. We assume that ATP energizes the degradation with one ATP per amino acid and that the entire protein is digested.

In the sole YidC pathway, YidC could function separately from Sec translocon on transmembrane proteins with only short hydrophilic translocated segments.^12^ Tail-anchored membrane proteins (TA) with one or two transmembrane regions (TMRs) and the targeting TMR located less than 65 amino acids from the C-terminus are post-translationally translocated by YidC, since the short amino acid distance leaves no time for cargo transfer to the Sec translocon. ^11^ In this pathway, cotranslational recognition of TMR by SRP is unnecessary, and after translation is completed, TAs could diffuse to the inner membrane and bind to YidC. Binding with YidC will assist in inserting a short TMR into the membrane without the Sec translocon. The rate at which YidC inserts TA proteins into the membrane was set at 1 s^−1^ under the argument that YidC incorporated only the short TMR of 20 AAs in the lipid membrane and the rate of insertion was faster than 20 aa/s.^65^

Translocation of secretory proteins could occur through both cotranslational and postranslational pathways. ^71^ In Syn3A, the previously well-recognized post-translational translocation pathway of secretory proteins^60^ is less likely. The translocation-competent state of the nascent polypeptide chain is required before/during the targeting and insertion, and this state is usually maintained by SecB chaperon which has been characterized in *E. coli*.^71^ Since SecB is not in Syn3A, and there is no direct evidence showing that existing chaperon systems DnaK/DnaJ/GrpE and ClpB or SecA could compensate for SecB’s role, therefore the translocation of translated nascent chain would not succeed.

The targeting factors for secretory proteins could be SRP/SR (more recognized for TMPs and preferring a high hydrophobic signal peptide) or SecA (more recognized for secretory proteins). With no measurement of the ribosome interactome with SRP and SecA from selective ribosome profiling to know the discrimination of substrates for SRP/SR and SecA, it was assumed here that the translocation of secretory proteins is facilitated by only SecA. Another factor that favors the cotranslational pathway is the absence of the trigger factor (TF), which hinders and delays SecA engagement, leading to post-translational translocation.^61,72^

In cotranslational translocation via SecA/SecC, SecA first recognizes the signal peptide emerging from the ribosome exit tunnel and directs the binding to the Sec translocon. In this fashion, protein misfolding is prevented and translocation is rapid. SecA selective ribosome profiling revealed that the length of the nascent polypeptide chain lies between 100 and 150 residues when SecA binds to ribosome,^61^ and here the recognition failure by SecA, similar to SRP, was also considered using the metric of 150 residues. However, lipoproteins 0481, 0605, and 0851 are of lengths 145, 102, and 68 AAs, thus we roughly assumed thresholds of half the protein lengths. SecA ATPase inserts the secreted polypeptide chain across the membrane under ATP hydrolysis, which is also enhanced by SecDF using the proton motive force. ^59^ Protein SecDF/0412 is a transmembrane protein with 12 TMRs and a large extracellular domain P1. As an auxiliary factor, SecDF is believed to be incorporated in insertion late after SecA, when its P1 domain binds to the unfolded protein on the outer leaflet membrane surface and pulling the chain out with repeated conformational transitions driven by the proton gradient.^73^ Here, it was assumed that the insertion by SecA and the pull by SecDF were much faster than the elongation of the polypeptide chain, and the Hofmeyer rate law was used to depict the rate of cotranslational translocation.

Modification of lipoproteins occurs upon translocation with Lgt and Lsp for proper function. Lgt transfers the diacylglyceryl group from phosphatidylglycerol (PG) to the sulfhydryl group of the cysteine residue in the lipobox, and then Lsp cleaves the signal peptide up to the cysteine residue, leaving an exposed *α*-amino group of diacylglyceryl-systeine.^74^ Mature lipoproteins anchor to the outer leaflet of the membrane by embedding the lipid-modified cysteine residue^75^ in Figure 2.

Extracellular proteins are also cotranslationally translocated across the membrane. However, without signal peptidase (SPase I) identified, it is assumed that the translocated proteins are released to extracellular media without signal peptide cleavage.

Proteins without signal peptides or transmembrane regions are translated in the cytoplasm. They could be further categorized into peripheral membrane proteins and cytoplasmic proteins. Peripheral membrane proteins could reversibly diffuse back/forth between the cytoplasm and the peripheral membrane region until they are trapped by specific and strong interactions, such as forming a complex with TMPs, The diffusive behavior on the membrane is generally constrained 2D sliding under nonspecific hydrophobic interactions with membrane lipids.^34^ The proteins in the cytoplasm are translated and then function in the cytoplasm. Under the spatially homogeneous assumption, no explicit diffusion was simulated, and instead, reversible exchange reactions of peripheral membrane proteins were implemented between the cytoplasm and the peripheral region to account for their spatial distribution.

The translocation pathways and diffusive behaviors of protein in different localization were summarized in Table S9. The implemented reactions of the preceding behaviors are summarized in Table S10.

### Revision of Protein Localization, and Identification of Targeting Sequences for Translocation

Targeting sequences were determined with the new prediction of two bioinformatics tools, Signal P 6.0^76^ and DeepTMHMM.^77^ SignalP 6.0, a machine learning-based tool, was used to identify signal peptides, short amino acid sequences that direct proteins to secretory pathways and are cleaved by signal peptidase (SPase) after translocation. In addition, it could differentiate the signal peptide for SPase, SPase I for Sec SPs in extracellular proteins, and SPase II for lipoprotein SPs in lipoproteins.^71^ DeepTMHMM was used to predict the locations of different protein segments, including the signal peptide and extracellular part of secretory proteins and the TMRs of transmembrane proteins. The prediction of signal peptides from DeepTMHMM was compared with that from SignalP to further verify our localization assignment of secretory proteins. The structural organization of TMRs determined the topology of transmembrane proteins, i.e. the positions and lengths of TMRs, and also intracellular and extracellular loops and their orientations, which affect the translocation mechanism. SignalP 6.0 and DeepTMHMM are available online. When performing SingalP 6.0, we chose Organism as other for bacteria, and Model as slow for accurate region borders of the targeting sequences.

Minor revision of protein localization from the previous publication were made. ^2,5^ A total of 455 proteins (subunits) are localized in four regions, 315 in the cytoplasm, 38 in the peripheral membrane, 84 in the transmembrane, 15 as secretory proteins, with 3 proteins left unidentified shown as pie chart in Figure S6.

The signal peptides predicted by SignalP all originate from the N-terminus and range in length from 15 to 28 residues. Transmembrane regions (TMRs), which serve as targeting sequences for transmembrane proteins (excluding tail-anchored proteins), typically begin within the first few to 50 residues from the N-terminus and span approximately 10 to 20 residues in length. See File S2 for the function, localization, signal peptide, and topology of each proteins. See Table S8 for the full list of tail-anchored membrane proteins.

### Machine Learning-Based Trajectory Inference for Metabolic Profile

To investigate how cells evolve into different metabolic phenotypes, we analyzed metabolic profiles using a three-step trajectory inference pipeline. First, we trained a deep neural network on the raw metabolite data to obtain a low-dimensional representation. The raw data set comprised 148 unique intracellular metabolites, collected at 6,000 time points, for each of the 97 cell replicates. Then we built a trajectory tree using this representation, where different lineages diverge from the same root/origin (due to the same initial concentrations) over time. Each lineage represents a different cell population, and we classified cells by projecting each cell’s trajectory onto every lineage, summing the path-to-lineage distances, and assigning the cell to the lineage with the smallest total distance. Finally, at the points where the lineages diverged, we identified metabolite species and reactions that are significantly different between emerging cell populations.

### Representation Learning via Autoencoder

In the first step, we applied an autoencoder to compress raw metabolite concentrations into low-dimensional representations. An autoencoder is a powerful tool in unsupervised representation learning^45,78,79^ and has been widely applied in biological data analysis.^80,81^ It consists of an encoder and a decoder. The encoder enables the compression of high-dimensional data into a low-dimensional representation, and the representation is optimized by reconstructing the original data via a decoder. This structure allows the model to preserve all essential signals in the representation while removing noise.

The metabolite concentrations recorded every second throughout a complete cell cycle are used as input for the autoencoder. In total, there are 148 intracellular species. The number of recorded time points varies between cells depending on the cell cycle duration, ranging from 93 to 110 minutes. The encoder is a fully connected three-layer network containing 64, 32 and 16 neurons, respectively. ReLU activation that takes the nonnegative part of its argument is used to introduce nonlinearity. The output of the final encoder layer serves as the compressed and learned representation of the input concentrations traces and is used in trajectory tree construction. The decoder mirrors the encoder’s structure, with an output layer matching the dimensionality of the input data. The optimal model is determined by minimizing the following loss function ℒ:

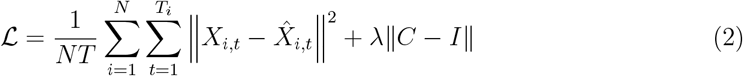

where *N* is 97 as the number of cell replicates, *T* is the summation of length of time points of all *N* cell replicates, *T*_*i*_ is the length of time points for cell *i, X*_*i,t*_ denotes the metabolite concentrations of cell *i* at time *t*, 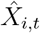 represents the reconstructed metabolite concentrations, *C* is the covariance matrix of latent factors, and *I* is the identity matrix, *λ* is the regularization coefficient. The first term quantifies the reconstruction loss, ensuring that the learned representation retains as much information from the original data as possible. The second term is a regularization term that decorrelates the different dimensions of the representations to prevent them from capturing redundant information. ^82,83^ The hyperparameter *λ* balances the weights of the loss of reconstruction and the regularization term. In practice, we select the value of *λ* to be 3 by grid search over a range of values.

### Trajectory Tree Construction and Cell Classification

In the second step, we constructed a trajectory tree with the learned representations using Slingshot.^84^ Slingshot is a lineage inference tool with two stages:

1. Identify global lineage structures by constructing a minimal spanning tree (MST) on clustered data points. Here we clustered the data points using the *k*-means algorithm. We determined the optimal number of clusters through the grid search and selected the value that allows the initial trajectory tree to capture the growth dynamics of the cells more effectively. The optimal number of clusters was found to be 9.
2. Refine each lineage using principal curves and estimate pseudo-time variables for cells along each lineage. The pseudo-time variable represents the growth stage of the cell along the lineage.

Each lineage on the tree represents the metabolite profile of a distinct cell population. Cells are assigned to the population that has the smallest distance to their trajectory path.

### Between-Lineage Differential Analysis

After cells are clustered into distinct populations, the next key question is to identify which metabolite species or reactions drive cell divergence. To address this, we performed a differential analysis between each pair of lineages. First, we define the branching point as the time at which the distance between the two lines exceeded a predefined threshold *ϵ*, indicating the starting point when cells start to diverge into different types. To identify hidden biological changes related to cell divergence, we compared metabolite concentrations and reaction fluxes at the branching point between different cell types. Two-sample *t*-tests are used to each metabolite and reaction to determine whether their differences between lineages are statistically significant. To adjust for multiple hypothesis testing and control false discovery rate, the Benjamini-Hochberg procedure is applied to adjust the *p*-values,^85^ and we considered a metabolite species or reaction significantly different if its adjusted *p*-value fell below significance level *α* = 0.05. We then quantified the magnitude of these differences using standard effect sizes (Cohen’s d statistic). Cohen’s d is calculated as the difference between the two group means divided by the pooled standard deviation, and the large Cohen’s d means a significant difference between the group means.

We also reported the separation time between each pair of lineages. To calculate this, we projected the cell trajectories that pass through these two lineages onto the tree and identified the time when each cell crossed the branching point. The separation time for this lineage pair was defined as the average of these times.

## Results and Discussion

### Time-dependent cell behavior from the augmented WCM of Syn3A

Before analyzing the complex assembly subsystem, we first examined whole-cell dynamics to provide context. The results in this subsection include simulations of ribosome assembly using a reduced pathway and membrane complexes with fast association rates.

Cell volume doubled at a median time of 67 minutes in Figure 3(a), after which the partitioning of the daughter chromosomes is assumed to occur. The cell cycle, defined as the time for surface area to double and the cells to divide, ranged from 95 to 111 with a median of 102 min in Figure S7(c), which is in good agreement with experimental measured 105 min.^2–4^

**Figure 3.**
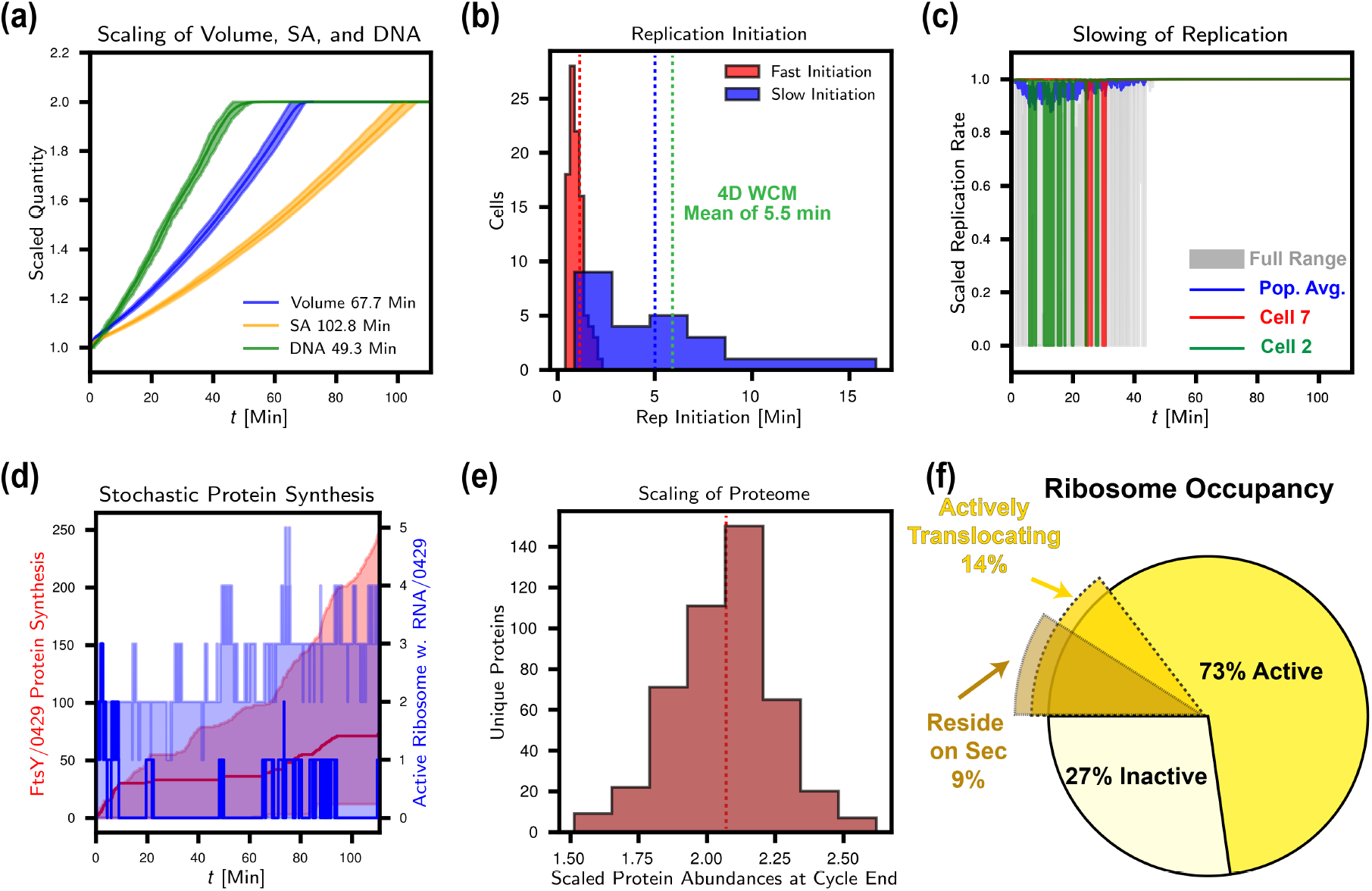
Time-dependent cell behavior from the augmented WCM Syn3A (a) Scaled chromosome, volume and surface area over the entire cell cycle. Median doubling times among cell population were shown in legends. (b) Comparison of replication initiation time with fast and slow DnaA binding rate with ssDNA with median of 1 and 5 min, respectively. The median time from 4DWCM is shown by green line. (c) Slowing down of gene replication due to the dTTP shortage. In cell replicates 2 and 7 that running out of dTTP, the replication rate was zero at a temporal resolution of 1 second, and recovered with dTTP synthesis in nucleotide metabolism. The population averaged replication rate fell to 90% of the maximum (the blue line), and the gray shaded area shows the full range of scaled replication rate. (d) Stochastic gene expression of FtsY/0429. The left y-axis is the synthesized FtsY/0429, and the right is the active ribosome translating the mRNA of FtsY/0429. The solid lines represent one single cell replicate, the shaded areas the full range. (e) Distribution of population averaged scaled protein abundance at the end of cell cycle with the median of 2.07. (f) Ribosome occupancy at steady-state. 73% out of total are actively translating mRNAs, 14.6% out of total are active in translocation of transmembrane proteins and lipoproteins, and 9% out of total are residing on Sec translocon.

Chromosome duplication, regulated by the replication initiator DnaA, is essential for the inheritance of genetic material and establishes the timing of gene doubling during the cell cycle. The higher DnaA binding rate with ssDNA on *Ori* gives unrealistic fast replication initiation between 0.5 and 2.25 min with the mean of 1 min in Figure 3(b), which means a very short B period. However, the 4DWCM including the explicit diffusion of DnaA predicted a longer initiation time of mean time 5.5 min, where some cell replicates requiring 10 min or longer to form the DnaA filament and initiate DNA replication. ^86^ Changing back to the slower DnaA binding rate in the spatially homogeneous CME-ODE WCM, the expected initiation time recovered with the median of 5 min. This highlights the influence of diffusion on the cellular behaviors, and the necessity to adopt different parameterization strategies with and without considering diffusion.

As the replication fork moves in a bidirectional manner along the circular chromosomes of half a million base pairs, each gene is replicated over a large time range. Gene *dnaA*/0001 was replicated quickly after the initiation, while second copies of genes near the terminus like *fakA*/0420 were available for transcription after 40 min in Figure S7(b). Progressive replication of the circular chromosome affects protein synthesis at the transcriptional stage and, consequently, downstream protein assembly. A notable example is the structural maintenance of chromosome (SMC–ScpAB) complex, encoded by *scpA*/0327, *scpB* /0328, and *smc*/0415. Because the *smc* gene doubles late in the replication cycle, SMC subunit production is delayed, limiting formation of the complete SMC–ScpAB complex.

On average, the entire chromosome was duplicated around 49 minutes as shown in Figure 3(b), 20 min faster than the predicted time in the previous spatially homogeneous model in^5^ where a slower DnaA binding rate was used. Apart from a faster initiation in this model, the main reason was the less halting of DNA replication from the shortage of deoxyribonucleotide monomers, as shown by the scaled replication rate in Figure 3(c). The DNA replication rate only dropped 10% in contrast to 25% drop in the previous model. ^5^ The elongation rates for transcription, and translation all show a slight decrease over parts of the cell cycle in Figure S7(d,e), but the majority cells are able to recover the monomer pools, and no cells were observed to run out of phosphoenol pyruvate (PEP).

RNAP, ribosome, and degradosome exhibit active fractions of 65%, 73%, and 14%, respectively, which remain pretty steady due to the synchronized assembly of all three complexes during the cell cycle. All three complexes roughly doubled their initial abundances at the end of cell cycle in Figure S7(f). The RNAP *α* subunit (RpoA/0645) had a proteomic count of 187 with a stoichiometry of 2, yielding an estimated initial RNAP abundance of 93 holoenzymes under a minimal assembly principle (full subunit list in Table S2). Consequently, *α* subunit was least synthesized during the cell cycle and limited the formation of RNAP from the supply side. The ratio of active RNAP is higher compared to the value of 36.2% for *E. coli* with 20 min doubling time because the lower RNAP to gene ratio in Syn3A of 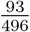 compared to 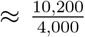 in *E. coli*.^87^ The high RNAP active fraction indicated strong competition for transcription among different genes. A notable emergent result was that genes with weak promoters were not transcribed in all cells within a single simulated cell cycle. However, the proteomic counts of RNAP subunits *α, β*, and *β*^*′*^ had a ratio of 1:3:3 that markedly deviating from the expected 2:1:1 stoichiometry observed in *E. coli* ^88,89^ and *Bacillus subtilis*.^90^ This discrepancy suggests the need for further investigation of the relative expression levels of these subunits using transcriptomic data.

The discreteness and stochasticity of protein synthesis is intrinsic at the single-cell level. A clear step-like trace of the synthesized protein FtsY/0429 is shown in Figure 3(d), where the stalled protein synthesis in the horizontal regions corresponded to no ribosomes translating this mRNA.

Upon transcription, the fate of the mRNAs is determined by the competition between its binding and translation by ribosomes into a protein, and its binding and degradation by degradosomes into NMPs. If one views the competition between the ribosome and the degradosome as a series of independent Bernoulli trials, where success means mRNA binding with the ribosome, and failure binding with the degradosome, then the distribution of the number of successes until the first failure occurs, which is the translations per mRNA transcript until mRNA degradation, follows a geometric distribution with the mean being the ratio of success over loss. The time-dependent ratio of mRNA binding with ribosome over binding over degradosome was calculated using 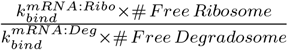. In our current implementations, the ratio between ribosome and degradosome binding rates was 6.3 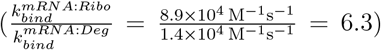. The number of available ribosomes over available degradosome was about 1.3 over the cell cycle, which further favor the mRNA to translation instead of degradation 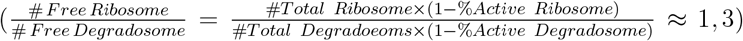.

Thus, the mean of translation per mRNA as 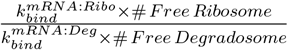 was 8.2. From our simulation, the translation events per each unique mRNA transcript were calculated by dividing the number of translated proteins by the transcribed mRNAs. The distribution ranged from 5.8 to 11, with a prominent peak around 7.7 that was close to the calculated 8.2 in Figure S7(h).

A key performance metric of the WCM is its ability to reproduce the duplication of protein abundances by the end of the cell cycle, given that cell division halves the cellular contents. The averaged scaled protein abundances range from 1.51 to 2.62, with only 32 of the 452 proteins having a value less than 1.8 in Figure 3(e). Two factors contributing to the distribution of the scaled abundance are the length of the protein and the initial proteomics counts. Shorter proteins with the lower initial counts will favor a higher scale abundance. Proteins longer than 1000 AAs or with initial counts greater than 800 almost failed to double over the course of a cell cycle. Polysomes, not included in this WCM, would serve as a mechanism to increase the expression levels of long proteins. ^5^ Proteins with low or zero initial counts in the proteomics data were assigned an initial count of 10, and these proteins are the majority of proteins with scaled abundance was greater than 2.4. While it is not clear whether multiple rounds of the cell cycle would narrow the distribution, improvement in the proteomics annotations of protein subunits in the membrane would improve the analysis. The exact values of scaled protein abundances are in File S2.

In the current translation and translocation network, the failure of cotranslational translocation of membrane proteins was considered to lead to the degradation of membrane proteins by FtsH, the membrane embedded protease. The predicted failure rate of targeting transmembrane proteins ranged from 4% to 8% with median 6.9%, and for lipoproteins, the rates were lower with a median of 3.5%, consistent with an experimental observation.^91^ The loss due to degradation slightly undermine the doubling of membrane proteins. The degradation of transmembrane proteins by FtsH to recycle free amino acids was minor compared to amino acid import via passive permeases and active ABC transporter, Opp in Figure S8(a,b). However, the analysis of amino acids’ economy and doubling of proteome could be improved by the degradation of cytoplasmic proteins by another protease, encoded by *Lon*/394.^92^

From the translocation network, we could also further differentiate the occupancy states of ribosome in Figure 3(f). The active ratio of ribosome at steady-state was 73% out of total on the population average as stated before. The active ribosomes are directed to the plasma membrane by the targeting factors SRP/SR and SecA for transmembrane proteins and secretory proteins, respectively. The ratio of ribosomes in SRP/SR/Sec and SecA/Sec translocation pathways over the active ribosomes was 19% 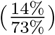 at steady state. This ratio of 20% was closely related to the summed mRNA counts of the transmembrane protein and secretory protein mRNAs over the total summed mRNA counts that could be roughly estimated by the ratio of genes encoding transmembrane proteins and secretory proteins, which is 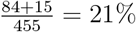. As a subset of ribosomes in two translocation pathways, the predicted ratio of the ribosome that resides in the Sec translocon was lower at 9% out of the total ribosomes. Based on cryo-ET images,^9^ we were also able to estimate this ratio by counting the ribosomes that were located 11 nm from the cell membrane. 11 nm, as half the ribosome size of 21 nm^93^ was selected due to the very close contact between the ribosome and the Sec translocon. ^94^ Two cryo-ETs gave ratios 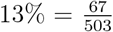 and 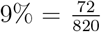, which supported our simulation. Of the involved translocation entities, the Sec translocon was the most active with a ratio of 74% due to the small number of Sec translocons compared to the ribosomes for transmembrane proteins and secretory proteins. Together, these assessments—including the predicted fraction of ribosomes bound to the Sec translocon and the measured failure rate of co-translational translocation, both consistent with experimental observations—support the accuracy of the translocation network at the whole-cell scale.

### Ribosome biogenesis under the constraint of subunit synthesis with the reduced SSU assembly pathway

Downstream of gene expression, the formation of macromolecular complex is constrained by subunit synthesis where proteins are translated from mRNA and three rRNAs in ribosomes are the direct transcription products. Assembly intermediates along pathways can accumulate and reduce assembly efficiency for two main reasons in the context of the time-dependent whole-cell model when no kinetic trapped intermediates considered. First, bottleneck subunits—those with the lowest available counts—limit the maximal possible amount of assembled complex and lead to the buildup of upstream intermediates. Second, the assembly process entails waiting periods necessary for the sequential binding of subunits to intermediates via bimolecular reactions. To quantify the impact of the second reason, we calculated the unassembled accumulation by subtracting the count of assembled holocomplexes from that of the bottleneck subunit. This value represents intermediates that are capable of progressing to fully assembled complexes but remain incomplete due to delays in bimolecular binding.

Ribosome biogenesis from 3 rRNAs and approximately 50 ribosome proteins using the reduced and sequential SSU assembly pathway in Figure 1(a) nearly duplicated the initial 500 ribosomes, the largest macromolecular complex in Syn3A with a median of 400 assembled at the end of the cell cycle (distribution in Figure S7(f)). Ribosome formation proceeded rapidly, with approximately 400 ribosomes assembled within 100 minutes. This high rate was driven by the substantial allocation of cellular resources to the production of ribosomal components. Specifically, transcription of rRNAs and mRNAs encoding ribosomal proteins accounted for 26% of the total transcriptional cost (measured by RNA nucleotide synthesis), while translation of ribosomal proteins consumed 14% of the total amino acid synthesis. SSU and LSU were assembled at consistent paces with nearly parallel trajectories shown in Figure 4(a). The monotonic increasing of assembled SSU, LSU, and then ribosome came from the continuing synthesis of rRNAs and ribosomal proteins as gene expression products during the cell cycle.

**Figure 4.**
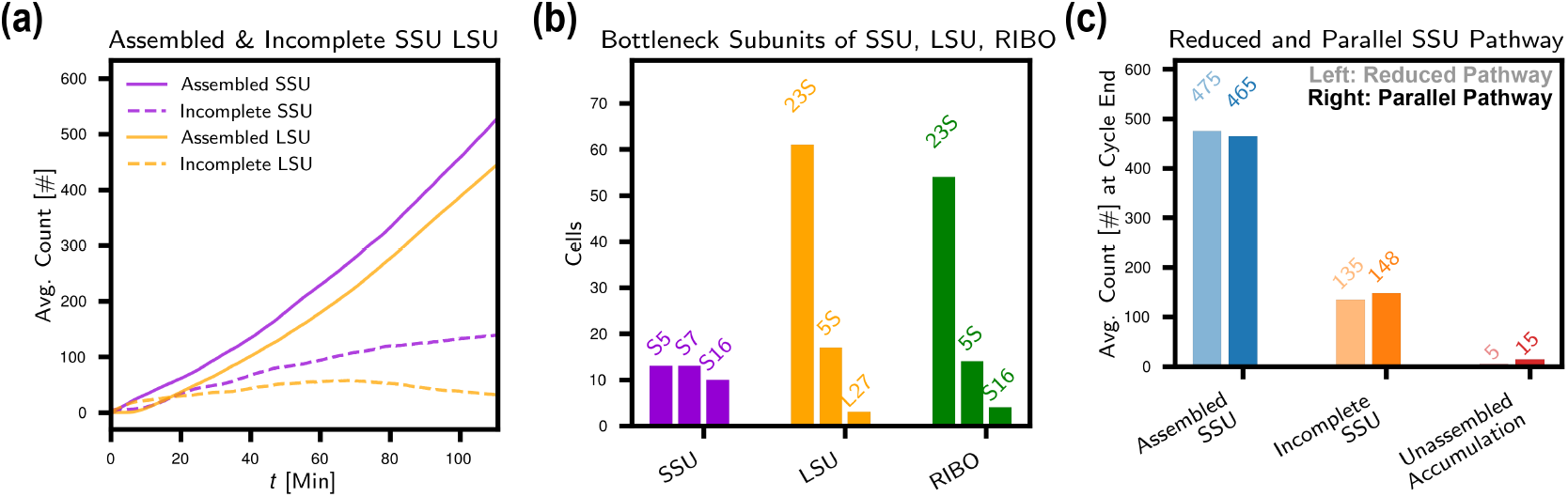
Ribosome biogenesis under the constraint of subunit synthesis with the reduced SSU assembly pathway. (a) Assembled and incomplete ribosome SSU and LSU with the reduced SSU assembly pathway. (b) Top three bottleneck subunits for ribosome SSU, ribosome LSU, and ribosome at the end of cell cycle. (c) Assembled SSU, incomplete SSU, and unassembled accumulation of SSU at the end of cell cycle with reduced or parallel SSU assembly pathways.

Since the formation of SSU was faster than LSU, a surplus up to 80 of SSU occurred at the end of the cell cycle. To reason the differences of assembled SSU and LSU, the bottleneck subunits of SSU, LSU, and the entire ribosome were identified in Figure 4(b). For SSU, ribosomal proteins S5, S17, and S16 mainly limits the assembly of SSU. For LSU, 23S rRNA (in more than half of the cells) as the starting point of assembly, limits the formation of LSU. The next bottleneck subunit is 5S rRNA, which associates with the LSU as the 17th of 32 total binders, and up to 10 copies of the intermediate that binds to 5S rRNA were observed to accumulate. The bottleneck subunits for LSU and ribosome were nearly identical, disclosing that LSU constrained the formation of ribosome, which explains the net accumulation of SSU.

By the end of the cell cycle, approximately 100 SSU intermediates and 40 LSU intermediates accumulate (Fig. 4a). Assembly intermediates were calculated as the summation of intermediate counts from bare rRNA to the last intermediate. Both the bottleneck subunits that stall assembly midway and the waiting times required for sequential bimolecular binding events can contribute to the accumulation under the approximation of no kinetic traps. For SSU, the bottleneck effect came from ribosomal proteins, which means that the assembly of SSU was stuck in the middle more due to the lack of proteins compared to excess 16S rRNAs, and the average abundance of total SSU intermediates were up to 130. However, the quantified unassembled accumulation was only 5 for SSU assembly with the reduced pathway, indicating minor delay from the association waiting times. In contrast to SSU, the synthesis of 23S rRNA, the starting point of assembly dominantly restricts the LSU assembly, thus the incomplete LSU was less trapped. Meanwhile, due to the longer assembly pathway, unassembled accumulation played a larger role for LSU and occupied 20 out of the 40 accumulation of intermediates. As another support of this, the assembled LSU started to emerge around 5 to 10 min, later than the SSU around 2 min since 31 LSU ribosome proteins and 5S rRNA must bind to 23S rRNA, while only 20 SSU ribosome proteins bind to 16S rRNA. These incomplete ribosomes are not kinetically trapped species; in subsequent generations, they provide productive starting points for ribosome assembly in daughter cells.

To investigate the impact of reducing the SSU assembly pathway from a parallel network of 145 intermediates to a nearly linear sequence of 19 intermediates, we compared the incomplete SSU and unassembled accumulation with two landscapes, as shown in Figure 4(c). (A separate set of 50 cell replicates was sampled.) When transitioning from the highly parallel to the reduced pathway, the number of SSU intermediates decreased from 148 to 135, and the unassembled accumulation dropped from 15 to 5. The underlying cause was the elimination of intermediates with slow binding kinetics when enforcing the fastest linear pathway. Biologically, these kinetically less favored pathways also contribute to the assembled ribosome, and more importantly provide additional redundancy when the fastest pathway was temporarily not accessible under stress conditions—such as rRNA misfolding.^15^ Over the course of the entire cell cycle, the total number of assembled SSUs reached 500, making the difference minor, and the reduction a reasonable approximation.

Our current model emphasized the basic principles of synthetic balances and kinetic associations of rRNAs and ribosomal proteins. Equally important in the orchestrated ribosome biogenesis but not explicitly simulated in our model is the involvement of accessory proteins that listed in Table S4 in File S1 and explained nearby. rRNAs as the long scaffolds for assembly overcome the local nonnative bottleneck with the assistance of GTPase, assembly factors (ribosome dedicated chaperons), and ribosomal proteins that reshaping the folding landscapes.^95^ Once misfolding, rRNA helicases resolve the kinetic trapped states to continue the efficient assembly.^96^ Early *in vitro* reconstruction study showed that GTPase Era, and assembly factors RimM and RimP enhanced the binding kinetics of SSU ribosomal proteins at different stages to several folds,^97^ with more recent knockout experiment of Era demonstrating the accumulation of immature SSU in *E. coli*.^98^ In gram-positive *B. subtilis*, LSU GTPase RbgA has been proved to be critical to unlock the intermediate state for maturing rRNA helices in the A, P and E functional sites.^99^ In this model, we based the kinetics association of rRNAs and ribosomal proteins on *in vitro* experiments that without assembly factors,^15,24^ and did not consider their altering effects on association kinetics between rRNAs and ribosomal proteins. Future incorporation of these factors could improve the speed and efficiency of ribosome biogenesis.

### Dependence of membrane complex assembly upon the 2D association rates

The assembly rates of many protein complexes remain poorly characterized, particularly for membrane-associated complexes. Furthermore, it is unclear how these assembly rates influence the overall rate of complex formation and the abundance of fully assembled structures, especially given the time scales associated with subunit synthesis. To investigate this relationship, we varied the bimolecular association rates on membrane 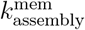 from 2.5 ×10^−5^ to 2.5 ×10^−4^, and 2.5 ×10^−3^ *µ*m^2^s^−1^ and uncoupled the influence from gene expression by defining the unassembled accumulation of assembly intermediates. The fastest 2.5 ×10^−3^ *µ*m^2^s^−1^ is close to the 2D rate measured by Huang *et al*.^40^

Here, we used the assembly of rnsBACD, the nucleoside ABC transporter as an example. Before the assembly of subunits, the translocation of subunits of membrane complexes occurs through the pathways in Figure 2. The two transmembrane subunits RnsD/0008 and RnsC/0009 as TMDs of rnsBACD are translocated via the SRP/SR/Sec pathway. The peripheral membrane ATP binding subunit RnsA/0010 as NBD is translated in the cytoplasm, and the substrate binding domain (SBP), lipoprotein RnsB/0011, is translocated via SecA/Sec pathway then undergoes modification by Lgt and Lsp. Upon translation and translocation of four subunits, the assembly assumably starts with two TMDs (RnsD/0008 and RnsC/0009) binding to form the permease channel. Then, the ATP binding domains (RnsA/0010) and substrate binding protein (SBP, RnsB/0011) associate with the channel. The initial count of the rnsBACD holocomplex was set to 145, same as the proteomics abundance of the peripheral membrane protein RnsA/0010. The initial counts of free RnsD, RnsC, RnsA, and RnsB are 0, 30, 0, and 0 respectively.

The unassembled accumulation of rnsBACD intermediates increased significantly with the slower assembly rate as shown in Figure 5(a). At the higher assembly rate of 2.5 ×10^−3^ *µ*m^2^s^−1^, unassembled accumulation was few and minor during the cell cycle, and the highest count of the holocomplex rnsBACD (109 of 145, 75% of the initial count) were assembled at the end of the cell cycle in Figure 5(b). Subunit RnsB/0011 late in the assembly pathway had a increase with decreasing assembly rate as to be expected in Figure 5(c). The variation in the numbers of the different intermediates and subunits demonstrates the complexity of the assembly landscapes.

**Figure 5.**
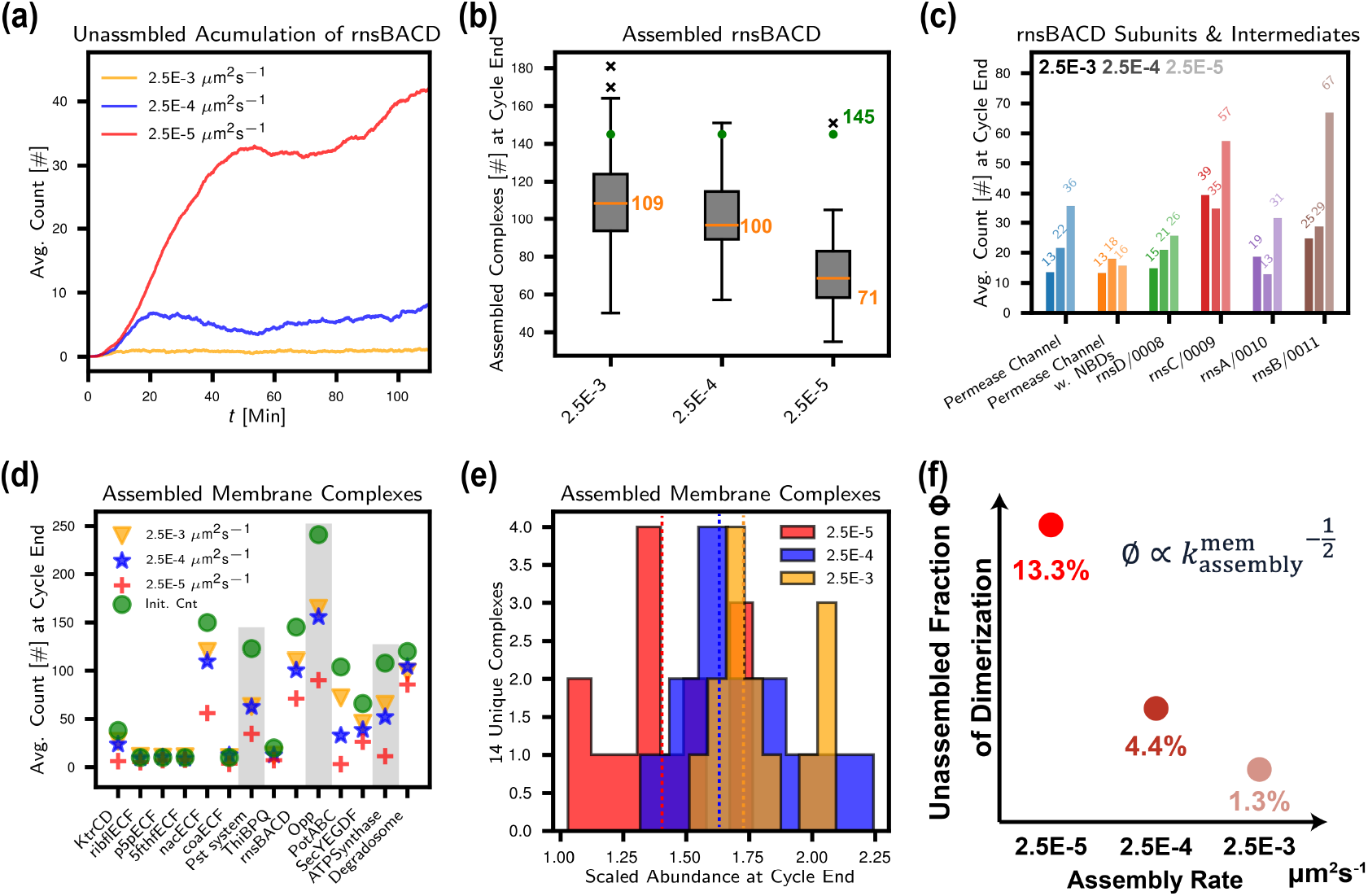
Assembly rate needs to match up with subunits synthesis speed from gene expression. (a) Unassembled accumulation of rnsBACD assembly under 2.5 × 10^−3^, 2.5 × 10^−4^, and 2.5 × 10^−5^ *µ*m^2^s^−1^ 2D association rate. (b) Distribution of assembled rnsBACD among population under three 2D rates. Initial count of complex is 145 shown as green dots. (c) Counts of rnsBACD assembly intermediates and subunits under three 2D rates at the end of the cell cycle. Left to right: 2.5 × 10^−3^, 2.5 × 10^−4^, and 2.5 × 10^−5^ *µ*m^2^s^−1^. (d) Comparison of population averaged assembled membrane complexes under three 2D rates. (e) Distribution of population averaged scaled assembled membrane complexes under three 2D rates with median of 1.73, 1.67, and 1.37, that marked by vertical lines respectively. (f) Unassembled fraction scales proportional to the reverse of root of assembly rate in a one step dimerization reaction hooked with gene expression of two subunits.

The correlation between decreased assembly yield and slower assembly rate occurred for most of the 14 membrane complexes, especially for ATP synthase with long assembly pathway in Figure 5(d). The fact that the scaled abundance for none of the holocomplexes was doubled at the end of the cell cycle, emphasizes the need for the fast association rate of membrane proteins. This result can be rationalized by considering the formation of a dimer from one-step association reaction given the kinetics for the coupled transcription, translation, and mRNA degradation in gene expression. The evaluated unassembled fraction *ϕ*, defined as the ratio of unassembled protein over the total synthesized protein at the end of the cell cycle, *t*_*cycle*_ was in Equation 4. *N*_*A*_ is the Avogadro constant, *V*_*cell*_ the cell volume, *SA*_*cell*_ the surface area, *N*_*Ava,RNAP*_, *N*_*Ava,Ribo*_, and *N*_*Ava,Degra*_ the count of free/available RNAP, ribosome, and degradosome, *N*_*gene*_ the count of the protein-coding gene, 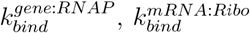, and 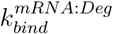 the binding rate in transcription, translation and degradation reaction, *k*_*assembly*_ is the protein assembly rate. *ϕ* increased from only 1.3% to 13% when *k*_*assembly*_ slowed from 2.5 ×10^−3^ to 2.5 ×10^−5^ *µ*m^2^s^−1^. The decreased efficiency will propagate along the assembly pathways and further suppress the yield of the final holocomplex, especially for long pathways, such as ATP synthase in Figure 5(d). Here, using the quantified unassembled fraction, we argue that the binding rate should be equal or larger than 2.5 ×10^−3^ *µ*m^2^s^−1^ to alleviate the unassembled accumulation of assembly intermediates in the context of the coupled subunit synthesis from gene expression.

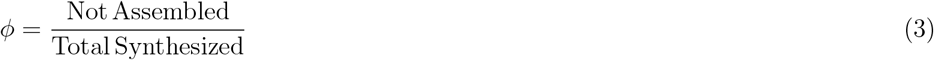

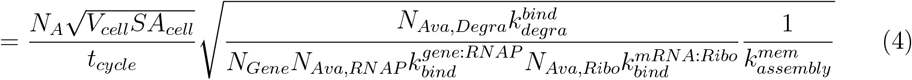

With the present kinetic parameters for genetic information processing, the abundances of the membrane complexes do not fully double over the cell cycle even with the fastest assembly rate in Figure 5(e). The primary reason is the constraint imposed by insufficient—and more importantly, non-stoichiometric—subunit synthesis from gene expression, as further discussed for ATP synthase in the following section.

### Assembly of ATP synthase highlights the co-expression of subunits and the alleviation of kinetic traps

An important example for the stoichiometric balance of subunits, ATP synthase has eight distinct subunits that possess different stoichiometries of 1 (subunit *ϵ, γ, δ*, and *a*), 2 (subunit *b*), 3 (subunit *β*, and *α*), or 10 (subunit *c*)^100^ to a complete structure of *α*_3_*β*_3_*γϵδc*_10_*ab*_2_. In our simulations, the initial count of intact ATP synthase was given as the number of peripheral F_1_ subunit *ϵ* (108) considering the stoichiometric ratio among the F_1_ sector subunits *α, β, γ*, and *ϵ* (See Table S2). To approximate the stoichiometric balance of subunit synthesis, our current implementation differentiated promoter strength at transcription level by linearly scaling the promoter strength with the subunit abundances. With this strategy, the synthesized mRNA (relative to *ϵ*/0789) has a rather linear relationship with the stoichiometries (*ϵ* 1/1, *β* 3/2.7, *γ* 1/1.1, *α* 3/2.5, *δ* 1/1, *b* 2/1.9, *c* 10/6.8, and *a* 1/0.87). However, experimentally obtained mRNA abundances show a larger deviation from the stoichiometries for subunit *δ, b, c*, and *a* as 1/0.5, 2/1.4, 10/4.3, and 1/1.9, respectively. The experimental abundances were derived from short-read RNA sequencing experiments.^4^ A similar deviation has also been observed in long-read RNA sequencing experiments in a synthetic relative to Syn3A, Syn1.0 (data unpublished). The inconsistency between the mRNA abundances and the stoichiometries for our WCM simulations and those observed experimentally is attributed to our implementation of gene expression in the model. mRNA processing, mRNA stability, differential transcription, and differential translation have been well characterized to impact mRNA (and protein) expression levels,^101^ however without the experimental detail to capture these processes accurately we simplified and assumed a linear link (proxy promoter) between protein and mRNA expression to ensure stoichiometric expression within complexes.

In general the stoichiometric balance of the eight subunits was roughly maintained on average as shown in Figure 6(a), where the synthesized subunits over their corresponding stoichiometries ranged from 100 to 130 at the end of the cell cycle. However, 65 intact ATP synthase (60% of the initial 108) on average were assembled at the end of the cell cycle, significantly less than the median of the synthesized subunits. We reason the inconsistency between synthesized subunits and assembled complete ATP synthase from two aspects. First, the strong heterogeneity of subunit synthesis at single-cell level exists and undermines the efficiency of assembly by the non-stoichiometric expression of the eight subunits. The quantified coefficient of variances (CVs) of synthesized subunits over the stoichiometries for each cell replicate were moderate with a median of 0.3. Thus, the surplus of certain precursors and the accumulation of intermediates occurred differently in each cell but could not be assembled into the holocomplex, then causing the waste of assembly building blocks. At the end of the cell cycle, the highest possible counts of ATP synthase (i.e. the lowest count of subunits over stoichiometries) had a mean of 75 and maximum of 102. This subtle point highlights the importance of co-expression of proteins by the polycistronic transcript from the ATP synthase operon.

**Figure 6.**
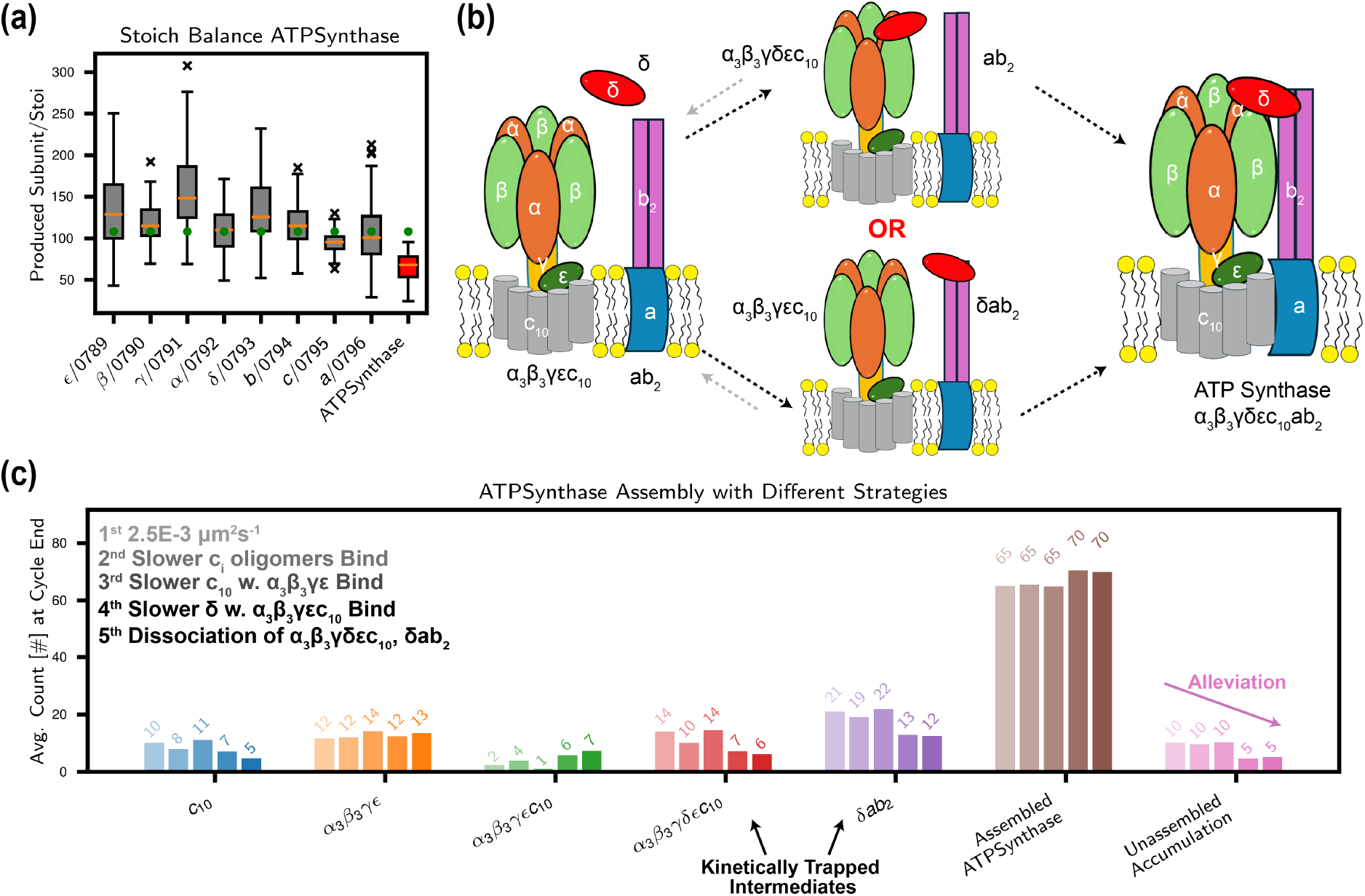
Assembly of ATP synthase with kinetic traps considered. (a) Distribution of synthesized ATP synthase’s eight subunits over stoichiometries at the end of cell cycle, and assembled ATP synthase. The green dots denote the initial 108 complete ATP synthases, and the orange lines show the median. (b) The last stage association of *ab*_2_ and *α*_3_*β*_3_*γϵδc*_10_ to ATP synthase occurs via the glue of *δ* subunit. However, no direct evidence was reported on the exact mechanism of this process. (c) Population averaged counts of intermediates that not incorporated into ATP synthase, assembled complete ATP synthase, and unassembled accumulation under five different assembly landscapes at the end of the cell cycle. Counts of assembled ATP synthase were corrected by assuming the same highest possible counts of ATP synthase of 75 as in the uniform rates.

Complexity of the assembly pathway seen in this study is the second factor. The peripheral F_1_ module *α*_3_*β*_3_*γϵ* was incorporated into the membrane by forming the complex *α*_3_*β*_3_*γϵc*_10_ with the *c*_10_ ring. Finally, the F_0_ module *ab*_2_ bind with *α*_3_*β*_3_*γϵc*_10_ by the gluing of peripheral subunit *δ* in Figure 6(b). With binding rates all set to 2.5 ×10^−3^ *µ*m^2^s^−1^, the pair of kinetically trapped assembly intermediates *α*_3_*β*_3_*γϵδc*_10_ and *δab*_2_ accumulated up to 14, undermining the assembly efficiency of holo ATP synthase. We probe the modular assembly landscape of ATP synthase^42^ by changing the binding rates between certain intermediates or adding reverse dissociation reactions to see the impacts (Detailed reaction schemes in Table S4). For alternative assembly landscapes beyond the uniform rate of 2.5 ×10^−3^ *µ*m^2^s^−1^, 50 separate cell replicates were simulated for each case to capture statistical variation.

In Figure 6(c), slowing the binding rates between the *c* subunit oligomers, *c*_*i*_, (*i* = 2 … 9), increases the accumulation of oligomers, but cannot increase the formation of complete ATP synthase significantly, since the *c*_10_ ring was cumulated in both cases. We tested three different strategies by differentiating the assembly rates or reactions to eliminate the non-functional structures, the pair of kinetically trapped intermediates *α*_3_*β*_3_*γϵδc*_10_ and *δab*_2_. Decreasing the binding rate between the *α*_3_*β*_3_*γϵ* and the *c*_10_-ring by one order of magnitude slightly suppressed the formation of *α*_3_*β*_3_*γϵc*_10_, yet not decreased the accumulation of the trapped complex *α*_3_*β*_3_*γϵδc*_10_, due to the fast binding of *δ* with *α*_3_*β*_3_*γϵc*_10_. Discriminating the association of the glue *δ* subunit with *α*_3_*β*_3_*γϵc*_10_ by the same one order of magnitude further increased the final generation of the complex to 70 (65% of the initial 108). Adding the dissociation reactions of two trapped complexes while making the dissociation of the unstable *α*_3_*β*_3_*γϵc*_10_ intermediate faster 10^−2^ s^−1^ could also upgrade the generation of the final ATP synthase up to 70. As further evidence, the unassembled accumulation of the assembly intermediates also decreased with the alleviation of kinetic traps in all three strategies. ATP synthase contains 21 subunits of eight unique types, creating a vast parameter space for potential assembly kinetics. In this study, we performed only minimal optimization beyond assigning uniform association rates, improving assembly efficiency primarily by introducing hierarchical association kinetics to alleviate kinetic traps. Similar strategies could operate *in vivo* to enhance modular assembly. More generally, the principle of identifying optimal kinetic protocols within a high-dimensional parameter space aligns with recent work using automatic differentiation to systematically explore association rates,^31^ which demonstrated that appropriate kinetic hierarchies can robustly avoid trapping. Such optimization strategies could, in principle, be extended to the assembly of other complex biological machines, including ATP synthase.

### Analysis of time-dependent cell metabolomics

We trained an autoencoder neural network to compress the time-dependent concentrations of 148 intracellular metabolites into a 16-dimensional latent representation. Trajectory inference applied to this lower-dimensional space identified three distinct cell lineages—lineages 1, 2, and 3—comprising 46, 17, and 37 cells, respectively. In the following, we refer to the cell lineage as a cell population. Metabolic phenotype divergence occurred at approximately 10 minutes for population 3 relative to populations 1 and 2, and at around 26 minutes for populations 1 and 2, corresponding to the early stage of the cell cycle. Key metabolites and reactions associated with the divergence were determined using two-sample *t*-tests, and the magnitudes of these differences using standard effect sizes (Cohen’s d statistic) are shown in section Machine learning on metabolomics to identify three phenotypes and Figure S9 in File S1.

To probe the effect of introducing the complex assembly into the whole-cell model, we conducted a control simulation based on the previous model where only representative subunits were used to measure the abundances of the functional protein complexes (The representative subunits are listed as Table S7 in File S1). A statistically significant difference was found in the metabolite concentrations and fluxes between the simulations. Higher concentrations of (deoxy)nucleotides in Figure 7(a) were produced in the control simulation, due to the higher fluxes in glycolysis and nucleotide metabolism shown in Figure 7(b). The underlying reason for such difference was probably the increased counts of ABC transporters for phosphate ion and nucleosides in Figure 7(c) that enhanced their import. In addition, more PEP (phosphoenolpyruvic acid) was generated in Figure 7(d), thus increasing the import of glucose in the phosphorelay system. In the metabolomics analysis of the simulation with complexes, a much higher population of cells had a lower concentration of ATP in better agreement with the experimentally reported value of 4.4 mM.^102^ Finally, we conclude that the incorporation of complex assembly prevents the overestimation of membrane transporters, leading to elevated import fluxes of nutrients and more generation of energetic metabolites. See section Comparative analysis on metabolomics between with and without complex assembly in File S1 for detailed algorithms.

**Figure 7.**
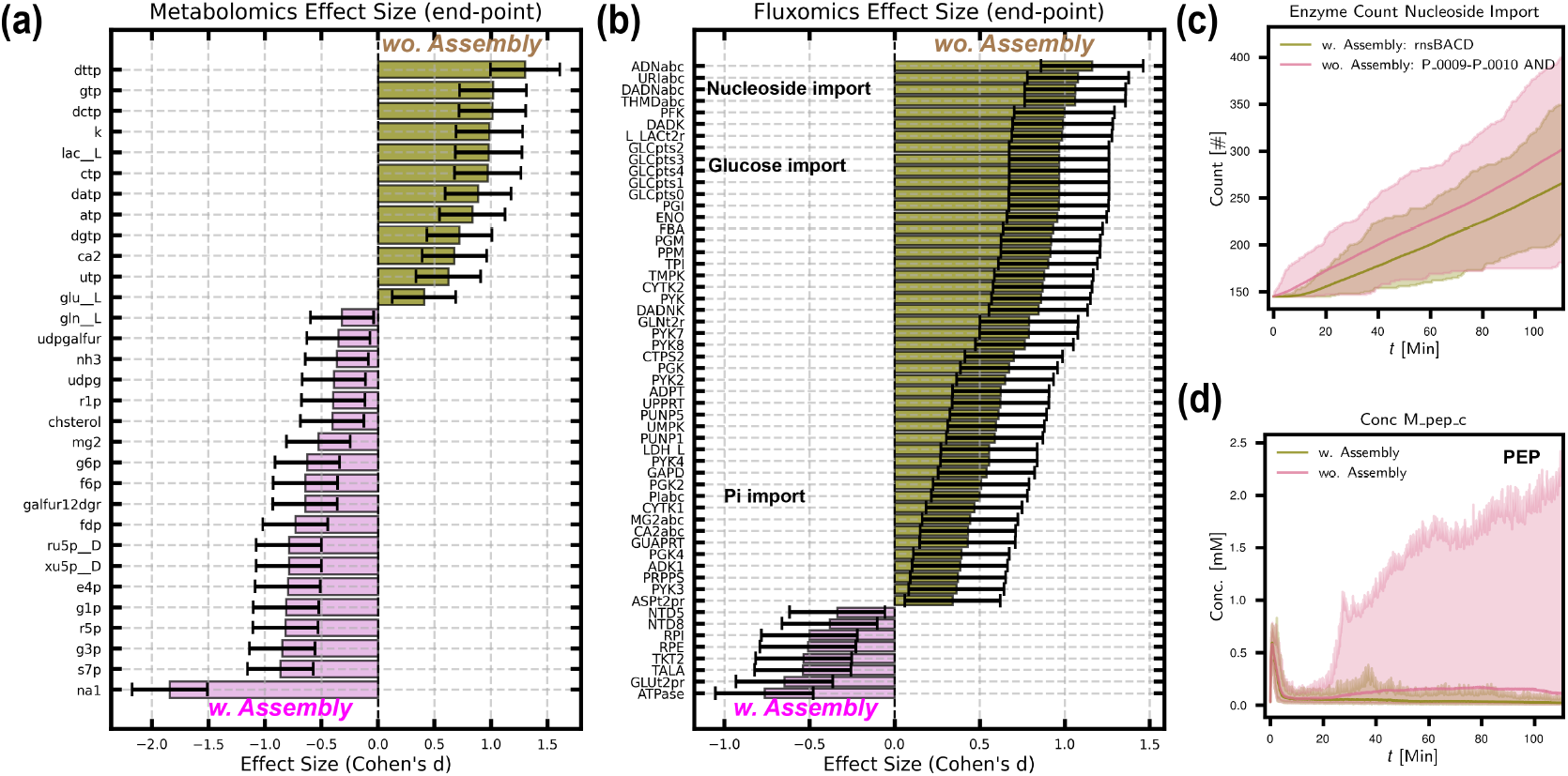
Comparison of metabolites and fluxes between simulations with and without complex assembly. (a) Differential metabolite concentrations at the end of cell cycle between simulations with (pink) and without (olive) complex assembly. (b) Differential reaction fluxes between simulations with (pink) and without (olive) complex assembly. The effect size shading indicates which simulation has the larger quantity of the metabolite or reaction flux. The black bars show the 95% confidence interval of the effect size, and the absence of zero indicates the difference are statistically significant. (c) Enzyme counts for nucleoside import were higher without complex assembly since only two subunits were used to represent the complete complex. (d) PEP concentration were much higher in simulation without complex assembly, thus accelerating the import of glucose in phosphorelay system.

We can only more completely identified which stochastic events in gene expression and complex formation lead to the metabolic phenotypes once our ongoing work to develop a multi-omics encoder-decoder architecture is completed.

## Conclusions

In the context of the well-stirred whole-cell model of the minimal cell, we showed in Figure 5 that the efficiency of membrane complex assembly can be significantly hindered by the reduction in the 2D association rate from 2.5 ×10^−3^ to 2.5 ×10^−5^ *µ*m^2^s^−1^. This highlights the temporal competition of gene expression and complex assembly at the whole-cell models.

The non-stoichiometric synthesis of protein subunits at the single-cell level was seen for the assembly ATP synthase, which contains both transmembrane F_0_ and peripheral membrane F_1_ components arising from eight genes in a polycistronic operon. Each subunits have distinct stoichiometries in the complex. This result underscores the need to incorporate co-expression from polycistronic mRNA operon into the transcription model.

Although lacking spatial heterogeneity, the well-stirred model successfully predicted key dynamics at the whole-cell scale. The simulated cell cycle durations—defined as the time required to double the cell surface area—ranged from 96 to 110 minutes, in agreement with experimental measurements.^2–4^ The total proteome approximately doubled by the end of the cell cycle. We quantified mRNA translation efficiency based on mRNA binding rates and the availability of ribosomes and degradosomes, which is influenced by their macromolecular complex assembly. The integrated whole-cell translation and translocation networks enabled us to estimate that approximately 9% of ribosomes were associated with the Sec translocon, consistent with experimental observations of a number of ribosomes near the cell membrane in Syn3A.^6^

Furthermore, a machine learning learning of the time-dependent metabolic profiles revealed the emergence of three distinct cell populations. More importantly, it emphasized that differences of metabolite concentrations and reaction fluxes throughout the metabolic network with and without additional complex assembly. The simulation with complex assembly gave better agreement to the ATP concentration reported in the recent review over all domains of life.^102^

### Limitation and Outlook

The simulations discussed were performed under the well-stirred assumption (CME and ODE), which ignores the spatial heterogeneity of bacterial cells. Therefore, processes such as the diffusion of DNA, RNA, and proteins are not considered, and their respective impacts on cellular dynamics are absent from our model. The differences resulting from the spatially homogeneous assumption are evident when the WCM model presented here is compared to a recently reported spatial WCM model (4DWCM).^86^ For example, DNA replication initiation within the spatially homogeneous simulations happens roughly twice as fast as in the 4DWCM. The difference in cellular behaviors results from diffusion that affects the binding of the protein that initiates DNA replication, DnaA. While the well-stirred model does treat the formation of membrane complexes constrained to 2D, spatial heterogeneity and diffusion will certainly play a bigger role in the assembly for larger bacterial and eukaryotic cells. The main advantage in the well-stirred model is the ability to test a wide range of parameters in tens to hundreds of replicate cells, compared to the computationally expensive 4D WCM.^86^

Our model provides a unique ability to capture the dynamics of many cellular processes simultaneously, predicting more quantities than can be measured in any single experiment, however our representation of bacterial life is incomplete. In our simulations, there are missing bacterial processes that are understood to be important in the stoichiometric expression of proteins,^101^ and the rate of ribosome assembly by accessory factors. Replacing the independent expression with the co-expression of polycistronic operons should improve the complex assembly efficiency, as demonstrated in a WCM of *E. coli*.^103^ Currently, efforts are underway to characterize the mRNA isoforms associated with polycistronic operons using long-read RNA sequencing data. With this data, the gene expression model could be refined to recapitulate the transcriptome (and thus the proteome) as has been experimentally observed in Syn3A.

## Supporting information

Updated SI PDF

## Acknowledgement

E.F and Z.LS are partially supported by NSF MCB-2221237 and the NSF Science and Technology Center for Quantitative Cell Biology (NSF DBI-2243257), T.B and B.G are partially supported by NSF MCB-2221237. S.W., B.Y., and R.W are partially supported by NSF DMS-2515171 and the NSF Science and Technology Center for Quantitative Cell Biology (NSF DBI-2243257).

Z.R.T.: Research reported in this publication was supported by the Cancer Center at Illinois - Beckman Institute Postdoctoral Fellows Program sponsored by the Cancer Center at Illinois and the Beckman Institute for Advanced Science and Technology, University of Illinois Urbana-Champaign. The content is solely the responsibility of the authors and does not necessarily represent the official views of the program sponsors.

## Supporting Information Available

The simulation input and script is available on GitHub at https://github.com/Luthey-Schulten-Lab/Minimal_Cell_ComplexFormation.

The Supporting Information is available free of charge.

- File S1 Supporting Information as PDF. Metabolic networks with annotated protein complexes, reactions of gene expression model, compositions and assembly pathways of complexes, translocation network, derivation of unassembled fraction, SI Table S1-S12, SI Figure S1-S9.
- File S2 Proteome, initial mRNA counts, and initial metabolites concentrations as Excel file. Tab 1 Comparative Proteomics: Comparative proteomics of Syn3A to other with the revised annotations and localizations of Syn3A’s proteome; annotations of certain proteins updated^53^ noted by slash separation. Tab 2 Protein Topologies: Prediction of signal peptide and transmembrane regions from SignalP and DeepTMHMM. Tab 3 Scaled Proteome: Partitioning of the proteomics counts for complex subunits; the scaled abundances of all proteins at the end of cell cycle. Tab 4 mRNA Count: Initial average counts of mRNAs. Tab 5 Intracellular Metabolites: Initial concentrations of intracellular metabolites, and time-averaged concentrations of intracellular metabolites. Tab 6 Experimental Medium: Experimental Defined Medium. Tab 7 Simulation Medium: Simulation Defined Medium.
- File S3 kinetic parameters as Excel file. Tab 1 Central: Kinetic parameter for central metabolism. Tab 2 Nucleotide: Kinetic parameter for nucleotide metabolism. Tab 3 Lipid: Kinetic parameter for lipid metabolism. Tab 4 Cofactor: Kinetic parameter for cofactor metabolism. Tab 5 Transport: Kinetic parameter for metabolic transport reactions. Tab 6 Non-Random-Binding Reactions: Kinetic parameter for non-random binding reactions. Tab 7 tRNA charging: Kinetic parameter for tRNA charging. Tab 8 Gene Expression: Kinetic parameters for gene expression. Tab 9 SSU Assembly: Assembly reactions and rates for ribosome SSU. Tab 10 LSU Assembly: Assembly reactions and rates of ribosome LSU. Tab 11 LSU Assembly w. 4 L7s: Assembly reactions and rates of ribosome LSU with four L7/L12 proteins.
- File S4 ML on metabolomics as ZIP file. Machine learning analysis on metabolomics with the traces of differential metabolites and reactions for all 3 lineage pairs; Comparative analysis between two sets of metabolomics with and without complex assembly.
- File S5 Trajectories of all time-dependent metabolite concentrations and metabolic fluxes as PDF file. Solid lines represent the population averages. Shaded regions represent the ranges observed for the entire population of simulated cell.

## TOC Graphic

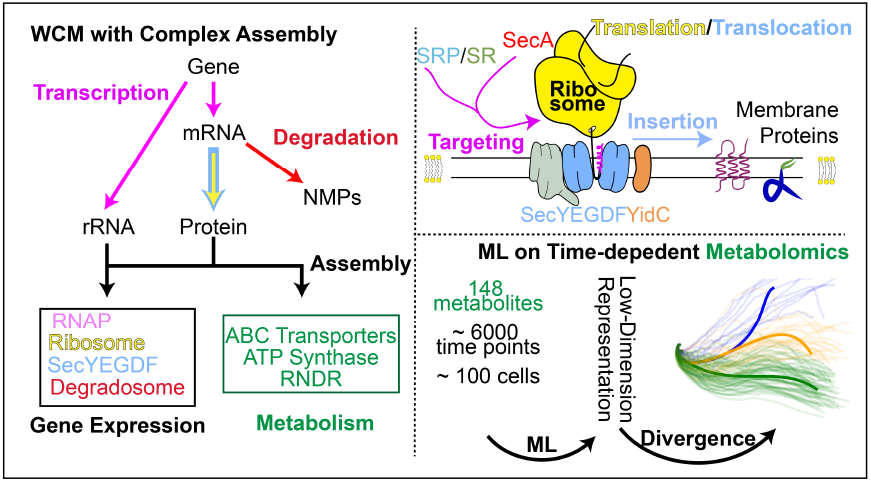

