## Supplementary material for "Assembly of Macromolecular Complexes in the Whole-Cell Model of a Minimal Cell": Updated SI PDF

### Contents


**1 Genetic Information Processes and Metabolism S3**

1.1 Metabolism . . . S3

1.1.1 Increased Rates in Phosphorelay and Central Metabolism . . . S4

1.1.2 Adjusted Rates in Nucleoside Import Reactions . . . S5

1.1.3 Gibbs Free Energy Analysis in Central Metabolism . . . S6

1.2 Genetic Information Process . . . S8

1.2.1 Replication of the circular chromosome . . . S8

1.2.2 Transcription of mRNA and tRNA, Translation and mRNA Degradation . . . S8

1.2.3 Transcription of rRNAs . . . S10

1.3 Communication between Genetic Information Process and Metabolism . . . S11

**2 Macromolecular Complexes in Syn3A and Assembly Pathways S13**

2.1 Compositions and Assembly Pathways . . . S13

2.2 RNA Polymerase . . . S16

2.3 Ribosome . . . S16

2.4 Chromosome Dynamics Related Complexes . . . S17

2.5 RNDR . . . S18

2.6 Degradosome . . . S18

2.7 Sec Translocon . . . S18

2.8 ATP Synthase . . . S19

2.9 ABC Importers . . . S21

2.10 ECF Transporters . . . S22

2.11 KtrCD, and Fak . . . S22

2.12 Representative Subunits of Macromolecular Complexes . . . S23

**3 Translocation Network in Syn3A S25**

3.1 Localization of 455 Proteins . . . S25

3.1.1 Two Extracellular Proteins, and Three Proteins with Unidentified Localization . . S25

3.1.2 Tail-anchored (TA) Transmembrane Proteins . . . S26

3.2 Translation and Translocation Reactions . . . S27

**4 Decreased Protein Assembly Rate Increases Unassembled Fraction S30**

**5 Supporting Figures S33**

**6 Comparative analysis on metabolomics between with and without complex assemblyS37**

6.1 Permutation-Based Two-Sample Test to Tell Statistically Significant Differences . . . S37

6.2 Differential Analysis of Metabolite Trajectories at End of the Cell Cycle . . . S37

6.3 Comparison of Cell Population Distribution of Metabolic Phenotypes Between Datasets . S37

### 1 Genetic Information Processes and Metabolism

#### 1.1 Metabolism

In lipid metabolism, fatty acid kinase catalyzes the phosphorylation of fatty acids, initiating the first step toward phospholipid synthesis. Coenzyme A, which is imported by the CoaECF transporter and also produced through central metabolism, donates the 4'-phosphopantetheine group to convert apo-ACP (apo–acyl carrier protein) into its active holo-ACP form.

In cofactor metabolism, the uptake of coenzyme A and four vitamin derivatives is mediated by five separate ECF transporters, all sharing the same ECF core module (0641/0642/0643), with the fourth S-component conferring substrate specificity. Spermine and thiamine diphosphate are imported via distinct ABC transporters.

For ion uptake, the Pst system, an ABC transporter, actively imports phosphate. Potassium ions are transported into the cytoplasm by KtrCD, which is activated by ATP and sodium ions. In the case of ATP synthase, proton flux is reversible and may result in either influx or efflux depending on substrate concentrations.

Free amino acids are taken up both actively by the Opp ABC transporter and passively through transmembrane permeases. Glutamate is imported via the permease GltP/0886, while the remaining 19 amino acids are imported through permeases 0876 and 0878, both of which have unknown substrate specificity (1).

In nucleotide metabolism, nucleosides are taken up by the nucleoside ABC transporter RnsBACD. Ribonucleotide diphosphate reductase (RNDR) irreversibly converts ribonucleoside diphosphates into deoxyribonucleoside diphosphates.

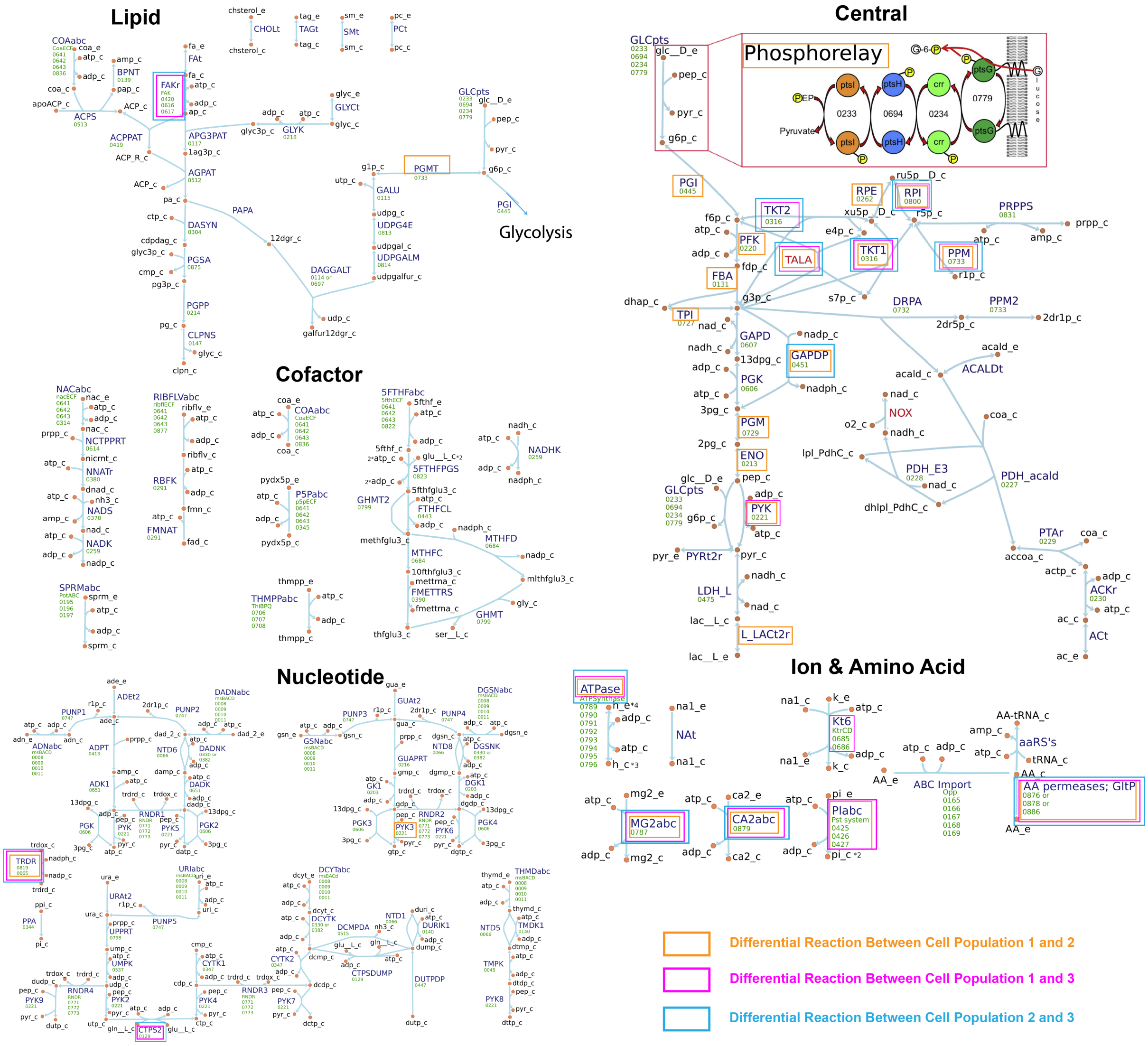


Figure S1: Essential metabolic networks of Syn3A. The bracketed reactions are the differential reactions identified for each pair of lineage/population. Reprinted with permission from Thornburg, Z. R.; Bianchi, D. M.; Brier, T. A.; Gilbert, B. R.; Earnest, T. M.;Melo, M. C.; Safronova, N.; Sáenz, J. P.; Cook, A. T.; Wise, K. S.; Hutchison, C. A.; Smith, H. O.; Glass, J. I.; Luthey-Schulten, Z. Fundamental behaviors emerge from simulations of a living minimal cell. *Cell* **2022**, *185*, 345–360.e28. Copyright 2022/CELL PRESS

##### 1.1.1 Increased Rates in Phosphorelay and Central Metabolism

As the first step in the phosphorelay system that import glucose, the rate of phosphotransfer from PEP to enzyme PtsI/0233 (NCBI Name: Phosphoenolpyruvate–protein phosphotransferase) $k_{f,GLCpts0}$ was measured by Roseman (2) in *E. coli*. Here, we increased the rate from $6{\times}10^6$ $\mathrm{M^{-1}s^{-1}}$ to $10{\times}10^6$ $\mathrm{M^{-1}s^{-1}}$, the maximal value reported.

The glucose uptake rate by phosphorated PtsG/0779 (NCBI Name: PTS sugar transporter) $k_{f,GLCpts4}$ was re-estimated from the $^{13}$C-glucose experiment conducted on *Mycoplasma pneumoniar* (3). In 2022, $k_{f,GLCpts4}$ was set to 0.88 $\mathrm{mM^{-1}\cdot s^{-1}}$. The PtsG count in *Mycoplasma pneumoniae* was reported between 251 and 431, and the glucose uptake rate per second between 18,000 and 24,000 with a media glucose concentration of 10 g/L. The lower limits of these two values were used. Then $k_{f,GLCpts4}$ was back calculated by the following equation.

$$k_{f,GLCpts4} = \frac{\text{per second glucose update flux}}{\text{conc. of ptsG} \times \text{conc. of glucose}} \tag{1}$$
$$= \frac{\frac{18000/s}{N_A V}}{\frac{251}{N_A V} \times \frac{10g/L}{180g/mol}} \tag{2}$$
$$= \frac{18000 \times 180}{251 \times 10}\ \mathrm{M}^{-1} \cdot \mathrm{s}^{-1} \tag{3}$$
$$= 1.29\ \mathrm{mM}^{-1} \cdot \mathrm{s}^{-1} \tag{4}$$

PEP serves a dual role: it is both a high-energy intermediate produced during glycolysis and the phosphate donor required for glucose import via the phosphorelay system. As reported in (4), a shortage of PEP can lead to ATP depletion. Since PEP is also consumed in phosphorelay system, we decided to upregulate the downstream flux of glycolysis to further increase PEP generation. To weaken the bottleneck effects of the FBA reaction (locusNum 0131, EC: 4.2.1.13) that prevent the downstream generation of PEP reported in (4), the largest forward rate $k_{f,FBA}$ 64.5 $s^{-1}$ reported in the BRENDA database (5) was used instead of the previous value of 21 $s^{-1}$. Following the same rationale, the forward rate constants $k_{f,\mathrm{PGK}_i}$ of PGK-catalyzed reactions involving different diphosphate nucleosides were increased by 75% of their previous values. The forward rate of the PGM-catalyzed reaction was doubled. To prevent excessive consumption of PEP in PYK-catalyzed reactions, the forward rate constants $k_{f,\mathrm{PYK}_i}$ were reduced to 66% of their previous values. The updated values all fall into the range of rates reported in BRENDA, and are available in File S3.

###### 1.1.2 Adjusted Rates in Nucleoside Import Reactions

The import rates of different (deoxy)nucleosides were changed on the basis of the estimation of the requirement of nucleotides in various cellular activities.

Imported nucleosides are further converted to nucleotides via different metabolic pathways. The sole fate of deoxynucleotides (dNTPs) is incorporated into chromosomes as monomers.

ATP is used in active import of nutrients by ABC and ECF transporters, converting nucleosides to nucleotides, chromosome replication, RNA transcription, and mRNA degradation, while GTP is used in a single translation process. Since ATP and GTP are converted to ADP and GDP, and adenosine and guanosine remain, these energetic processes put demands on glucose import rather than nucleoside import. CTP and UTP are also involved in lipid metabolism.

To estimate the $k_f$s in nucleoside import rate via rnsBACD ABC importer, a simple rate form that maximize the import rate was assumed:

$$\text{speed of import (/s)} = k_f\ \text{(/s)} \times \text{count of rnsBACD} \tag{5}$$

With notation $v_{in}$ and $n_{rnsBACD}(t)$:

$$v_{in} = k_f\ \text{(/s)} \times n_{rnsBACD}(t) \tag{6}$$

Here, a linear time-dependent number of rnsBACD transporter as assumed:

$$n_{rnsBACD}(t) = n(0)(1 + \frac{t}{T}) \tag{7}$$

where $n(0)$= 145 that was estimated initial count of rnsBACD based on counts of subunits P_0009 and P_0010, and $T$ is the cell cycle of roughly 100 minutes from the simulation.
So, in the time interval from 0 to t, the total number of imported nucleosides $n_{in}(t)$ is:

$$n_{in}(t) = \int_0^t k_f \times n_{rnsBACD}(t)dt \tag{8}$$
$$= k_f \int_0^t n(0)(1 + \frac{t}{T})dt \tag{9}$$
$$= k_f n(0)(\frac{1}{2T}t^2 + t) \tag{10}$$

For different nucleotides, $n_{in,nucleoside}(t)$ should be larger than $n_{cost,nucleoside}$, so that

$$k_f \geq \frac{n_{cost,nucleoside}}{n(0)(\frac{1}{2T}t^2 + t)} \tag{11}$$

The costs of nucleosides are listed:

- Doubling Chromosome in 1 hour
- Doubling mRNAs and tRNAs in 100 minutes
- Generate 500 of each 5S, 16S, and 23S rRNAs for ribosome biogenesis in 100 minutes

Due to the absence of an enzyme assigned that further converts cytidine to nucleotide (1), cytidine uptake was not simulated; instead CTP was generated primarily by conversion of UTP in the CTPS2 reaction. That was why the requirement for uridine was heaviest. The uptake of deoxyuridine was also not simulated, since it was not fed to the cell in the medium.

Based on the above information, we were able to calculate the minimal import rate of each nucleoside, shown in Table S1. We adopted $k_f$ of adenosine 1.5 $s^{-1}$, twice of the minimal requirement since ATP is heavily used in other energetic reactions; slightly lower guanosine import rate because GTP can also be generated by the conversion of deoxyguanosine by the DGSNK and PUNP4 reactions. The changed rates of nucleoside import are also available in File S3.

Deoxythymidine is bulkier than uridine due to its C5 methyl group, which increases steric hindrance and hydrophobicity, likely slowing its transport compared to the smaller, more polar uridine. Similarly, guanosine has additional functional groups at C2 and C6, making it bulkier than adenosine, which may hinder its interaction with nucleoside transporters and slow its import rate.

Table S1: Comparison of $k_f$ ($s^{-1}$) for nucleoside transport

| Nucleoside | Required | Current | Cell 2022 (4) |
|---|---|---|---|
| Adenosine | 0.74 | 1.5 | 2 |
| Guanosine | 0.63 | 0.5 | 1 |
| Uridine | 1.09 | 1.5 | 2 |
| Thymidine | 0.61 | 1 | 1 |
| Deoxyadenosine | 0.61 | 1 | 1.5 |
| Deoxyguanosine | 0.19 | 0.5 | 1 |
| Deoxycytidine | 0.19 | 0.5 | 0.5 |

##### 1.1.3 Gibbs Free Energy Analysis in Central Metabolism

To assess whether the central metabolic network still thermodynamically consistent after manipulating the parameters, the same Gibbs free energy change analysis was repeated and shown in Figure S2 as in Figure S6 in our previous publication (4). We calculate $\Delta G_{\text{kinetic}}$ of our metabolic network based on the updated set of kinetic parameters and the concentration of metabolites from the time-dependent simulations as Equations 12, 13, 14, where $\Delta G_{0,\text{kinetic}}$ is the standard Gibbs free energy, Q is the reaction quotient, $k_f$ and $k_r$ are the forward and backward rates, $K_M$ is the Michaelis-Menten constant.

$$\Delta G_{\text{kinetic}} = \Delta G_{0,\text{kinetic}} + RTlogQ \tag{12}$$

$$\Delta G_{0,\text{kinetic}} = -RTlog[\frac{k_f \prod_i^{products} K_{M,i}}{k_r \prod_j^{substractes} K_{M,j}}] \tag{13}$$

$$Q = \frac{\prod_i^{products}[Conc_i]}{\prod_j^{substrates}[Conc_j]} \tag{14}$$

First, we compare the updated Gibbs free energy values, $\Delta G_{\text{kinetic}}$ calculated from the above equations with the previous ones in 2022 (4) in Figure S2(a). A significant increase in the Gibbs free energy change ($\Delta G$) of the FBA reaction—from −30 kJ/mol to −20 kJ/mol, i.e., an increase of +10 kJ/mol—was observed. This change resulted from a decreased concentration of fructose-1,6-bisphosphate (FDP), the substrate of the FBA reaction, and increased concentrations of glyceraldehyde-3-phosphate (G3P) and dihydroxyacetone phosphate (DHAP), the products. This increase in $\Delta G$ was compensated by a corresponding decrease in the $\Delta G$ of the subsequent GAPD reaction, which dropped from 10 kJ/mol to 0 kJ/mol. As a result, the cumulative Gibbs free energy change along the pathway remained at approximately $-80$ kJ/mol, consistent with experimentally reported values in Figure S2(b).

We also calculated another set of $\Delta G_{0,\mathrm{thermo}}$ from equilibrium thermodynamics using Equilibrator, and computed $\Delta G_{\mathrm{thermo}} = \Delta G_{0,\mathrm{thermo}} + Q$ with the same reaction quotient. For reactions DRPA, FBA, and TPI, the deviations between $\Delta G_{\mathrm{thermo}}$ and $\Delta G_{\mathrm{kinetic}}$ were still large same as 2022, since we need to choose the parameter set to make these reaction to forward given the abundances of these three enzymes from proteomics (4).

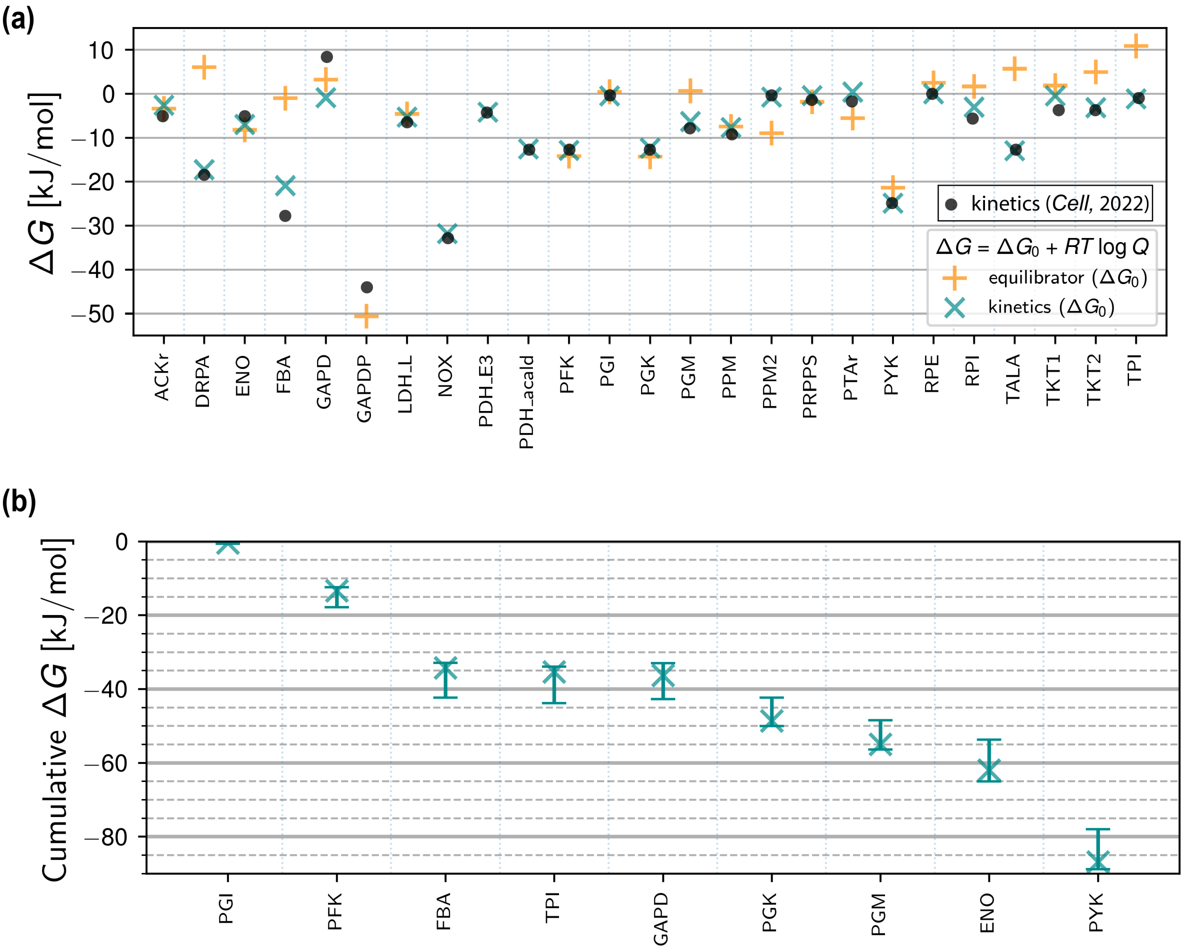


Figure S2: (a) Gibbs free energy change of reactions in central metabolism using reaction quotients calculated with time-averaged metabolite concentrations. pH was set to 7, and temperature as 303 K. Gibbs energy changes from kinetic parameters in 2022 (4) were annotated as black dots; from kinetic parameters in this work as darkcyan X; from thermodynamics in this work as orange cross. (b) Cumulative Gibbs free energy change along the Embden-Meyerhof-Parnas (EMP) glycolytic pathway of Syn3A calculated from kinetic parameters in this work.

#### 1.2 Genetic Information Process

The binding rates of RNAP to DNA genes, ribosome to mRNAs, and degradosome to mRNAs were estimated on the basis of the frequency of the binding events reported for *E. coli.* The stochastic binding reactions follow a simple bimolecular form (A+B to C). In the CME, the propensity $a_r$, which quantifies the probability per unit time that reaction $r$ will occur, is calculated for a second order binding reaction as $\frac{k_{\text{bind}} n_A n_B}{N_A V}$, where $k_{\text{bind}}$ is macroscopic binding rate reported in units of $\text{M}^{-1}\text{s}^{-1}$, $n_A$ and $n_B$ are the absolute counts of molecules A and B respectively, $N_A$ is Avogadro's constant, and $V$ is the volume. Over long times, the average number of reaction events per unit of time equals the time-averaged propensity, and we approximated the propensity by the observed frequency of binding events. Rearranging, we obtain the formula for the macroscopic binding rate $k_{\text{bind}}$:

$$k_{\text{bind}} = \frac{\text{frequency} \times N_A \times V}{n_A \times n_B} \tag{15}$$

##### 1.2.1 Replication of the circular chromosome

We start the simulation with one copy of the complete chromosome. Upon the replication initiation, two replisomes will bidirectionally duplicate the chromosome gene by gene until they meet near the *Ter*. The Hofmeyr rate form is also applied to the replication of each gene, where the entire gene sequence of promoter, coding region, and termination sequence, together with the intergenic region, were considered as $L_{polymer}$ to ensure that the entire circular genome was replicated. The replication speed is 100 bp/s, and dissociation constant of deoxynucleotides $K_{D,\text{dNTP}}$ is 1 $\mu$M.

##### 1.2.2 Transcription of mRNA and tRNA, Translation and mRNA Degradation

First, we describe the parameterization for transcription reactions, starting with the transcription of tRNA-coding genes. Due to the lack of measurements for binding of RNAP with tRNA-coding genes, we instead take the value measured for rRNA-coding genes. We use the values of 10 initiations of transcription of rRNA genes per minute and 11,400 RNAP molecules in a single *E. coli* as measured by Bremer et al. (6) The volume of a single *E. coli* cell is taken as 1fL, so the binding rate between tRNA genes with RNAP $k_{\text{bind}}^{\text{tRNAgenes:RNAP}}$ was calculated as

$$k_{\text{bind}}^{\text{tRNAgenes:RNAP}} = \frac{\frac{10}{60} \times V_{E.coli} \times N_A}{11400} = 8.8 \times 10^3\,\text{M}^{-1}\text{s}^{-1} \tag{16}$$

Thus, the binding between tRNA-coding genes with RNAP occurs via

$$\text{RNAP} + G_{\text{tRNA}} \rightarrow \text{RNAP} : G_{\text{tRNA}} + \sigma, \quad k_{\text{bind}}^{\text{tRNA\_genes:RNAP}} = 8.8 \times 10^3\,\text{M}^{-1}\text{s}^{-1} \tag{17}$$

where the sigma factor $\sigma$ dissociates from the core enzyme once transcription elongation starts.

The elongation speed of RNAP on tRNA-coding genes is taken as 85 nt/s, which is the value measured for RNAP on rRNA-coding genes in *E. coli* (6). The elongation occurs via

$$\text{RNAP} : G_{\text{tRNA}} \rightarrow \text{tRNA} + \text{RNAP} + G_{\text{tRNA}}, \quad v_{\text{polymerization}}^{\text{transcription,tRNA}} \tag{18}$$

$$v_{\text{polymerization}}^{\text{transcription,tRNA}} = \frac{85\,\text{nt/s}}{\frac{K_{D1,\text{NTP}_1} K_{D2,\text{NTP}_2}}{[\text{NTP}_1][\text{NTP}_2]} + \sum_i \frac{n_i K_{Di,\text{NTP}_i}}{[\text{NTP}_i]} + L_{\text{tRNA}} - 1} N_{\text{RNAP:G}_{\text{tRNA}}} \tag{19}$$

A promoter strength $S_{\text{promoter}}^{\text{locusNum}}$ was introduced as a prefactor at the transcription level to distinguish the expression strengths of the 455 different mRNAs. It was incorporated into both the binding and the elongation reactions to account for variations in both the RNAP binding affinity and transcription elongation speed. Promoter strengths were scaled according to the absolute protein counts, (4) using the formula.

$$S_{\text{promoter}}^{\text{locusNum}} = \frac{Init.Total\,Ptn.Cnt.}{180}, \tag{20}$$

where 180 is the average count of the 455 mRNA-coding proteins, and the initial protein count is the total initial protein count described in the section Methods - Abundances of Protein Complexes and Involved Protein Subunits.

The average binding rate of RNAP with mRNA-coding genes $k_{\text{bind}}^{\text{mRNAgenes:RNAP}}$ was scaled as

$$k_{\text{bind}}^{\text{mRNAgenes:RNAP}} = 8.8 \times 10^3 \times \frac{20}{85} = 2.1 \times 10^3\,\text{M}^{-1}\text{s}^{-1}, \tag{21}$$

since the average elongation speed of RNAP on protein encoding genes is taken as 20 nt/s. (7)

The binding rate of RNAP to each unique gene $k_{\text{bind}}^{G\_\text{locusNum:RNAP}}$ is

$$\text{RNAP} + G_{\text{mRNA}}^{\text{locusNum}} \rightarrow \text{RNAP} : G_{\text{mRNA}}^{\text{locusNum}} + \sigma, \quad k_{\text{bind}}^{G_{\text{locusNum:RNAP}}} = S_{\text{promoter}}^{\text{locusNum}} k_{\text{bind}}^{\text{mRNAgenes:RNAP}} \tag{22}$$

The elongation speed of RNAP on mRNA-coding genes is assumed to scale linearly with the promoter strength $S_{\text{promoter}}^{\text{locusNum}}$ as $S_{\text{promoter}}^{\text{locusNum}} \times 20$ nt/s, where 85 nt/s is assigned as the upper limit. Then, the polymerization of mRNA proceeds as follows.

$$\text{RNAP} : G_{\text{mRNA}}^{\text{locusNum}} \rightarrow \text{mRNA} + \text{RNAP} + G_{\text{mRNA}}^{\text{locusNum}}, \quad v_{\text{polymerization}}^{\text{transcription,mRNA}} \tag{23}$$

$$v_{\text{polymerization}}^{\text{transcription,mRNA}} = \frac{\min(85\,\text{nt/s}, S_{\text{promoter}}^{\text{locusNum}} \times 20\,\text{nt/s})}{\frac{K_{D1,\text{NTP}_1} K_{D2,\text{NTP}_2}}{[\text{NTP}_1][\text{NTP}_2]} + \sum_i \frac{n_i K_{Di,\text{NTP}_i}}{[\text{NTP}_i]} + L_{\text{mRNA}} - 1} N_{\text{RNAP}:G_{\text{mRNA}}^{\text{locusNum}}} \tag{24}$$

Next, we describe the parameterization for translation, which was conducted in a similar fashion as that for transcription. The reported values for the mean translation initiation time per mRNA vary widely, with a minimum of 1 second and a median of 15 seconds.(8) In our model, we assume a translation initiation frequency of 60 events per minute. The number of ribosomes in slow-growing *E. coli* has been experimentally measured as 6800.(9) Then the binding rate between mRNA and ribosome $k_{\text{bind}}^{\text{mRNA:ribosome}}$ was calculated as

$$k_{\text{bind}}^{\text{mRNA:ribosome}} = \frac{\frac{60}{60} \times V_{E.coli} \times N_A}{6800} = 8.9 \times 10^4\,\text{M}^{-1}\text{s}^{-1} \tag{25}$$

The translation speed was set to 12 aa/s.(10) The synthesis of proteins happens via the binding and elongation reactions.

$$\text{Ribosome} + \text{mRNA} \rightarrow \text{Ribosome} : \text{mRNA}, \quad k_{\text{bind}}^{\text{mRNA:ribosome}} = 8.9 \times 10^4\,\text{M}^{-1}\text{s}^{-1} \tag{26}$$

$$\text{Ribosome} : \text{mRNA} \rightarrow \text{Ribosome} + \text{mRNA} + \text{protein}, \quad v_{\text{polymerization}}^{\text{translation}} \tag{27}$$

$$v_{\text{polymerization}}^{\text{translation}} = \frac{12\,\text{aa/s}}{\frac{K_{D1,\text{tRNA}_1} K_{D2,\text{tRNA}_2}}{[\text{tRNA:aa}_1][\text{tRNA:aa}_2]} + \sum_i \frac{n_i K_{Di,\text{tRNA}_i}}{[\text{tRNA:aa}_i]} + L_{\text{protein}} - 1} \tag{28}$$

Finally, we describe the parameterization of mRNA degradation. The binding rate between mRNA and the degradosome $k_{\text{bind}}^{\text{mRNA:degradosome}}$ is estimated based on the observed value of 11 RNase E cleavage events per minute per RNase E (11), and 7800 mRNAs (12) measured in *E. coli*.

$$k_{\text{bind}}^{\text{mRNA:degradosome}} = \frac{\frac{11}{60} \times V_{E.coli} \times N_A}{7800} = 1.4 \times 10^4\,\text{M}^{-1}\text{s}^{-1} \tag{29}$$

The degradation rate of an mRNA polymer chain was considered constant over the length of the chain (4), with the speed taken to be 88 nt/s (13). The product of mRNA degradation is a collection of nucleotide monophosphates. The respective reactions are

$$\text{Degradosome} + \text{mRNA} \rightarrow \text{Degradosome} : \text{mRNA}, \quad k_{\text{bind}}^{\text{mRNA:degradosome}} = 1.4 \times 10^4\,\text{M}^{-1}\text{s}^{-1} \tag{30}$$

$$\text{Degradosome} : \text{mRNA} \rightarrow \text{Degradosome} + \sum_i \text{NMP}_i, \quad v_{\text{degradation}}^{\text{mRNA}} = \frac{88\,\text{nt/s}}{L_{\text{mRNA}}} \tag{31}$$

ATP energizes bond formation in tRNAs and mRNAs and breaks down mRNAs at the expense of one ATP molecule per monomer. Two GTP molecules are required in protein chain elongation for each amino acid, one for the delivery of aa: tRNA to the A site and another for the shift of the ribosome to the next codon.

##### 1.2.3 Transcription of rRNAs

In this model, we approximated the phenomenon of the famous 'Chrismas tree' (14), multiple RNAP reading the rRNA operon at the same time by increasing the rate/affinity of RNAP binding. For transcription of six rRNA-coding genes, higher affinities of RNAP binding (more frequently transcription initiation) were granted to ensure adequate rRNAs transcribed for ribosome biogenesis. The lengths of 5S, 16S, and 23S rRNA are 108, 2913, and 1534, respectively. To represent the multiple RNAPs, we scale the binding rate by nine, six, and two for 23S, 16S, and 5S rRNAs to be $7.92 \times 10^4$, $5.28 \times 10^4$, and $1.76 \times 10^4$ $\mathrm{M}^{-1}\mathrm{s}^{-1}$ respectively in a way that is determined by the length of the rRNA and the assumption that the RNAP needs a spacing interval of 300 to 400 nucleotides. Based on the average waiting times in gene expression processes, the rate-determining step of transcription is RNAP binding with gene. Then we could approximate the two-step reactions as one pesudo first-order reaction, where the speed of RNA synthesis is linear to the RNAP binding rate, thus we roughly imitate the affect of multiple RNAP transcribing rRNAs.

$$\mathrm{RNAP} + G_{\mathrm{rRNA}} \rightarrow \mathrm{RNAP} : G_{\mathrm{rRNA}} + \sigma, \quad k_{\mathrm{bind}}^{\mathrm{rRNA\_genes:RNAP}} \tag{32}$$

The elongation speed of RNAP on rRNA-coding genes is taken as fixed 85 nt/s (6), and the elongation occurs via the following formula.

$$\mathrm{RNAP} : G_{\mathrm{rRNA}} \rightarrow \mathrm{rRNA} + \mathrm{RNAP} + G_{\mathrm{rRNA}}, \quad v_{\mathrm{polymerization}}^{\mathrm{transcription,rRNA}} \tag{33}$$

$$v_{\mathrm{polymerization}}^{\mathrm{transcription,rRNA}} = \frac{85\,\mathrm{nt/s}}{\frac{K_{D1,\mathrm{NTP}_1} K_{D2,\mathrm{NTP}_2}}{[\mathrm{NTP}_1][\mathrm{NTP}_2]} + \sum_i \frac{n_i K_{Di,\mathrm{NTP}_i}}{[\mathrm{NTP}_i]} + L_{\mathrm{rRNA}} - 1} N_{\mathrm{RNAP:G_{rRNA}}} \tag{34}$$

#### 1.3 Communication between Genetic Information Process and Metabolism

The discreteness and stochasticity of chemical kinetics play a role when the number of reactants is significantly low. For the species in genetic information processes and complex formation, most mRNAs have a number of 1 or 2. Based on the initial volume of JCVI-syn3A (200 nm radius, 0.035 fL, 1 particle count equals 50 nM), mRNA concentrations are 50 or 100 nM. Therefore, it is necessary to simulate the kinetics in genetic information processes (GIP) and protein complex assembly with stochastic CME and metabolism with deterministic ODE.

In CME formula 35, the state of the system $\mathbf{x}$ as a vector is the counts of all species. The transition between different states is the fire of single chemical reaction $r$ out of all reactions $R$ with stoichiometry $\mathbf{S_r}$. The probability for the fire of each single chemical reaction in next time step $dt$ is $a_r(\mathbf{x})dt$, where we name $a_r(\mathbf{x})$ propensity of reaction $r$ under system state $\mathbf{x}$.

$$\frac{dP(\mathbf{x},t)}{dt} = \sum_{r}^{R}[-a_r(\mathbf{x})P(\mathbf{x},t) + a_r(\mathbf{x}_\nu - \mathbf{S_r})P(\mathbf{x} - \mathbf{S_r}, t)] \tag{35}$$

To simulate the co-evolution of GIP, complex formation with metabolism, the communication needs to be performed to describe the interactions between these two subsystems. We first discretize the entire simulation length into piecewise communication time steps (hook intervals, $t_H$). During each communication time step, **(a)** a CME simulation of length $t_H$ is performed to describe the kinetics in GIP, **(b)** followed by the communication from CME to ODE by passing the protein counts, the consumption of monomers (dNTP, NTP, and amino acid charged tRNAs), and the recycling of nucleoside monophosphates (NMP) in the degradation of mRNA and recycling of amino acids in the degradation of membrane protein. **(c)** Then a $t_H$ length ODE simulation is performed with the updated concentrations of proteins and metabolites. **(d)** The impacts of metabolism on GIP are two-fold: the abundance of metabolites explicitly in GIP and the concentrations of monomers that affect the rates in the polymerization of gene, RNA, and protein. Here, we used `getReactionCountsView()` and `getReactionRateConstantView(reactionNumber)` methods in Python problem-solving environment (PSE) integrated in LM to access the counts and rates in the computer's memory to update them per communication step, thus applying the impacts.

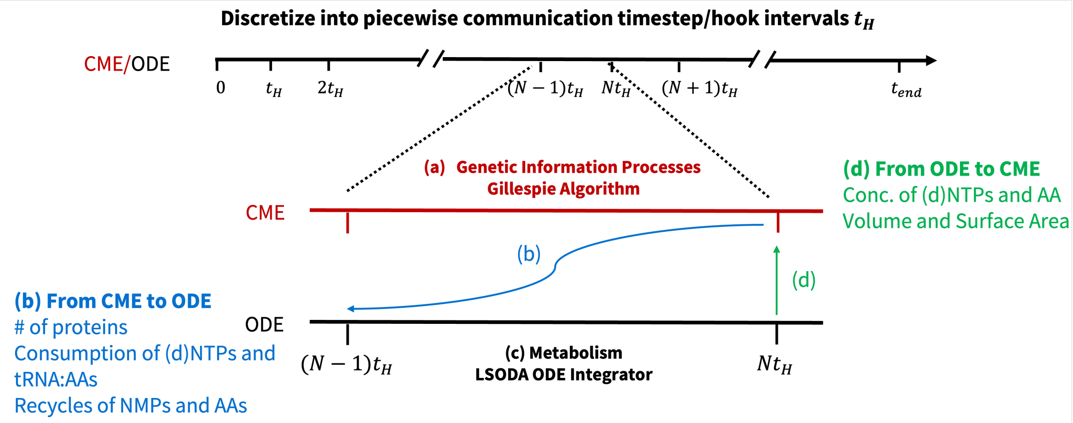


Figure S3: Time-line scheme of communication between genetic information processes in CME and metabolism in ODE

As shown in Figure S4, the CME simulation is executed using Lattice Microbes (LM) with direct Gillespie algorithm. We employ the `hookSimulation` function to interrupt the CME timeline and enable communication with the ODE solver. For the ODE simulation, we use the odecell software developed by the Luthey-Schulten Lab, which maps metabolic reactions to ordinary differential equations and specifies the corresponding kinetic parameters. The resulting ODE system is solved using the `lsoda` algorithm from the SciPy library.

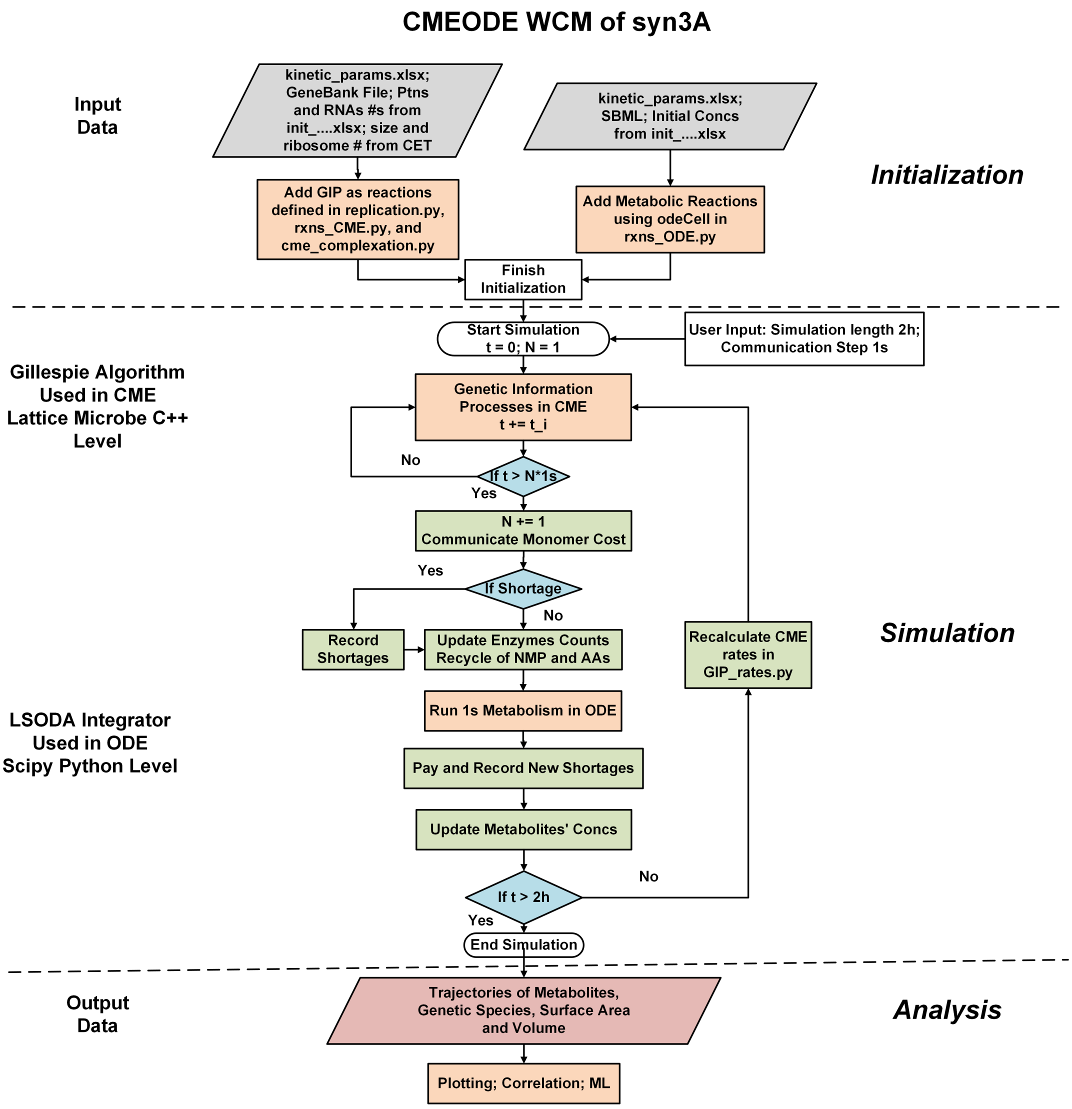


Figure S4: Flowchart of CME/ODE Communication

The CME/ODE simulations can be efficiently parallelized across up to 25 cell replicates or more, with each replicate requiring less than 2GB of RAM. On systems equipped with Intel Xeon Gold 6154 CPUs @ 3.00 GHz or AMD EPYC 7763 "Milan" processors, the parallel simulations typically complete within 6 hours.

### 2 Macromolecular Complexes in Syn3A and Assembly Pathways

#### 2.1 Compositions and Assembly Pathways

Table S2 shows the compositions of macromolecular complexes simulated in this work. The initial count of complete complexes (*Init. Cplx. Cnt*) were determined as described in the Methods. The reasoning of the compositions were discussed in detail in the following subsections. The experimentally measured counts of subunits (*Exp. Ptn Cnt*) were available at File S2, and some of them were discussed in the following subsections. The assembly pathways of the complexes are also summarized in Table S3.

Table S2: Macromolecular Complexes and Their Compositions [a]

| Name | *Init. Cplx. Cnt* | Protein Subunit | *Exp. Ptn Cnt* (Stoich.) | Translocation [b] |
|---|---|---|---|---|
| RNAP | 93 | $\alpha$ (RpoA)/0645 | 187 (2) | - |
| | | $\beta$ (RpoB)/0804 | 529 (1) | - |
| | | $\beta'$(RpoC)/0803 | 546 (1) | - |
| | | $\sigma$ (RpoD)/0407 | 227 (1) | - |
| SMC-ScpAB | 101 | SMC/0415 | 202 (2) | - |
| | | ScpA/0327 | 1 (1) | - |
| | | ScpB/0328 | 31 (2) | - |
| Gyrase | 122 | GyrA/0007 | 298 (2) | - |
| | | GyrB/0006 | 244 (2) | - |
| Topo-IV | 78 | ParC/0453 | 156 (2) | - |
| | | ParE/0452 | 157 (2) | - |
| RNDR | 50 | NrdE/0771 | 382 (2) | - |
| | | NrdI/0772 | 50 (1) | - |
| | | NrdF/0773 | 297 (2) | - |
| Degradosome | 120 | RNaseY (Rny)/0359 | 206 (1) | SRP/SR/Sec |
| | | RNaseJ1 (RnjA)/0600 | 120 (1) | Diffusion |
| | | RNaseJ2 (RnjB)/0257 | 335 (1) | Diffusion |
| | | YhaM/0437 | 517 (1) | Diffusion |
| Sec Translocon | 66 | SecY/0652 | 66 (1) | SRP/SR/Sec |
| | | SecE/0839 | 8 (1) | SRP/SR/Sec |
| | | SecG/0774 | 4 (1) | SRP/SR/Sec |
| | | SecDF/0412 | 215 (1) | SRP/SR/Sec |
| ATP Synthase | 108 | $F_0$ $a$ (AtpB)/0796 | 8 (1) | SRP/SR/Sec |
| | | $F_0$ $c$ (AtpE)/0795 | - (10) | YidC |
| | | $F_0$ $b$ (AtpF)/0794 | 182 (2) | SRP/SR/Sec |
| | | $F_1$ $\delta$ (AtpH)/0793 | 32 (1) | Diffusion |
| | | $F_1$ $\alpha$ (AtpA)/0792 | 327 (3) | Diffusion |
| | | $F_1$ $\gamma$ (AtpG)/0791 | 136 (1) | Diffusion |
| | | $F_1$ $\beta$ (AtpD)/0790 | 347 (3) | Diffusion |
| | | $F_1$ $\epsilon$ (AtpC)/0789 | 108 (1) | Diffusion |
| ECF core [c] | 25 | EcfT/0641 | 25 (1) | SRP/SR/Sec |
| | | EcfA/0642 | 271 (1) | Diffusion |
| | | EcfA/0643 | 0 (1) | Diffusion |
| ribflECF | 10 | FmnP/0877 | 10 (1) | SRP/SR/Sec |
| p5pECF | 10 | EcfS/0345 | 10 (1) | SRP/SR/Sec |
| 5fthfECF | 10 | EcfS/0822 | 10 (1) | SRP/SR/Sec |
| nacECF | 150 | EcfS/0314 | 150 (1) | SRP/SR/Sec |
| CoaECF | 10 | EcfS/0836 | 10 (1) | SRP/SR/Sec |
| KtrCD | 38 | KtrC/0686 | 305 (8) | Diffusion |
| | | KtrD/0685 | 17 (2) | SRP/SR/Sec |
| Fak [d] | 205 | FakA/0420 | 205 (1) | Diffusion |
| | | FakB/0616 | 366 (1) | Diffusion |
| | | FakB/0617 | 205 (1) | Diffusion |

[a] Cytoplasmic macromolecular complexes in the first half; Membrane complexes in the second half; The 3 rRNAs and 51 ribosomal proteins not shown for ribosome; The composition of ABC transporters in Table S6.
[b] Transmembrane proteins are translocated via SRP/SR/Sec or YidC pathways; Peripheral membrane protein diffuse to the peripheral region in Table S9.
[c] Five ECF transporters share the same ECF core module.
[d] Two unique FakB subunits compete to bind with FakA.

Table S3: Assembly Pathways of Macromolecular Complexes [a]

| Name | Reaction | Location [b] | Rate |
|---|---|---|---|
| RNAP | $\alpha + \alpha \rightarrow \alpha_2$ | Cyto | $k^{\text{cyto}}_{\text{assembly}}$ |
| | $\alpha_2 + \beta \rightarrow \alpha_2\beta$ | Cyto | $k^{\text{cyto}}_{\text{assembly}}$ |
| | $\alpha_2\beta + \beta' \rightarrow \text{core}$ | Cyto | $k^{\text{cyto}}_{\text{assembly}}$ |
| | $\alpha_2 + \beta' \rightarrow \alpha_2\beta'$ | Cyto | $k^{\text{cyto}}_{\text{assembly}}$ |
| | $\alpha_2\beta' + \beta \rightarrow \text{core}$ | Cyto | $k^{\text{cyto}}_{\text{assembly}}$ |
| | $\text{core} + \sigma \rightarrow RNAP$ | Cyto | $k^{\text{cyto}}_{\text{assembly}}$ |
| SMC-ScpAB | $SMC + SMC \rightarrow SMC_2$ | Cyto | $k^{\text{cyto}}_{\text{assembly}}$ |
| | $SMC_2 + ScpA \rightarrow SMC_2ScpA$ | Cyto | $k^{\text{cyto}}_{\text{assembly}}$ |
| | $ScpB + ScpB \rightarrow ScpB_2$ | Cyto | $k^{\text{cyto}}_{\text{assembly}}$ |
| | $SMC_2ScpA + ScpB_2 \rightarrow SMC - ScpAScpB$ | Cyto | $k^{\text{cyto}}_{\text{assembly}}$ |
| Gyrase | $GyrA + GyrA \rightarrow GyrA_2$ | Cyto | $k^{\text{cyto}}_{\text{assembly}}$ |
| | $GyrA_2 + GyrB \rightarrow GyrA_2B$ | Cyto | $k^{\text{cyto}}_{\text{assembly}}$ |
| | $GyrA_2B + GyrB \rightarrow Gyrase$ | Cyto | $k^{\text{cyto}}_{\text{assembly}}$ |
| Topo-IV | $ParC + ParC \rightarrow ParC_2$ | Cyto | $k^{\text{cyto}}_{\text{assembly}}$ |
| | $ParC_2 + ParE \rightarrow ParC_2E$ | Cyto | $k^{\text{cyto}}_{\text{assembly}}$ |
| | $ParC_2E + ParE \rightarrow Topo - IV$ | Cyto | $k^{\text{cyto}}_{\text{assembly}}$ |
| RNDR | $nrdE + nrdE \rightarrow nrdE_2$ | Cyto | $k^{\text{cyto}}_{\text{assembly}}$ |
| | $nrdF + nrdF \rightarrow nrdF_2$ | Cyto | $k^{\text{cyto}}_{\text{assembly}}$ |
| | $nrdE_2 + nrdF_2 \rightarrow nrdE_2F_2$ | Cyto | $k^{\text{cyto}}_{\text{assembly}}$ |
| | $nrdE_2F_2 + nrdI \rightarrow RNDR$ | Cyto | $k^{\text{cyto}}_{\text{assembly}}$ |
| Degradosome | $RNase\_J1 + RNase\_J2 \rightarrow RNase\_J1J2$ | PM | $k^{\text{mem}}_{\text{assembly}}$ |
| | $RNase\_Y + RNase\_J1J2 \rightarrow RNase\_YJ1J2$ | PM | $k^{\text{mem}}_{\text{assembly}}$ |
| | $RNase\_YJ1J2 + yhaM \rightarrow Degradosome$ | PM | $k^{\text{mem}}_{\text{assembly}}$ |
| Sec Translocon | $SecY + SecE \rightarrow SecYE$ | TM | $k^{\text{mem}}_{\text{assembly}}$ |
| | $SecYE + SecG \rightarrow SecYEG$ | TM | $k^{\text{mem}}_{\text{assembly}}$ |
| | $SecYEG + SecDF \rightarrow SecYEGDF$ | TM | $k^{\text{mem}}_{\text{assembly}}$ |
| ABCTransporters [c] | TMDs → Channel | TM | $k^{\text{mem}}_{\text{assembly}}$ |
| | Channel + NBDs →→ Channel_NBDs | PM | $k^{\text{mem}}_{\text{assembly}}$ |
| | Channel_NBDs + SBP → ABC Transporter | TM | $k^{\text{mem}}_{\text{assembly}}$ |
| ECF Core module | $EcfT + EcfA \rightarrow EcfTA$ | PM | $k^{\text{mem}}_{\text{assembly}}$ |
| | $EcfTA + EcfA' \rightarrow ECF$ | PM | $k^{\text{mem}}_{\text{assembly}}$ |
| | $EcfT + EcfA' \rightarrow EcfTA'$ | PM | $k^{\text{mem}}_{\text{assembly}}$ |
| | $EcfTA + EcfA \rightarrow ECF$ | PM | $k^{\text{mem}}_{\text{assembly}}$ |
| ECF Transporters | $ECF + S_i \rightarrow S_iECf$ | TM | $k^{\text{mem}}_{\text{assembly}}$ |
| | $S_iECf \rightarrow ECf + S_i$ | TM | $5 \times 10^{-4}$ s$^{-1}$ [d] |
| KtrCD | $KtrD + KtrD \rightarrow KtrD_2$ | TM | $k^{\text{mem}}_{\text{assembly}}$ |
| | $KtrC + KtrC \rightarrow KtrC_2$ | PM | $k^{\text{mem}}_{\text{assembly}}$ |
| | $KtrD_2 + KtrC_2 \rightarrow KtrC_2D_2$ | PM | $k^{\text{mem}}_{\text{assembly}}$ |
| | $KtrC_2D_2 + KtrC_2 \rightarrow KtrC_4D_2$ | PM | $k^{\text{mem}}_{\text{assembly}}$ |
| | $KtrC_2D_2 + KtrC_2 \rightarrow KtrC_6D_2$ | PM | $k^{\text{mem}}_{\text{assembly}}$ |
| | $KtrC_2D_2 + KtrC_2 \rightarrow KtrCD$ | PM | $k^{\text{mem}}_{\text{assembly}}$ |
| Fak | $FakA + FakB/0616 \rightarrow Fak$ | PM | $k^{\text{mem}}_{\text{assembly}}$ |
| | $FakA + FakB/0617 \rightarrow Fak$ | PM | $k^{\text{mem}}_{\text{assembly}}$ |

[a] Assembly pathways of ribosome SSU and LSU in ribo_assembly.xlsx in File S3; pathways of ATP synthase in Table S5.

[b] Cyto: cytoplasm; PM: peripheral membrane; TM: transmembrane.

[c] General assembly scheme of five ABC transporters, where TMD is transmembrane domain, NBD nucleotide binding domain (ATPase), SBP substrate binding protein. The exact reactions depends on the correspondence between protein subunits and domains in TableS6.

[d]: Half-life of 23 min assumed.

#### 2.2 RNA Polymerase

RNA polymerase (RNAP) transcribes the DNA sequence to the RNA sequence with a controlled and organized process. The composition of bacterial RNAP is well conserved, where the RNAP core enzyme $\alpha_2\beta\beta'$ and $\sigma$ factors are essential (Table S2) with the accessory subunits $\delta$, $\epsilon$ and $\omega$ (15).

The $\sigma$ factor transiently associates with the RNAP core enzyme $\alpha_2\beta\beta'$ to recognize the promoter and dissociates once RNAP initiates the processive synthesis of RNA. In the minimal Syn3A cell, only one $\sigma$ factor was found, inherited from its parent organism *Mycoplasma mycoides.* This single $\sigma$ factor is homologous to the $\sigma^{70}$ type factor in *E. coli* or the $\sigma^{A}$ vegetative type of Gram-positive bacteria (16).

The only accessory subunit identified in Syn3A was the $\delta$ subunit, also aligning with *Mycoplasma mycoides.* The $\delta$ subunit was confirmed to enhance transcriptional specificity (17) and the recycling of the RNAP core enzyme (18) at transcription termination.

The assembly of the RNAP core enzyme $\alpha_2\beta\beta'$ is the first step in the transcription cycle and begins with the dimerization of two $\alpha$ subunits and the further incorporation of the $\beta$ and $\beta'$ subunits. $\beta$ and $\beta'$ subunits form the catalytic center of RNA synthesis and provide the binding sites during transcription. The $\sigma$ factor transiently associates with the core enzyme for promoter recognition and dissociates from the core enzyme once transcription elongation starts. After transcription, the intact core enzyme also detaches from the DNA chain (19). See Table S3 for the assembly pathways of RNAP and transcription reactions.

#### 2.3 Ribosome

The bacterial ribosome comprises a small subunit (30S SSU) and a large subunit (50S LSU), each of which is a ribonucleoprotein complex that assembles independently (14; 20). The monocistronic rRNA operons (*rrsA*/0069, *rrlA*/0068, *rrfA*/0067 and *rrsB*/0534, *rrlB*/0533, *rrfB*/0532) encode the 16S, 23S, and 5S rRNAs, respectively. 16S rRNA with 20 unique kinds of SSU ribosomal proteins assemble in the SSU; 23S, 5S, and 31 kinds of LSU ribosomal proteins assemble in the LSU.

In this model, three rRNAs were expressed separately; the same was true for ribosomal proteins. The SSU assembly pathways were derived from the previous study in *E. coli* (21; 22) by reducing the number of assembly intermediates from 145 to 19 to maintain the smallest number of complete pathways with the highest fluxes. The binding rates of each SSU ribosomal protein remain the same as in the original study. The fastest rate of of SSU assembly was $3.1 \times 10^7$ $M^{-1}$ $s^{-1}$ for ribosomal protein S6 as the third binder to associate, and lowest rate was $2 \times 10^4$ $M^{-1}$ $s^{-1}$ for the last binder, S12. Same as the previous study, ribosomal proteins S2 and S21 are not included due to the lack of kinetic data (21).

For LSU, we sampled one linear assembly pathway on the rerouted Nierhaus assembly map in Figure 3(B) (20) where the order of protein binding was revisited by characterization of *in vivo* assembly intermediates using quantitative mass spectrometry and cryo-EM. Based on the strength of protein binding cooperativities measured by Nierhaus, we categorized binding as fast ($10^6$ $M^{-1}$ $s^{-1}$) or slow ($10^4$ $M^{-1}$ $s^{-1}$). SSUs and LSUs will finally associate to form intact ribosomes at a fast rate that will bind to mRNA for the translation process.

Assembly factors function significantly to assist the ribosome biogenesis. A scanning of the annotated Syn3A's proteome (File S2-Tab 1) that updated after the computational analysis leveraging AlphaFold (23) identified seven rRNA processing proteins, ten rRNA modification proteins, one rRNA helicase, seven ribosome-dependent GTPase, and five ribosome dedicated chaperons or assembly factors in Table S4. Most of these rRNA processing and modification enzymes (24), rRNA helicase (25), GTPase (26), and assembly factors/dedicated chaperons (27) are essential for viability in Syn3A, except for most enzymes in rRNA modification.

During bacterial ribosome biogenesis, three broad classes of accessory proteins act in concert to mature the rRNA scaffold. Processing and modification enzymes (endonucleases, exonucleases, methyltransferases, pseudouridine synthases) carry out the irreversible chemical steps—cleaving precursor transcripts and installing stabilizing nucleotide modifications that define the final rRNA sequence and chemistry. RNA helicases provide the mechanical force to rearrange rRNA secondary and tertiary structures in an ATP-dependent but reversible manner, resolving kinetic traps and exposing otherwise buried sites for cleavage, modification, or ribosomal-protein binding. Assembly GTPases and dedicated chaperones (or assembly factors) serve as energy-driven checkpoints and escort proteins, sensing when local rRNA domains have achieved the correct fold and coordinating the ordered incorporation of ribosomal proteins and late processing factors. Together these activities ensure that the 16S, 23S, and 5S rRNAs are precisely processed, chemically matured, and folded into the stable architecture required for formation of translationally competent ribosomal subunits.

Table S4: Ribosome Biogenesis Factors: 31 kinds of proteins in five categories

| Type | Protein | NCBI Description | Ess.[a]/Abundance |
|---|---|---|---|
| rRNA Processing [b] | RnmV/0003 | Ribonuclease M5 | Q/65 |
| | YqgF/0215 | Pre-16S rRNA nuclease | E/49 |
| | RnjB/0257 | RNase J family beta-CASP ribonuclease | E/335 |
| | Rny/0359 | Ribonuclease Y | E/206 |
| | YbeY/0402 | rRNA maturation RNase | E/64 |
| | RnC/0418 | Ribonuclease III | E/59 |
| | RnpA/0909 | Ribonuclease P protein component | E/6 |
| rRNA modification | KsgA/0004 | 16S rRNA (adenine(1518)-N(6)/adenine(1519)-N(6))-dimethyltransferase | N/40 |
| | RsmD/0202 | 16S rRNA (guanine(966)-N(2))-methyltransferase | Q/125 |
| | RlmH/0361 | 23S rRNA (pseudouridine(1915)-N3)-methyltransferase | N/123 |
| | RlmFO/0434 | 23S rRNA (uridine(1939)-m5)-methyltransferase | N/70 |
| | SpoU/0448 | Uncharacterized rRNA methyltransferase | E/317 |
| | RsmL/0504 | 16S rRNA (cytidine(1402)-2'-O)-methyltransferase | N/38 |
| | RluD/0517 | Uncharacterized RNA pseudouridine synthase | N/97 |
| | MraW/0524 | 16S rRNA (cytosine(1402)-N(4))-methyltransferase | N/33 |
| | RlmB/0838 | 23S rRNA (guanosine(2251)-2'-O)-methyltransferase | N/135 |
| | RsmG/0874 | 16S rRNA (guanine(527)-N(7))-methyltransferase | N/131 |
| rRNA helicase | CshB/0410 | Degradosome RNA helicase | Q/120 |
| GTPase | YihA/0247 | Ribosome biogenesis GTP-binding protein | E/63 |
| | Eng/0263 | Ribosome small subunit-dependent GTPase A | Q/- |
| | EngA/0348 | Ribosome biogenesis GTPase | E/34 |
| | RbgA/0366 | L16-binding dependent 50S subunit-maturation GTPase | E/146 |
| | ObgE/0377 | Ribosome GTPase | E/150 |
| | Era/0403 | Ribosome GTPase | E/46 |
| | YchF/0872 | Ribosome Binding Factor ATPase | Q/168 |
| Assembly factors | RbfA/0289 | Ribosome-binding factor A | Q/194 |
| | RimP/0301 | Ribosome assembly cofactor | Q/140 |
| | RimM/0363 | 16S rRNA processing protein | E/5 |
| | YsxB/0500 | Maturation protease for ribosomal protein L27 | E/42 |
| | YlbN/0527 | 23S rRNA biogenesis regulators (DUF177/YceD) | N/331 |

[a] Ess.: Essentiality, E: essential, Q: quasiessential, N: nonessential [b] Overlap with ribonucleases in degradosome.

#### 2.4 Chromosome Dynamics Related Complexes

The minimal chromosome organizing complexes encoded by Syn3A's genome include structural maintenance of the chromosome complex SMC-ScpAB and two type II topoisomerases, DNA gyrase and topoisomerase IV. Their functions in Syn3A have previously been explored using a continuum bead-on-string polymer model (28).

SMC-ScpAB complex has a composition of $SMC_2ScpAScpB_2$ shown in Table S2 (29). We inferred that dimerization of the SMC subunits could occur separately from ScpA and ScpB (30) based on the homodimer structure of the hinge domain of the SMC subunit in *Pyrococcus furiosus*(PDB ID: 4RSJ). Another structure of homodimer with ScpA from *Bacillus subtilis* (PDB ID: 5XG3) indicates that first ScpA binds to the SMC dimer and then the ScpB subunit. See the SI Table S3 for assembly reactions.

DNA-gyrase (gyrase) and topoisomerase-IV (TopoIV) are of compositions $GyrA_2GyrB_2$ and $ParC_2ParE_2$, where the GyrA and ParC subunits contain DNA binding and catalytical domain, and ATP binding resides in the GyrB and ParE subunits (31). Here, we begin the assembly of both heterotetramers from the dimerization of GyrA and ParC, supported by the structures of their homodimer

(32; 33), and the complex with DNA(34). Then another subunit, GyrB or ParE, could bind to form the holocomplex. See SI Table S3 for assembly reactions.

Exonuclease VII (ExoVII), the ubiquitous bacterial nuclease, was recently reported to comprise a highly elongated $XseA_4 \cdot XseB_{24}$ holocomplex, and observed in its crystal structure (PDB ID: 8TXR) (35). ExoVII in bacteria functions to degrade single-stranded DNA in both 5' to 3' and 3' to 5' directions, playing a critical role in DNA repair and recombination. Due to the novelty of such structure and its function not considered in the current WCM, the assembly of ExoVII was not considered in this study.

Stoichiometries among the SMC-ScpAB subunits were reported in (29), and of gyrase, Topo-IV, and ExoVII supported by the PDB entry 6RKS (36), 2Q2E (32), and 8TXR (35), respectively.

#### 2.5 RNDR

Ribonucleoside-diphosphate reductase (RNDR, encoded by *nrdE*/0771, *nrdI*/0772, and *nrdF*/0773) (1) converts ribonucleoside diphosphates to their deoxy version that could be used as monomers in gene replication, and is essential enzymes for bacterial (37). Class 1b RNDR $\alpha_2\beta_2$ consists of two homodimeric subunits (37), where $\alpha$ subunit as catalytic center encoded by *nrdE*/0771 and $\beta$ as radical-generating subunit by *nrdF*/0773. The protein NrdI/0772 is a flavodoxin that mediates a two-electron reduction of $\beta_2$ (38). Here, we model the holocomplex as $\alpha_2\beta_2$NrdI that is capable of catalyzing the reaction in nucleotide metabolism. The initial count of the holocomplex was set to 50, the minimum possible count that is the count of subunits NrdI. The assembly pathway of KtrCD was assumed to be in Table S3.

#### 2.6 Degradosome

Primary mRNA transcripts in bacteria are protected at their 5' end by a triphosphate group inherited from transcription. Since there is no existence of RNA pyrophosphohydrolase (e.g. RppH in *B. subtilis*) that could convert 5' triphosphate into 5' monophosphate in Syn3A, mRNA degradation should only be initiated by the endoribonuclease cut by single-strand-specific RNase Y or double-strand-specific Rnase III (39). The downstream product generated with the 5' monophosphate extremity can be attacked and degraded by 5'-3' exoribonuclease Rnase J1 and J2. The 3' hydroxyl of the upstream product is degraded by the 3'-5' exoribonuclease RNase R and YhaM.

The scaffold and endoribonuclease RNase Y was observed to form a dimer *in vitro* by a long parallel N-terminus coil structure that was verified by structural analysis of AlphaFold predictions (40). However, the effect of dimerization on the binding and cutting of mRNA was not evaluated. Because of the independence of the catalytic domain, we assume that the RNase Y monomer could initiate mRNA degradation. The interactions between the components of the mRNA degradosome are weak and transient in the Gram-positive model organism, *B. subtilis*, supported by the fact that RNase Y was localized on the membrane through its TMD, while RNase J1 and J2 and the helicase were distributed in the peripheral region and the glycolytic enzymes were uniform throughout the cell (41).

Here, we modeled the "degradosome", a functional holocomplex of mRNA degradation as one copy of RNase Y as scaffold and endoribonuclease, one copy of RNase J1/J2 heterodimer as the 5' to 3' exoribonucleases (39), and one copy of YhaM as the 3' to 5' exoribonucleases. The initial count of this particle was set to 120, the count of RNase J1. We assumed that two glycolytic enzymes of Eno/0213 and PfkA/0220 as the compositions of degradosome without explicitly simulating their association for two reasons: first, the abundances of two enzymes are 998, and 551 respectively and will not limit the assembly; second, these glycolytic enzymes were reported to uniformly distributed in the whole cell of *B. subtilis* (41), and the exact function of those interacting with degradosomes was still largely unknown. Thus, we assumed that two enzymes function as needed in both glycolysis in metabolism and degradation of mRNAs.

Consequently, mRNA degradation was simulated by binding an mRNA to a "degradosome" to form a complex and digested to final product mononucleotides (NMP) at a speed of 88 nucleotides per second (4). See Table S3 for assembly reactions of degradosome and mRNA degradation reactions.

#### 2.7 Sec Translocon

Sec translocon is a conserved protein complex in eukaryotes and prokaryotes for the translocation of transmembrane and secretary proteins. Based on the study of *in vitro* translation and assembly of SecY/0652, SecE/0839, and SecG/0774 subunits (42), it is likely that SecY first binds to SecE to form the SecYE complex and then incorporates SecG.

SecDF/0412 could accelerate the insertion of secretory proteins in Ribosome/SecYEG/SecA complex (43). The SecYEGDFyajC complex was identified in the immunoprecipitation study, indicating its stability (44). Here, we assume that SecDF could bind to SecYEG through probable interactions between SecF and SecY (45), and that the holocomplex of SecYEGDF is active in translocations of transmembrane proteins and secretory proteins.

#### 2.8 ATP Synthase

$F_1F_0$-ATP synthase couples reversible ATP synthesis with an electrochemical proton concentration gradient across cytoplasm and extracellular space in Gram-positive Syn3A. Protons flow back into the cytoplasm through the membrane-embedded $F_0$ sector of ATP synthase, driving the rotation of the peripheral $F_1$ sector and powering ATP synthesis. Or reversely, ATP is hydrolyzed to actively pump protons out to extracellular space when there is a high intracellular concentration of ATP and protons, thus maintaining the proton motive force (PMF) that is crucial for proper nutrient uptake and translocation of transmembrane proteins and secretory proteins.

9 ATP synthase genes are organized in a single operon read from gene 0797 to gene 0789 (Table S2), and are all identified as essential (1). Since the atomic structure of ATP synthase in Syn3A or its parent organism *Mycoplasma mycoides* is still not available, we borrow the typical structural composition and possible assembly pathways from model bacterium, *E. coli* that was intensively reviewed by Ruhle and Leister (46). The $F_1$ sector consists of four distinct subunits $\alpha$, $\beta$, $\gamma$, $\delta$ and $\epsilon$ with stoichiometry 3, 3, 1, 1, and 1, which were assumed to be translated into the cytoplasm and diffuse to the peripheral region. The membrane-embedded $F_0$ sector comprises three subunits $a$, $b$, and $c$ with stoichiometry 1, 2 and 10. The transmembrane chaperon AtpI assists in the formation of the $c_{10}$ ring and is not incorporated into the final holocomplex. This chaperoning process has not been explicitly simulated in this work with no experimental evidence (47; 48).

Although previous research indicates that all protein subunits are translated from a single polycistronic mRNA (49) and differential synthesis is primarily regulated by translation initiation (50; 51), we still assume independent transcription, translation and translocation of the 9 subunits due to the absence of organism-specific experimental data from Syn3A. The translocation of the transmembrane protein subunits of ATP synthase occur through the SRP/SR/Sec pathway, except for subunit $F_0$ c that had been verified to be translocated without SRP and SecYEG translocon (52) to support the sole YidC translocation mechanism. The peripheral $F_1$ subunits are translated in the cytoplasm and diffuse into the region of the peripheral membrane for assembly.

In the simulation, the stoichiometric balance between the subunits is approximated by adjusting the promoter strength that defined in section Genetic Information Process, a prefactor that affects the transcription rate and was defined as proportional to the abundances of proteins (53; 4). The initial count of intact ATP synthase was given as the number of $F_1$ subunit $\epsilon$ 108, considering the good stoichiometric ratio among the $F_1$ sector subunits $\alpha$, $\beta$, $\gamma$, and $\epsilon$ of 3:3:1:1. In contrast, possibly due to the deficit of mass spectrometry in capturing proteomics, the counts of transmembrane subunit a is only 8, and no subunit c detected.

The assembly of ATP synthase occurs in a modular pattern (detailed reactions and kinetic rates listed in Table S5 ). Upon translocation, the $F_0$ sector transmembrane subunit $c$ could assembly independently from other subunits to form the $c_{10}$-ring. In gel electrophoresis experiments, the intermediate oligomer $c$ subunits, $c_i$ ($i = 2 \ldots 9$), were not labeled with some bands between, which remained unrecognized (48). Here, we used a most favorable oligomerization scheme for the formation of $c_{10}$ rings, where the incorporation of a single C subunit was allowed to occur by one in the oligomer intermediates and also binding between the oligomers. To test the effect of binding between oligomers on $c_{10}$-ring formation, we compared two binding rates, 2.5 $\times 10^{-4}$ $\mu m^2 s^{-1}$ or 2.5 $\times 10^{-3}$ $\mu m^2 s^{-1}$. A lower binding rate of 2.5 $\times 10^{-4}$ $\mu m^2 s^{-1}$ was used considering the slower diffusion of the oligomers compared to the sole subunit $c$.

The $F_0$ sector subunit $b$ will dimerize to $b_2$, which will bind and stabilize the subunit $a$ (54) by forming a heterocomplex $ab_2$. The module of the $F_1$ peripheral sector $\alpha_3\beta_3\gamma\epsilon$ is formed by trimerization of the heterodimer $\alpha\beta$ and subsequent binding of $\gamma\epsilon$. This module could bind to the $c_{10}$-ring by the interaction between the central stalk $\gamma\epsilon$ and the ring to form $\alpha_3\beta_3\gamma\epsilon c_{10}$. Taking into account the large displacement in binding between the $\alpha_3\beta_3\gamma\epsilon$ and the $c_{10}$-ring, we tested two parameters 2.5 $\times 10^{-4}$ $\mu m^2 s^{-1}$ or 2.5 $\times 10^{-3}$ $\mu m^2 s^{-1}$ to see their influence on ATP synthase assembly.

Finally, the transmembrane module $ab_2$ and the peripheral module $\alpha_3\beta_3\gamma\epsilon c_{10}$ need to be joined by the binding of the $F_1$ $\delta$ subunit, a short 181 AAs long peripheral membrane protein. The intermediate of assembly $ab_2c_{10}\delta$ was confirmed to exist when these four subunits were expressed in *in vitro* reconstitution experiments. In contrast, the $\delta$ subunit was not incorporated into the subcomplex $\alpha_3\beta_3\gamma\epsilon c_{10}$ when only the subunit $b$ was eliminated (55). This indicated that the $\delta$ subunit has a stronger affinity to $ab_2$. Here, we tested two possible discrimination methods for binding of the $\delta$ subunit to $ab_2$ or $\alpha_3\beta_3\gamma\epsilon c_{10}$. The first was using 2.5 $\times 10^{-4}$ $\mu$m$^2$s$^{-1}$ or 2.5 $\times 10^{-3}$ $\mu$m$^2$s$^{-1}$, and the binding between $\delta$ and $ab_2$ remained 2.5 $\times 10^{-3}$ $\mu$m$^2$s$^{-1}$. The second was that reverse dissociation reactions were applied to $\alpha_3\beta_3\gamma\delta\epsilon c_{10}$ with a higher rate of $10^{-2}$ s$^{-1}$ and to $\delta ab_2$ with a lower rate of $10^{-4}$ s$^{-1}$. $10^{-4}$ s$^{-1}$ corresponds to the half-life of 1.9 hours ($\frac{\ln(2)}{10^{-4}} \times \frac{1}{3600}$), and $10^{-4}$ s$^{-1}$ corresponds to the half-life of 1.1 min.

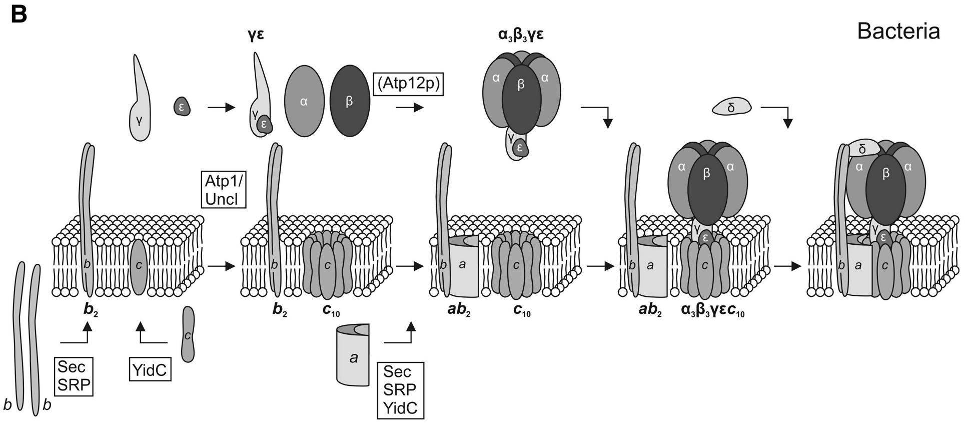


Figure S5: Assembly of ATP synthase in Bacteria. Reprinted with permission from Ruhle, T, Leister, D (2015) Assembly of F1F0 ATP synthase. *Biochimica et Biophysica Acta (BBA) - Bioenergetics* **2015**, **1847**:849-860. Copyright 2015/ELSEVIER

Table S5: Assembly Pathways of ATP synthase (46)

| Reaction | Type | Location | Rate |
|---|---|---|---|
| Translocation of subunits a, b, c | Co-/Post-translational Translocation | Cyto and TM | - |
| Translation of subunits $\alpha, \beta, \gamma, \delta$ and $\epsilon$ | Translation | Cyto | - |
| $\alpha + \beta \rightarrow \alpha_1\beta_1$ | Diffusion & Binding | C | $k^{\text{cyto}}_{\text{assembly}}$ |
| $\alpha_1\beta_1 + \alpha_1\beta_1 \rightarrow \alpha_2\beta_2$ | Diffusion & Binding | Cyto | $k^{\text{cyto}}_{\text{assembly}}$ |
| $\alpha_2\beta_2 + \alpha_1\beta_1 \rightarrow \alpha_3\beta_3$ | Diffusion & Binding | Cyto | $k^{\text{cyto}}_{\text{assembly}}$ |
| $\gamma + \epsilon \rightarrow \gamma\epsilon$ | Diffusion & Binding | Cyto | $k^{\text{cyto}}_{\text{assembly}}$ |
| $\alpha_3\beta_3 + \gamma\epsilon \rightarrow \alpha_3\beta_3\gamma\epsilon$ | Diffusion & Binding | Cyto | $k^{\text{cyto}}_{\text{assembly}}$ |
| $c_i + c \rightarrow c_{i+1}, i = 1, \ldots 9$ | Diffusion & Binding | TM | $k^{\text{mem}}_{\text{assembly}}$ |
| $c_i + c_j \rightarrow c_{i+j}, i = 2, ..8, i + j \leq 10$ | Diffusion & Binding | TM | $2.5 \times 10^{-4}$ $\mu$m$^2$s$^{-1}$ [a] or $k^{\text{mem}}_{\text{assembly}}$ |
| $b + b \rightarrow b_2$ | Diffusion & Binding | TM | $k^{\text{mem}}_{\text{assembly}}$ |
| $a + b_2 \rightarrow ab_2$ | Diffusion & Binding | TM | $k^{\text{mem}}_{\text{assembly}}$ |
| $\alpha_3\beta_3\gamma\epsilon + c_{10} \rightarrow \alpha_3\beta_3\gamma\epsilon c_{10}$ | Diffusion & Docking | PM | $2.5 \times 10^{-4}$ $\mu$m$^2$s$^{-1}$ [b] or $k^{\text{mem}}_{\text{assembly}}$ |
| $\delta + \alpha_3\beta_3\gamma\epsilon c_{10} \rightarrow \alpha_3\beta_3\gamma\delta\epsilon c_{10}$ | Diffusion & Docking | PM | $2.5 \times 10^{-4}$ $\mu$m$^2$s$^{-1}$ [c] or $k^{\text{mem}}_{\text{assembly}}$ |
| $\alpha_3\beta_3\gamma\delta\epsilon c_{10} \rightarrow \alpha_3\beta_3\gamma\epsilon c_{10} + \delta$ [d] | Disassociation | PM | $10^{-2}$ s$^{-1}$ |
| $\alpha_3\beta_3\gamma\delta\epsilon c_{10} + ab_2 \rightarrow ATPsynthase$ | Diffusion & Binding | PM and M | $k^{\text{mem}}_{\text{assembly}}$ |
| $\delta + ab_2 \rightarrow \delta ab_2$ | Diffusion & Docking | PM | $k^{\text{mem}}_{\text{assembly}}$ |
| $\delta ab_2 \rightarrow ab_2 + \delta$ [d] | Disassociation | PM | $10^{-4}$ s$^{-1}$ |
| $\delta ab_2 + \alpha_3\beta_3\gamma\epsilon c_{10} \rightarrow ATPsynthase$ | Diffusion & Binding | PM and M | $k^{\text{mem}}_{\text{assembly}}$ |

[a] Weaken affinity among subunit C oligomers

[b] Weaken affinity between $C_{10}$ ring with peripheral stalk $\alpha_3\beta_3\gamma\epsilon$

[c] Weaken affinity between $\delta$ subunit with $\alpha_3\beta_3\gamma\epsilon c_{10}$

[d] Add reverse disassociation reactions of $\alpha_3\beta_3\gamma\delta\epsilon c_{10}$ and $\delta ab_2$

#### 2.9 ABC Importers

During the construction of essential metabolism (1; 4), ABC transporters in Syn3A are identified to import crucial molecular precursors for metabolism: rnsBACD for nucleoside, Pst system for phosphate, PotABC and ThiBPQ for spermine and thiamine diphosphate, and Opp for oligopeptide and amino acid. ABC importers in Gram-positive organisms have a typical composition of two high-conserved peripheral nucleotide binding domains (NBD), two transmembrane domains (TMD) as permeases, and a substrate binding protein (SBP) facing extracellular space that delivers the substrates to the TMDs (56).

Although with a common modular architecture, the correspondence between the protein subunits with domains shows the variety of structures. Of the five ABC importers, only Opp has a one-to-one relationship. The gene $rnsA$/0010 encodes a heterodimer of NBDs for the nucleoside importer rnsBACD (57). A permease encoded by the spermine ABC transporter permease encoded by $potC$/0195 has 7 TMRs and 500 amino acids outside the membrane, indicating that the SBP fused with a TMD. For the Pst system, the SBP is a transmembrane protein with a TMR embedded, whereas the SBPs of rnsBACD, ThiBPQ, and Opp are all membrane-anchored lipoproteins.

Table S6: ABC Transporters and Their Compositions in Syn3A[a]

| Name | Init. Cplx. Cnt | Exp. Ptn Cnt (Stoich.) | Subunit (Localization) | Domain Correspondence |
|---|---|---|---|---|
| rnsBACD | 145 | 5 (1) | RnsD/0008 (TM) | TMD |
| | | 175 (1) | RnsC/0009 (TM) | TMD |
| | | 145 (1) | RnsA/0010 (PM) | 2NBDs |
| | | 81 (1) | RnsB/0011 (Lipo) | SBP |
| Opp | 241 | 59 (1) | AmiC/0165 (TM) | TMD |
| | | 64 (1) | OppC/0166 (TM) | TMD |
| | | 242 (1) | AmiE/0167 (PM) | NBD |
| | | 241 (1) | AmiF/0168 (PM) | NBD |
| | | 314 (1) | AmiA/0169 (Lipo) | SBP |
| PotABC | 104 | 182 (1) | PotC/0195 (TM) | TMD, SBP |
| | | 19 (1) | PotB/0196 (TM) | TMD |
| | | 208 (1) | PotA/0197 (PM) | 2NBDs |
| Pst system | 123 | 56 (1) | PstS/0425 (TM) | SBP |
| | | 26 (1) | PstA/0426 (TM) | 2TMDs |
| | | 123(2) | PstB/0427 (PM) | 2NBDs |
| ThiBPQ | 20 | 2 (1) | ThiP/0706 (TM) | 2TMDs |
| | | 41 (2) | ThiQ/0707 (PM) | 2NBDs |
| | | 210 (1) | ThiB/0708 (Lipo) | SBP |

[a] Listed subunits of each ABC importer add up to an intact complex.

The assembly pathways of the ABC transporters were assumed to obey the following order. The TMDs first bind to each other to form the permease channel, and the peripheral NBDs bind to the channel to form the functional core. SBP will bind with the core at last, if separated from the TMDs. The underlying reason was that SBP and NBD all need to interact with the permease channel to bind to the holocomplex. The rates of association are $k^{\mathrm{mem}}_{\mathrm{assembly}}$.

#### 2.10 ECF Transporters

Energy-coupling factor (ECF) transporters are specific for micronutrients, such as enzymatic cofactors or their precursors (such as B-type vitamins). Unlike ABC transporters, ECF transporters do not make use of periplasmic or extracellular substrate binding proteins (SBPs) (56) to import substances. Based on existing structures, ECF transporters consist of a membrane permease EcfT/0641 as a scaffold, a dimer of ATPase EcfA/0642 and EcfA/0643, and a substrate binding protein EcfS, S-components. The first three subunits form an ECF module (core) that is highly conserved, and different S-components import specific substrates(58). The assembly of ECF module presumably happens via EcfT, binding to either one of the ATPase or another one, under the argument that the EcfT subunit is the scaffold. The initial count of the ECF module was set to 25, the same as the experimentally measured count of EcfT subunit.

ECF-type ABC transporters are classified into three groups: group I, group II, and solitary. Group II transporters are modular, meaning that different S-components can compete to interact and bind to the same ECF module (59). Five ECF transporters (S-components) for nicotinate (Conjugate base of Vitamin B3), pyridoxal (Vitamin B6), 5-formyltetrahydrofolate (derivative of folate, Vitamin B9), coenzyme A, and riboflavin (Vitamin B2) were characterized in the reconstruction of the essential metabolism (1). Four out of five ECF transporters are classified as Group II with unknown CoaECF based on the summary by Slotboom *et al.* (59). So, reversible binding and unbinding reactions of the ECF module with different S components were included in the simulation. The unbinding rate was $5 \times 10^{-4}$ $s^{-1}$ correponding to the half-life of 23 min. The initial counts of each ECF transporter were assumed to be the count of their S-Components. See Table S3 for detailed assembly reactions of ECF transporters.

#### 2.11 KtrCD, and Fak

Ktr ion transporters are crucial for potassium uptake, and the complex consists of two components: transmembrane permeases (KtrB or KtrD) and peripheral regulatory protein (KtrA or KtrC) (60). In the protein BLAST test against model Gram-positive organism, *Bacillus subtilis*, we found that the transmembrane protein encoded by gene *natA*/0685 has 95% query cover with KtrD (NCBI Reference Sequence P_134155297.1), and also gene *trkA*/0686 92% query cover with KtrC (NCBI Reference Sequence WP_041335737.1). Based on the BLAST test, we assumed that gene *trkD*/0685 and *trkA*/0686 encodes KtrCD complex for potassium uptake. Compared with the KtrAB complex, KtrCD is less studied with no structure deposited. Here, it was assumed that the KtrCD complex has the same compositions of $KtrC_8D_2$ as $KtrA_8B_2$, dimeric transmembrane proteins with the peripheral octamer (homodimer tetramer) as shown in PDB entry 4J7C (60; 61). The initial count of the complex KtrCD was set to 38, as the count of the peripheral subunit KtrD 305 over its stoichiometry 8. The assembly pathway of KtrCD was assumed to be in Table S3.

The fatty acid kinase (Fak) system activates the phosphorylation of fatty acids to acyl phosphate by ATP at the initial stage of lipid metabolism, termed reaction FaKr. A Fak complex consists of an ATP-binding kinase, FakA, and a fatty acid-binding protein, FakB, as confirmed by gel filtration and separation (62). Thus, the formation of a Fak complex is a straightforward single-step binding event between a FakA protein and a FakB protein. The *fakA*/0420 gene encodes the kinase FakA, while two FakB proteins, FakB/0616 and FakB/0617, are present in Syn3A (1), presumably favoring different lipid substrates, as observed in another Gram-positive bacterium, *Staphylococcus aureus* (63). However, in current lipid metabolism, the saturation states and double-bond positions of the fatty acids were not distinguished. For simplicity, it was assumed that half of the FakA subunits bind to FakB/0616, named Fak1 and the other half to FakB/0617, named Fak2. In the FaKr reaction, the abundances of enzymes Fak are the sum of Fak1 and Fak2.

#### 2.12 Representative Subunits of Macromolecular Complexes

Table S7 summarized the representative subunits used in the control simulation.

Table S7: Representative subunits of Macromolecular Complexes in Control Simulations [a]

| Name | *Init. Cplx. Cnt* | Representative Protein Subunit [b] | *Exp. Ptn Cnt* (Stoich.) | Translocation |
|---|---|---|---|---|
| RNAP | 93 | $\alpha$ (RpoA)/0645 | 187 (2) | - |
| Sec Translocon | 66 | SecY/0652 | 66 (1) | SRP/SR/Sec |
| Degradosome | 120 | RNaseY (Rny)/0359 | 206 (1) | SRP/SR/Sec |
| | | RNaseJ1 (RnjA)/0600 | 120 (1) | Diffusion |
| RNDR | 50 | NrdE/0771 | 382 (2) | - |
| | | NrdI/0772 | 50 (1) | - |
| | | NrdF/0773 | 297 (2) | - |
| ATP Synthase | 108 | $F_1$ $\epsilon$ (AtpC)/0789 | 108 (1) | Diffusion |
| ECF core [c] | 25 | EcfT/0641 | 25 (1) | SRP/SR/Sec |
| | | EcfA/0642 | 271 (1) | Diffusion |
| | | EcfA/0643 | 0 (1) | Diffusion |
| ribflECF | 10 | FmnP/0877 | 10 (1) | SRP/SR/Sec |
| p5pECF | 10 | EcfS/0345 | 10 (1) | SRP/SR/Sec |
| 5fthfECF | 10 | EcfS/0822 | 10 (1) | SRP/SR/Sec |
| nacECF [d] | 150 | EcfS/0314 | 150 (1) | SRP/SR/Sec |
| CoaECF | 10 | EcfS/0836 | 10 (1) | SRP/SR/Sec |
| KtrCD | 38 | KtrC/0686 | 305 (8) | Diffusion |
| Fak | 205 | FakA/0420 | 205 (1) | Diffusion |
| | | FakB/0616 | 366 (1) | Diffusion |
| | | FakB/0617 | 205 (1) | Diffusion |
| rnsBACD | 145 | RnsC/0009 | 175 (1) | SRP/SR/Sec |
| | | RnsA/0010 | 145 (1) | Diffusion |
| Opp | 241 | AmiF/0168 | 241 (1) | SRP/SR/Sec |
| PotABC | 104 | PotC/0195 | 182 (1) | SRP/SR/Sec |
| | | PotB/0196 | 19 (1) | SRP/SR/Sec |
| | | PotA/0197 | 208 (2) | Diffusion |
| Pst System | 123 | PstS/0425 | 56 (1) | SRP/SR/Sec |
| | | PstA/0426 | 26 (1) | SRP/SR/Sec |
| | | PstB/0427 | 123 (2) | Diffusion |
| ThiBPQ | 20 | ThiP/0706 | 2 (1) | SRP/SR/Sec |
| | | ThiQ/0707 | 41 (2) | SRP/SR/Sec |
| | | ThiB/0708 | 210 (1) | SecA/Sec |

[a] Macromolecular complexes in gene expression in the first half; complexes in enzymatic and transport reactions in the second half.

[b] The abundances of macromolecular complexes equal to the minimal of counts over stoichiometries of all representative protein subunits with exceptions discussed in the main text.

[c] Five ECF transporters all need the ECF core module to import nutrients. Four ECF transporters except nacECF have four representative subunits, three in ECF core module, and one S-component.

[d] The only representative subunit assumed for nacECF is EcfS/0314 due to its high experimental protein abundance.

#### 3 Translocation Network in Syn3A

##### 3.1 Localization of 455 Proteins

84 transmembrane proteins were predicted to have 1 (single-pass) to 14 (multi-pass) transmembrane regions (TMRs). Typically, the TMRs are composed of hydrophobic amino acid residues and the N-terminus, C-terminus. The loops between the TMRs are hydrophilic. Depending on the predicted topologies, 12 tail-anchored proteins (TA) were identified, which contain single-pass and two-pass TMPs. Of 12 proteins, only 0839, 0774, and 0795 were characterized, respectively, as the preprotein translocase subunit, the preprotein translocase subunit and the ATP synthase $F_0$ subunit c, while others were not characterized. Experimentally, ATP synthase subunit c was verified to be translocated without the SRP and Sec translocon to support the sole YidC translocation pathway (52). See Table S8 for the full list of TA proteins and their TMRs. 15 secretory proteins were identified that have signal peptides, and 13 of 15 are lipoproteins. Among the 13 lipoproteins, 3 of them are substrate binding proteins in ABC transporters (0011 for nucleosides, 0169 for oligopeptides/amino acids, 0708 for thiamine), while the other 11 have unclear functions.

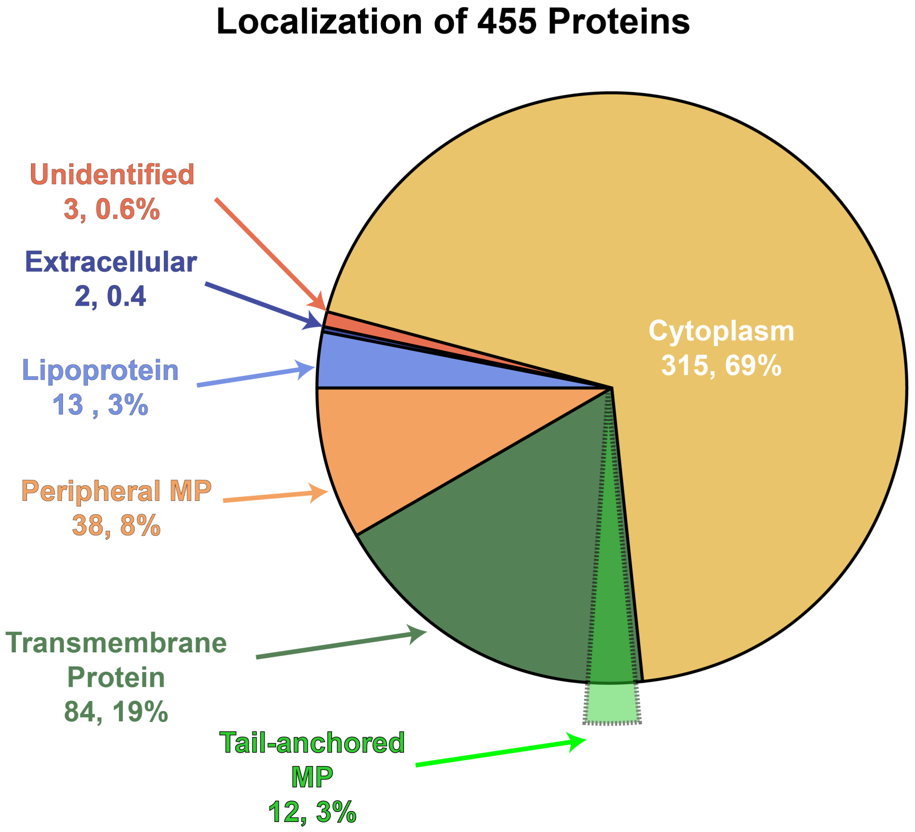


Figure S6: Localization of the entire proteome, consisting of total 455 kinds of proteins. Out of 84 transmembrane proteins, 12 are tail-anchored (TA) membrane proteins. TA membrane proteins are translocated by YidC. Extracellular proteins and lipoproteins are both secretory proteins.

###### 3.1.1 Two Extracellular Proteins, and Three Proteins with Unidentified Localization

Proteins 0503 and 0505 are secreted to the extracellular media, supported by the prediction from DeepTMHMM that the rest of 0503 and 0505's polypeptide chains are outside the membrane. However, three outlier proteins (with locusnum 0033, 0392, and 0623) were still predicted to be secreted into the extracellular space by DeepTMHMM but no signal peptide predicted by Signal P. Without further verification, we classified them as unidentified and assumed that they were cytoplasmic proteins.

##### 3.1.2 Tail-anchored (TA) Transmembrane Proteins

YidC as the insertase could function separately on transmembrane proteins with only short hydrophilic translocated segments (64). Tail-anchored (TA) proteins with the targeting TMR located less than 65 amino acids from the C terminus are typically posttranslational translocated by YidC since the short amino acids distance left upon SRP binding with ribosome leaving no time for the cargo transfer to Sec translocon (65). Depending on the predicted topologies, the following TA proteins are identified, containing both single-pass and two-pass TMPs.

Table S8: Tail-anchored Proteins Translocated via Sole YidC Pathway

| Single-Pass | | | Two-Pass | | |
|---|---|---|---|---|---|
| LocusNum | Length | TMR [a] | LocusNum | Length | TMR |
| 0116 | 146 | 83-100 | 0774 | 94 | 20-29, 69-89 |
| 0235 | 60 | 37-57 | 0778 | 83 | 15-33, 61-81 |
| 0839 | 107 | 76-97 | 0795 | 101 | 35-60, 73-99 |
| 0346 | 231 | 210-230 | 0830 | 86 | 15-35, 60-80 |
| 0264 | 371 | 350-370 | 0852 | 76 | 8-28, 46-67 |
| 0317 | 81 | 13-34 | | | |
| 0379 | 92 | 29-38 | | | |

[a] TMR: Transmembrane region, shown as the indices of amino acids from N-terminus to C-terminus

Out of 12 proteins, only 0839, 0774, and 0795 were characterized respectively, as preprotein translocase subunit, preprotein translocase subunit, and ATP synthase F0 subunit c, while others uncharacterized. Experimentally, ATP synthase subunit c had been verified to be translocated without SRP and SecYEG translocon to support the sole YidC translocation mechanism (52).

#### 3.2 Translation and Translocation Reactions

We summarize the localization and translocation behaviors of the entire proteome in Table S9. Secretory proteins are targeted to and translocated to the outer leaflet of the membrane via SecA/Sec Translocon, among which lipoproteins are anchored to and diffuse on the 2D membrane. Most transmembrane proteins are targeted by SRP/SR/Sec Translocon pathway, and inserted into the membrane during translation. Single pass tail anchored transmembrane proteins and some short two pass transmembrane proteins could be targeted post-translationally, and inserted via sole YidC or even without assistance (64). Proteins without signal peptides or transmembrane regions are translated in the cytoplasm. They could be further categorized into peripheral membrane proteins and cytoplasmic proteins. Peripheral membrane proteins could diffuse to the peripheral membrane region, and bind to the transmembrane proteins and/or membrane via various interactions thus functioning there. Proteins in cytoplasm are translated and then function in the cytoplasm.

Table S9: Proteins Subunits' Localizations, Translocation Pathway and Diffusive Behavior[a]

| Localization | Number | Translocation Pathway | Diffusive Behavior |
|---|---|---|---|
| Cytoplasm | 315 | - | Diffusion in cytoplasm |
| Peripheral Membrane | 38 | Diffusion | (De)Attach to and Slide along membrane; |
| Transmembrane | 84 | SRP/SR/Sec or YidC | 2D Diffusion within Membrane |
| Lipoprotein | 13 | SecA/Sec | 2D Diffusion within Membrane |
| Extracellular | 2 | SecA/Sec | Secreted into and Diffusion in extracellular space |
| Unidentified | 3 | - | - |
| Total | 455 | - | - |

[a] Lipoproteins and extracellular proteins are secretory proteins

Based on the discussion in the Methods, we summarize the reactions in the translation and translocation networks in TableS10

Table S10: Translation and Translocation Networks in Syn3A [a]

| Pathway | Reaction | Location | Rate |
|---|---|---|---|
| Ribo binding | $Ribosome + mRNA \rightarrow RIBO\_mRNA_{locusNum}$ | Cyto | $8.9 \times 10^4$ $M^{-1}s^{-1}$ |
| SRP/SR/Sec | $RIBO\_mRNA_{locusNum} + SRP \rightarrow SRP\_RIBO\_mRNA_{locusNum}$ | Cyto | $10^5$ $M^{-1}s^{-1}$ |
| | $SRP\_RIBO\_mRNA_{locusNum} + SR \rightarrow SR\_SRP\_RIBO\_mRNA_{locusNum}$ | PM | $10^7$ $M^{-1}s^{-1}$ (66) |
| | $SR\_SRP\_RIBO\_mRNA_{locusNum} + Sec \rightarrow Sec\_SR\_SRP\_RIBO\_mRNA_{locusNum}$ | PM | 2.5 $\times 10^{-3}$ $\mu m^2 s^{-1}$ |
| | $Sec\_SR\_SRP\_RIBO\_mRNA_{locusNum} \rightarrow Sec\_RIBO\_mRNA_{locusNum} + SRP + SR$ | PM | 0.95 $s^{-1}$ (67) |
| | $Sec\_RIBO\_mRNA_{locusNum} \rightarrow P_{locusNum} + Ribosome + mRNA + Sec$ | PM | Hofmeyr $s^{-1}$ |
| YidC | $RIBO\_mRNA_{locusNum} \rightarrow CP_{locusNum} + Ribosome + mRNA$ | Cyto | Hofmeyr $s^{-1}$ |
| | $CP_{locusNum} + YidC \rightarrow YidC\_CP_{locusNum}$ | PM | $10^5$ $M^{-1}s^{-1}$ |
| | $YidC\_CP_{locusNum} \rightarrow YidC + P_{locusNum}$ | TM | 1 $s^{-1}$ |
| SecA/Sec | $RIBO\_mRNA_{locusNum} + SecA \rightarrow SecA\_RIBO\_mRNA_{locusNum}$ | Cyto | $10^5$ $M^{-1}s^{-1}$ |
| | $SecA\_RIBO\_mRNA_{locusNum} + Sec \rightarrow Sec\_SecA\_RIBO\_mRNA_{locusNum}$ | PM | $10^5$ $M^{-1}s^{-1}$ |
| | $Sec\_SecA\_RIBO\_mRNA_{locusNum} \rightarrow Pre\_P_{locusNum} + Ribosome + mRNA + Sec + SecA$ | TM | Hofmeyr $s^{-1}$ |
| PMP Diffusion | $RIBO\_mRNA_{locusNum} \rightarrow Ribosome + CP_{locusNum} + mRNA$ | Cyto | Hofmeyr $s^{-1}$ |
| | $CP_{locusNum} \rightarrow P_{locusNum}$ | Cyto to PM | 10 $s^{-1}$ |
| | $P_{locusNum} \rightarrow CP_{locusNum}$ | PM to Cyto | 10 $s^{-1}$ |
| Failed Target [b] | $RIBO\_mRNA_{locusNum} \rightarrow RIBO\_mRNA_{locusNum}\_Long$ | Cyto | Hofmeyr $s^{-1}$ |
| | $RIBO\_mRNA_{locusNum}\_Long \rightarrow Ribosome + mRNA + CP_{locusNum}$ | Cyto | Hofmeyr $s^{-1}$ |
| | $CP_{locusNum} + FtsH \rightarrow FtsH\_CP_{locusNum}$ | PM | $10^5$ $M^{-1}s^{-1}$ |
| | $FtsH\_CP_{locusNum} \rightarrow FtsH + \sum aas$ | PM | $\approx$ 0.22 $min^{-1}$ (68) |
| Lipo Modification | $Pre\_P_{locusNum} + Lgt \rightarrow Lgt\_Pre\_P_{locusNum}$ | TM | 2.5 $\times 10^{-3}$ $\mu m^2 s^{-1}$ |
| | $Lgt\_Pre\_P_{locusNum} \rightarrow Lgt + Pro\_P_{locusNum}$ | TM | 0.8 $min^{-1}$ (69) |
| | $Pro\_P_{locusNum} + Lsp \rightarrow Lsp\_Pro\_P_{locusNum}$ | TM | 2.5 $\times 10^{-3}$ $\mu m^2 s^{-1}$ |
| | $Lsp\_Pro\_P_{locusNum} \rightarrow Lsp + Modified\_P\_locusNum$ | TM | 2 $min^{-1}$ |

[a] Different translocation pathways apply to different substrate in TableS9. Transmembrane proteins are translocated cotranslationally by SRP/SR/Sec pathway or posttranslationlly by YidC pathway. Secretory proteins (lipoproteins and extracellular proteins) are translocated cotranslationally by SecA/Sec pathway. Peripheral membrane proteins (PMP) diffuse to peripheral membrane region. No translocation occur for cytoplasmic proteins.

[b] Cotranslational translocations of transmembrane proteins, lipoproteins are subject to failure.

Table S10 showed the reactions for the cotranslational translocation of transmembrane proteins. The lower part is the edge case where the active ribosome could not bind with SRP in time, thus leading to the failure of translocation, and protein degradation happened. The rate of $RIBO\_mRNA$ to $RIBO\_mRNA\_Long$ also obeys Hofmeyr's rate form, yet only the first 100 amino acids' segment of the protein were included; from $RIBO\_mRNA\_Long$ to $RIBO\_mRNA + CP_{locusNum}$ included the rest polypeptide chain.

The following Table S11 further showed the correspondence between the simulated reactions with biological process.

Table S11: Biological Correspondence of Sec/SRP Reactions

| Reaction | Biological Processes |
|---|---|
| $Ribosome + mRNA \rightarrow RIBO\_mRNA$ | Binding and Initiation of Translation |
| $RIBO\_mRNA + SRP \rightarrow SRP\_RIBO\_mRNA$ | Signal Recognition |
| $SRP\_RIBO\_mRNA + SR \rightarrow SR\_SRP\_RIBO\_mRNA$ | Signal Reception via peripheral SR and SRP/SR forming closed conformation |
| $SR\_SRP\_RIBO\_mRNA_{locusNum} + Sec \rightarrow Sec\_SR\_SRP\_RIBO\_mRNA_{locusNum}$ | Formation of Quaternary Complex |
| $Sec\_SR\_SRP\_RIBO\_mRNA \rightarrow Sec\_RIBO\_mRNA + SRP + SR$ | GTPase Hydrolysis & Handover of Cargo & Recycle SRP and SR |
| $Sec\_RIBO\_mRNA \rightarrow Protein + Ribosome + mRNA + Sec$ | Elongation and Translocation the nascent chain into Membrane |

Sole YidC pathway can translocate TA proteins listed in Table S8.

Translocation of secretory proteins happens cotranslationally. After inserted into the membrane, two modifier will bind with lipoproteins sequentially for their maturation. $K_M$ between lgt and one synthetic signal peptide was reported to be 0.02 mM with specific activity 0.014864 $\mu$mol/min/mg, equivalent to $k_{cat}$ 0.8 min$^{-1}$ (69). $K_M$ for lspA with proliprotein was 6 $\mu$M (70) with no kinetics in the cleavage step. On the basis of low $K_M$s, we assumed that the binding step was diffusion controlled, and that the modification steps were rate-limiting. The speed of signal sequence cleavage was estimated to satisfy double the number of mature lipoproteins, which was approximately 2200 calculated from proteomics. The $k_{cat}$ was roughly 2/min or 0.033/s. The involvement of PG was neglected because of its huge abundance ($2 \times 10^4$) compared to lipoprotein.

Similar to SRP/SR/Sec pathway for transmembrane proteins, we also consider the failed targeting of secretory proteins but with upper limit of 150 amino acid long nascent polypeptide chain.

Peripheral membrane proteins are translated without any targeting since no signal sequence emerging during translation. The exchange rate of peripheral membrane protein from cytoplasm to peripheral region was set to 10 s$^{-1}$, and from peripheral region to cytoplasm 10 s$^{-1}$. Two arguments to support these two values: first the equilibrium partition of peripheral membrane protein in the cytoplasm and peripheral region was assumed to be 1:1 considering the space of sphere 180 nm and peripheral shell of 20 nm had a ratio of 2.6 to 1, and since the affinity of peripheral membrane protein with the membrane, we assume peripheral membrane protein were equally partitioned; second, the characteristic diffusion time of a protein with diffusion coefficient 1$\mu$m$^2$s$^{-1}$ to the membrane inside a sphere of 200 nm was $\frac{L^2}{6D} = \frac{1}{150}s$, which was fast. Thus, we use a fast exchange rate of 1 s$^{-1}$ to depict the kinetics.

SecA and SRP are recruited to the membrane via the interaction with ribosome, SecYEG, and ribosome, SR, SecYEG respectively, and their translation and cycling between cytoplasm and peripheral membrane are simulated with the reactions in Table S10.

### 4 Decreased Protein Assembly Rate Increases Unassembled Fraction

We use a dimerization reaction of two subunits on the membrane, with transcription, translation (translocation omitted), and mRNA degradation reactions in gene expression to derive an analytical formula on the unassebled fraction of synthesized protein subunit at the end of the cell cycle. We assume two protein subunits have the same expression strength and no initial surplus of free subunits. For the simplicity, gene copy number was fixed to 1 and no cell growth and volume increase was included.

We approximate the transcription and degradation from two-step reactions in our CME simulation to one-step first order reaction since for both processes, the binding step is the rate-determining step as shown in Table S12. RNAP binding to gene takes roughly 300 s, and polymerization of RNA chain takes less than 20 s. mRNA binding to degradosome takes 14 s, and digestion of mRNA chain takes 4 s.

In the following reactions, $k_{bind}$s are the binding rates between RNAP/gene, mRNA/ribosome, and mRNA/degradosome, and $N_{Ava}$s are the free/available counts of RNAP, ribosome, and degradosome.

Table S12: Characteristic Time Used in Transcription, Translation and mRNA degradation. Model Protein is 350 AAs long with initial count of 180.

| Process | Reaction Step | Kinetic Parameter | Value | Available Cnt | Avg. Time/s |
|---|---|---|---|---|---|
| Transcription | RNAP w. Gene | $k_{bind}^{gene:RNAP}$ | $2.1 \times 10^3$ M$^{-1}$ s$^{-1}$ | RNAP 33 | 295 |
| | Elongation | $k_{cat}^{transcription}$ | 20 nt/s | - | 54 |
| Translation | mRNA w. Ribosome | $k_{bind}^{mRNA:Ribo}$ | $8.9 \times 10^4$ M$^{-1}$ s$^{-1}$ | Ribo 125 | 2 |
| | Elongation | $k_{cat}^{translation}$ | 12 AA/s | - | 29 |
| Degradation | mRNA w. Degradosome | $k_{bind}^{mRNA:Deg}$ | $1.4 \times 10^4$ M$^{-1}$ s$^{-1}$ | Deg 102 | 14 |
| | Chain Digestion | $k_{cat}^{degradation}$ | 88 nt/s | - | 12 |

$$G_1 \to G_1 + R_1, \quad v_{trsc,1} = \frac{N_{G_1} N_{Ava,RNAP} k_{bind}^{gene:RNAP}}{N_A V_{cell}} \tag{36}$$

$$R_1 + Ribosome \to RIBO\_mRNA_1, \quad v_{trans,1}^{bind} = \frac{N_{R_1} N_{Ava,Ribo} k_{bind}^{mRNA:Ribo}}{N_A V_{cell}} \tag{37}$$

$$RIBO\_mRNA_1 \to Ribosome + R_1 + P_1, \quad v_{trans,1}^{elongation} = \frac{N_{RIBO\_mRNA_1} 12aa/s}{L_{R_1}} \tag{38}$$

$$R_1 \to \emptyset \quad v_{degra,1} = \frac{N_{R_1} N_{Ava,degra} k_{bind}^{mRNA:Deg}}{N_A V_{cell}} \tag{39}$$

$$G_2 \to G_2 + R_2, \quad v_{trsc,2} = \frac{N_{G_2} N_{Ava,RNAP} k_{bind}^{gene:RNAP}}{N_A V_{cell}} \tag{40}$$

$$R_2 + Ribosome \to RIBO\_mRNA_2, \quad v_{trans,2}^{bind} = \frac{N_{R_2} N_{Ava,Ribo} k_{bind}^{mRNA:Ribo}}{N_A V_{cell}} \tag{41}$$

$$RIBO\_mRNA_2 \to Ribosome + R_2 + P_2, \quad v_{trans,2}^{elongation} = \frac{N_{RIBO\_mRNA_2} 12aa/s}{L_{R_2}} \tag{42}$$

$$R_2 \to \emptyset \quad v_{degra,2} = \frac{N_{R_2} N_{Ava,degra} k_{bind}^{mRNA:Deg}}{N_A V_{cell}} \tag{43}$$

$$P_1 + P_2 \to P_1 P_2, \quad v_{assembly} = \frac{N_{P_1} N_{P_2} k_{assembly}}{N_A SA_{cell}} \tag{44}$$

At steady-state, the mRNA generation and consumption fluxes equal:

$$v_{trsc,1} + v_{trans,1}^{elongation} - v_{trans,1}^{bind} - v_{degra,1} = 0 \quad (45)$$
$$v_{trsc,2} + v_{trans,2}^{elongation} - v_{trans,2}^{bind} - v_{degra,2} = 0 \quad (46)$$

Also, the active ribosome bind and unbind fluxes equal:

$$v_{trans,1}^{elongation} - v_{trans,1}^{bind} = 0 \quad (47)$$
$$v_{trans,2}^{elongation} - v_{trans,2}^{bind} = 0 \quad (48)$$

Then, the abundances of mRNAs at steady state are as follows, which is determined by the numbers of available RNAP, degradosome, and the binding rate with them.

$$R_1 = \frac{N_{G_1} N_{Ava,RNAP} k_{bind}^{gene:RNAP}}{N_{Ava,degra} k_{bind}^{mRNA:Deg}} \quad (49)$$
$$R_2 = \frac{N_{G_2} N_{Ava,RNAP} k_{bind}^{gene:RNAP}}{N_{Ava,degra} k_{bind}^{mRNA:Deg}} \quad (50)$$

Further, the abundances of active ribosomes are calculated as:

$$RIBO\_mRNA_1 = R_1 \frac{N_{Ava,Ribo} k_{bind}^{mRNA:Ribo}}{N_A V_{cell}} \frac{L_{R_1}}{12aa/s} \quad (51)$$
$$RIBO\_mRNA_2 = R_2 \frac{N_{Ava,Ribo} k_{bind}^{mRNA:Ribo}}{N_A V_{cell}} \frac{L_{R_2}}{12aa/s} \quad (52)$$
$$(53)$$

The flux of protein synthesis $v_{ptn}^{synthesis}$ is

$$v_{P_1}^{synthesis} = \frac{N_{RIBO\_mRNA_1} 12aa/s}{L_{R_1}} \quad (54)$$
$$= R_1 \frac{N_{Ava,Ribo} k_{bind}^{mRNA:Ribo}}{N_A V_{cell}} \quad (55)$$
$$= \frac{1}{N_A V_{cell}} \frac{N_{G_1} N_{Ava,RNAP} N_{Ava,Ribo}}{N_{Ava,Degra}} \frac{k_{bind}^{gene:RNAP} k_{bind}^{mRNA:Ribo}}{k_{bind}^{mRNA:Deg}} \quad (56)$$
$$v_{P_2}^{synthesis} = \frac{1}{N_A V_{cell}} \frac{N_{G_2} N_{Ava,RNAP} N_{Ava,Ribo}}{N_{Ava,Degra}} \frac{k_{bind}^{gene:RNAP} k_{bind}^{mRNA:Ribo}}{k_{bind}^{mRNA:Deg}} \quad (57)$$
$$(58)$$

The protein synthesis and protein assembly will reach steady-state:

$$\frac{dP_1}{dt} = v_{P_1}^{synthesis} - v_{assembly} = 0 \quad (59)$$
$$\frac{dP_2}{dt} = v_{P_2}^{synthesis} - v_{assembly} = 0 \quad (60)$$
$$(61)$$

then the abundance of proteins at steady-state will be:

$$[N_{P_1} N_{P_2}]_{SS} = \frac{SA_{cell}}{V_{cell}} \frac{N_{G_{1,2}} N_{Ava,RNAP} N_{Ava,Ribo}}{N_{Ava,Degra}} \frac{k_{bind}^{gene:RNAP} k_{bind}^{mRNA:Ribo}}{k_{bind}^{mRNA:Deg} k_{assembly}} \quad (62)$$
$$N_{P_1,SS} = \sqrt{\frac{SA_{cell}}{V_{cell}} \frac{N_{G_{1,2}} N_{Ava,RNAP} N_{Ava,Ribo}}{N_{Ava,Degra}} \frac{k_{bind}^{gene:RNAP} k_{bind}^{mRNA:Ribo}}{k_{bind}^{mRNA:Deg} k_{assembly}}} \quad (63)$$
$$N_{P_1,SS} = N_{P_2,SS} \quad (64)$$

The unassembled fraction of synthesized protein defined as the count not incorporated into the complex $N_{P_1,SS}$ at the end of the cycle over the number of synthesized over the cell cycle:

$$\phi = \frac{\text{Not Assembled}}{\text{Total Synthesized}} \tag{65}$$

$$= \frac{N_{P_1,SS}}{v_{P_1}^{synthesis} t_{cycle}} \tag{66}$$

$$= \frac{N_A\sqrt{V_{cell}SA_{cell}}}{t_{cycle}} \sqrt{\frac{N_{Ava,Degra}}{N_{G_{1,2}} N_{Ava,RNAP} N_{Ava,Ribo}} \frac{k_{bind}^{mRNA:Deg}}{k_{bind}^{gene:RNAP} k_{bind}^{mRNA:Ribo} k_{assembly}}} \tag{67}$$

Here, we assume initial count of proteins as 180, the average proteomics number; $t_{cycle}$ = 6000 s; $V_{cell}$ = 3.35E-17 L as initial value; $SA_{cell}$=5.02E+5 nm$^2$ as initial value. The unassembled fraction $\phi$ is proportional to the reverse of the root of assembly rate $k_{assembly}$. The synthesized count of protein was 166, and the unassembled count of protein was 2, 7, and 23 for assembly rate of 2.5 $\times 10^{-5}$, 2.5 $\times 10^{-4}$, and 2.5 $\times 10^{-3}$ $\mu$m$^{-2}$s$^{-1}$ with unassembled fraction of 1.3%, 4.4%, and 13.3%.

### 5 Supporting Figures

##### Statistics in genetic information processes

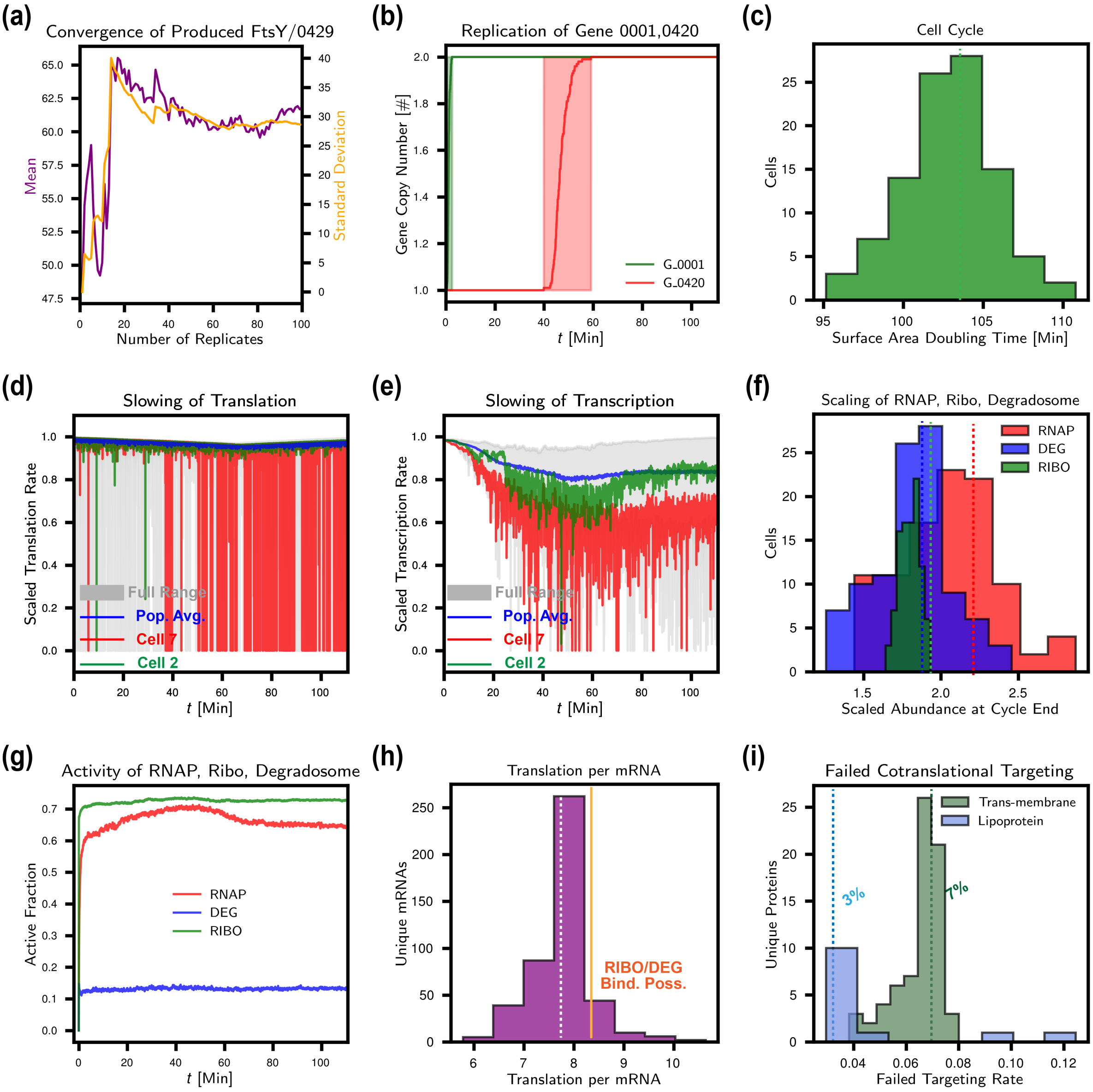


Figure S7: SI Figures regarding genetic information processes. (a) Convergence of synthesized protein FtsY/0429 at 90 min. Same for protein SecA/0095 (not shown) (b) Early replication of Gene 0001 near the *Ori* and late replication of gene 0420 near the *Ter*. (b) Distribution of replication initiation time. (c) Distribution of the cell cycle, i.e. surface area doubling times with median of 102 min. (d) Slowing down of translation due to the transient and temporary charged tRNAs shortage at the temporal resolution of 1 second. The population averaged translation kinetics remained roughly constant. Cell 7's translation rate was frequently zero at the late stage of the cell cycle due to the shortage of certain charged tRNAs. (e) Slowing down of transcription due to the low concentration of UTP. (f) Distribution of scaled RNAP, ribosome, and degradosome at the end of cell cycle with median of 2.09, 1.84, and 1.80. The dotted lines show the median of distribution. (g) Active ratios of RNAP, ribosome and degradosome over the entire cell cycle. The shoulder of RNAP around 40 min came from the faster doubling of the entire chromosomes. (h) Distribution of translation per mRNA for 455 unique mRNAs with median 782 marked by purple dotted line. 8.2 is calculated from the theoretical equations. (i) Failure rate of targeting 72 transmembrane proteins and 13 lipoproteins with median of 7% and 3%. Two outlier in lipoportein 0605 and 0851 is because the use of stricter threshold of the lengths of nascent polypeptide of 51 and 30 AAs, half of their protein lengths because they are shorter than 150.

#### Balance of Amino Acids

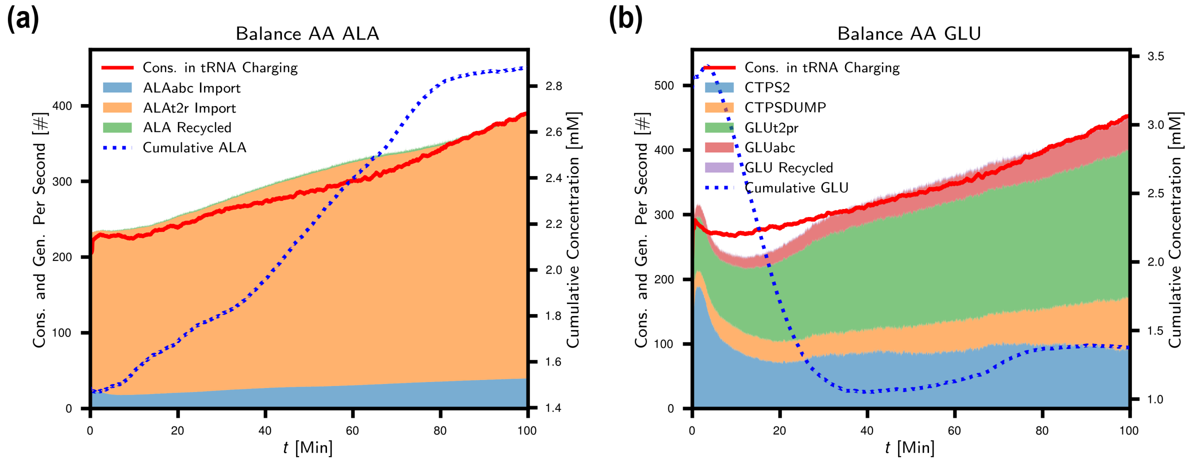


Figure S8: (a) Balance of Amino Acid, Alanine. ALAabc import was the active import by Opp ABC transporter, and ALAt2r the passive import by permeases, 0876 and 0878. The passive import was the dominant contributor to amino acid import. Recycled Alanine from membrane proteins that were failed to target to the membrane was plotted at the top, minor compared to import counts. (b) Balance of Amino Acid, Glutamate. Active import of GLU by GLUabc reaction, the same Opp ABC transporter. Passive by GLUt2pr reaction by permease 0886. Glutamate was also involved in nucleotide and co-factor metabolism, except from tRNA charging and import. The recycled GLU was minor.

**Machine learning on metabolomics to identify three phenotypes in simulation with complex assembly**

More D-Glucose 6-phosphate (G6P) and D-Fructose 6-phosphate (F6P) as early glycolytic metabolites clearly separated cell population 3 from the other two populations before 10 minutes. Correspondingly, population 3 was relatively short in downstream products, such as ATP and phosphoribosyl pyrophosphate (PRPP), and had less intracellular potassium ($K^+$) and calcium ($Ca^{2+}$) that actively imported by membrane ATPase transporters. Population 1 and 2 had more subtle differences with smaller standard effect sizes and diverged later around 26 minutes. The differential reactions disclosed higher fluxes in phosphorelay, glycolysis, pentose phosphate pathway, and phosphate ion import (reaction PIabc) in population 2, leading to more accumulation of PRPP, lactate, and slightly more ATP. In contrast, population 1 had more upstream metabolites, including G6P, F6P, and ribose 5-phosphate (R5P), and intermediates, such as adenosine (ADN) and adenosine diphosphate (ADP). The comparison of listed time-dependent metabolite and fluxes are given in File S4.

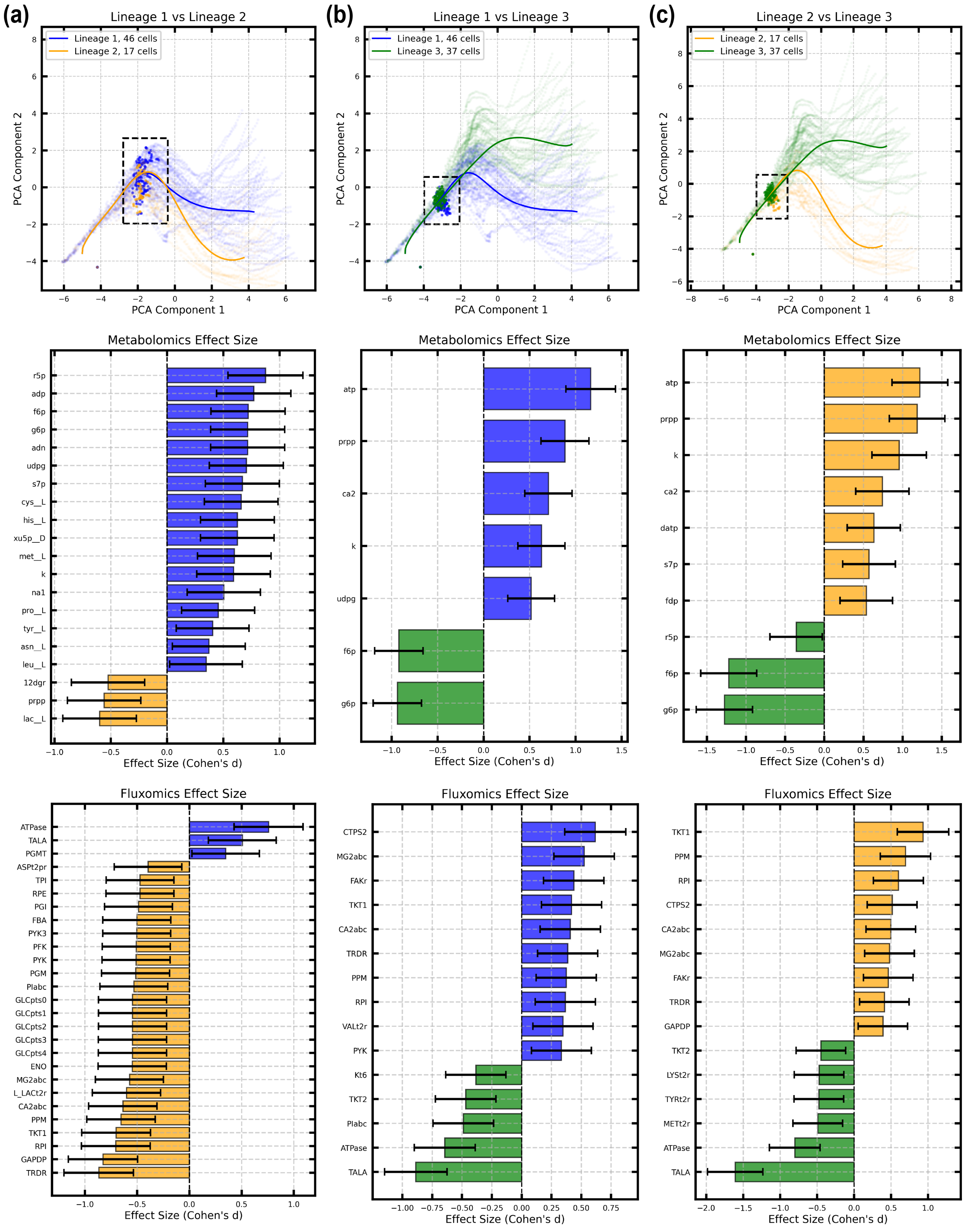


Figure S9: Differential metabolites and reactions for each pair of populations. The top part shows the divergence of the entire metabolome visualized by the first two components in PCA analysis. The dotted brackets highlight the approximate bifurcation point. The effect sizes are shown as the bars that denoted by the color of the lineage, where the reactions have higher fluxes. 95% confidence interval (CI) of the effect size is depicted by the black error bar. These differential reactions are highlighted in reaction map in Figure S1 (a) Cell population 1 and 2. (b) Cell population 1 and 3. (c) Cell population 2 and 3. These differential reactions were marked on the metabolic maps in FigureS1.

### 6 Comparative analysis on metabolomics between with and without complex assembly

#### 6.1 Permutation-Based Two-Sample Test to Tell Statistically Significant Differences

We applied a permutation-based two-sample test using the Maximum Mean Discrepancy (MMD) to evaluate whether metabolite trajectories from the two datasets have the same underlying distribution (71). MMD is a kernel-based statistic that measures the differences between distributions. We denote the metabolite trajectories with assembly and without assembly datasets as $\{X_i\}_{i=1}^{N_1}$ and $\{Y_i\}_{i=1}^{N_2}$, respectively, where

$$X_i = [X_{i,1}, \ldots, X_{i,T_i}], \quad Y_i = [Y_{i,1}, \ldots, Y_{i,T_i}],$$

represent the metabolite concentration of cell $i$ across $T_i$ time points, and $N_1, N_2$ are the numbers of replicates in each dataset. We estimated MMD based on observed trajectories $\{X_i\}_{i=1}^{N_1}$ and $\{Y_j\}_{j=1}^{N_2}$ as

$$\widehat{\mathrm{MMD}}^2_{\mathrm{obs}} = \frac{1}{N_1(N_1-1)} \sum_{i \neq i'} k(X_i, X_{i'}) + \frac{1}{N_2(N_2-1)} \sum_{j \neq j'} k(Y_j, Y_{j'}) - \frac{2}{N_1 N_2} \sum_{i,j} k(X_i, Y_j),$$

where $k(\cdot,\cdot)$ is the kernel defined based on based on multivariate dynamic time warping (DTW) distance (72)

$$k(X, Y) = \exp(-d_{\mathrm{DTW}}(X, Y)/\sigma).$$

DTW measures the minimal cumulative distance between two time series after optimal temporal alignment, which allows the comparison between series with different length. The bandwidth parameter $\sigma$ is selected to be 0.5 in the implementation.

The $p$-value was calculated using a permutation-based approach. Specifically, the dataset labels were permuted $B$ times, and the empirical MMD statistic $\widehat{\mathrm{MMD}}_b$ was computed for each permutation. The $p$-value was then estimated as the proportion of permuted MMD values exceeding the observed statistic:

$$p\text{-value} = \frac{\sum_{b=1}^{B} \mathbb{I}\left(\widehat{\mathrm{MMD}}^2_b \geq \widehat{\mathrm{MMD}}^2_{\mathrm{obs}}\right) + 1}{B+1}$$

where $\mathbb{I}(\cdot)$ is the indicator function. In practice, we select $B = 10000$. If $p$-value is smaller than the significance level of $\alpha = 0.05$, we conclude that the metabolite trajectories from two dataset are significantly different.

This test revealed significant differences in distribution between two datasets ($p = 0.0034$), indicating that metabolite trajectories largely differ depending on whether assembly reactions are included.

#### 6.2 Differential Analysis of Metabolite Trajectories at End of the Cell Cycle

To identify metabolite species and reactions showing differential abundance at the end of the cell cycle, we compared metabolite concentrations and reaction fluxes between the with assembly and without assembly datasets using two-sample $t$-tests. We adjusted the $p$-value using the Benjamini-Hochberg procedure to control false discovery rate. Features with adjusted $p$-values below the significance threshold ($\alpha = 0.05$) were identified as differential between two datasets. The magnitude of differences was quantified by Cohen's $d$, representing the standardized effect size between the two groups.

As shown in Figure **??**, dataset without complex assembly showed higher concentrations of (deoxy)nucleotides and lactate, and higher fluxes of glycolysis and nucleotide metabolism.

#### 6.3 Comparison of Cell Population Distribution of Metabolic Phenotypes Between Datasets

To further examine differences in metabolic phenotypes between datasets, cells in the without assembly dataset were classified into three phenotypes that identified in the dataset with assembly following the procedure described in Section Trajectory Tree Construction and Cell Classification under Methods in the main text. Specifically, latent representations of cells were obtained using the pre-trained autoencoder, and each cell was then classified based on its projected distance to the lineages of trajectory tree. We compared the proportions of different cell types across datasets using a $\chi^2$ test of homogeneity (73). We arranged the observed cell counts into a contingency table, with rows representing datasets and columns representing cell lineages. We denote the observed number of cells of lineage $j$ in the assembly and non-assembly datasets as $N_{1,j}$ and $N_{2,j}$, respectively. The chi-square test statistic is defined as

$$\chi^2 = \sum_{k=1}^{2} \sum_{j=1}^{J} \frac{(N_{k,j} - E_{k,j})^2}{E_{k,j}},$$

where

$$E_{k,j} = \frac{\sum_{j=1}^{J} N_{k,j} \sum_{k=1}^{2} N_{k,j}}{\sum_{k=1}^{2} \sum_{j=1}^{J} N_{k,j}},$$

and $J$ is the number of cell lineages. Under the null hypothesis that there is no difference on subpopulation proportions across datasets, and the test statistic follows a chi-squared distribution with $(J-1)$ degrees of freedom. If the $p$-value is smaller than significance level 0.05, we conclude that subpopulation proportions differ between datasets. The 95% confidence intervals of proportion difference were computed using the Wilson score method (74; 75).

The distribution of populations among three phenotypes changed from 46, 17, and 37 to 47, 40, and 12, respectively, and differed significantly ($\chi^2$ test, $p = 1.635 \times 10^{-5}$). Pairwise proportion tests showed that the dataset without assembly contained more cells in population 2 with higher ATP concentrations ($\Delta$ = 0.234, 95% CI = [0.111, 0.350]), whereas the assembly dataset had more cells in population 3 with lower ATP concentrations ($\Delta$ =-0.249, 95% CI = [-0.344, -0.134]).
